# Genomic signatures of past and present chromosomal instability in the evolution of Barrett’s esophagus to esophageal adenocarcinoma

**DOI:** 10.1101/2021.03.26.437288

**Authors:** Matthew D. Stachler, Chunyang Bao, Richard W. Tourdot, Gregory J. Brunette, Chip Stewart, Lili Sun, Hideo Baba, Masayuki Watanabe, Agoston Agoston, Kunal Jajoo, Jon M. Davison, Katie Nason, Gad Getz, Kenneth K. Wang, Yu Imamura, Robert Odze, Adam J. Bass, Cheng-Zhong Zhang

## Abstract

The progression of precancerous lesions to malignancy is often accompanied by increasing complexity of chromosomal alterations but how these alterations arise is poorly understood. Here we performed haplotype-specific analysis of chromosomal copy-number evolution in the progression of Barrett’s esophagus (BE) to esophageal adenocarcinoma (EAC) on multiregional whole-genome sequencing data of BE with dysplasia and microscopic EAC foci. We identified distinct patterns of copy-number evolution indicating multigenerational chromosomal instability that is initiated by cell division errors but propagated only after p53 loss. While abnormal mitosis, including whole-genome duplication, underlies chromosomal copy-number changes, segmental alterations display signatures of successive breakage-fusion-bridge cycles and chromothripsis of unstable dicentric chromosomes. Our analysis elucidates how multigenerational chromosomal instability generates copy-number variation in BE cells, precipitates complex alterations including DNA amplifications, and promotes their independent clonal expansion and transformation. In particular, we suggest sloping copy-number variation as a signature of ongoing chromosomal instability that precedes copy-number complexity.

*These findings suggest copy-number heterogeneity in advanced cancers originates from chromosomal instability in precancerous cells and such instability may be identified from the presence of sloping copy-number variation in bulk sequencing data.*

---

Large-scale chromosomal rearrangements and copy-number alterations are prevalent in cancer and generally attributed to genomic or chromosomal instability of cancer cells^1–3^. Although much is known about the patterns of genomic rearrangements in fully formed cancers^4, 5^ and the biological mechanisms of genome instability^6–8^, little is understood about what mechanisms are active during cancer evolution and how they generate complex cancer genomes.

Genomic analyses of normal tissues have revealed clonally expanded point mutations but not large structural chromosomal aberrations^9, 10^. Early-stage precancerous lesions also show significantly less genome complexity than late-stage dysplasia^11–15^ or cancer^4, 16, 17^. These observations have led to the prevailing view that most chromosomal rearrangements arise late during cancer progression in an episodic manner^18, 19^, in contrast to the gradual accumulation of short sequence variants (single-nucleotide substitutions or short insertions/deletions)^20, 21^. However, the apparently simple genomes of precancerous lesions at the clonal level does not exclude genome instability or complexity at the cellular level. Cells with unstable genomes will generate copy-number variation in the progeny^22, 23^, but such variation is invisible at the population level due to counterbalancing of random copy-number gains and losses in single cells in the absence of selection (i.e., neutral evolution). Genetic variation is further suppressed by positive selection (e.g., for oncogene amplifications) or negative selection (against large DNA deletions or aneuploidy in general^24^). Based on these considerations, we expect the footprint of genome instability in somatic genome evolution to be most visible in small precancerous lesions with *in situ* clonal expansion of copy-number variation generated by genome instability. This idea has led us to perform multiregional analysis of Barrett’s esophagus (BE)^25–27^ to dissect the origin of genome complexity in esophageal adenocarcinoma (EAC).

BE is the only known precursor of EAC and estimated to be present in 60-90% of newly diagnosed EAC cases^28^. In contrast to fully formed EACs with complex chromosomal changes^29^, BE tissue samples can contain lesions of different histopathological states with varying genomic complexity^30, 31^. By analyzing copy-number alterations in concurrent BE (both non-dysplastic and dysplastic) and early EAC (either intramucosal or T1) lesions, we reveal copy-number heterogeneity in BE cells before transformation, relate copy-number evolution patterns in BE cells to those derived from experimental models of chromosomal instability^32–38^, and provide mechanistic insight into the evolution of EAC genome complexity.

We find that both copy-number heterogeneity and complexity can predate the appearance of cancers or dysplastic lesions and are present in both single BE cells and BE subclones with intact p53. Importantly, p53 loss enables episodic but multigenerational genome evolution initiated by catastrophic events such as whole genome duplication^32, 33^, chromothripsis^34–36^, and dicentric chromosome formation^37, 38^: We provide evidence that both copy-number heterogeneity and complex copy-number gains in BE cells reflect multigenerational genome or chromosome instability precipitated by these events. We further demonstrate that ongoing chromosomal instability underlies both progressive DNA deletions in BE cells that result in sloping copy-number variation, and distinct oncogenic amplifications in independently transformed cancers within a single BE field. Together, these findings elucidate how genome instability drives copy-number evolution to promote tumor progression.

## Results

### Copy-number heterogeneity suggests early onset of chromosomal instability in precancer BE cells

Endoscopic mucosal resection (EMR) is routinely performed in patients with dysplastic BE. In reviewing more than 500 formalin-fixed, paraffin-embedded (FFPE) EMR samples, we identified 14 cases showing unexpected microscopic foci of invasive cancers and one case (Patient 1) with an early cancer removed via esophagectomy. All cancers were either intramucosal or T1 and all samples were collected before treatment. Following independent pathologic re-review by two or more pathologists to confirm the diagnoses (**Methods**), we delineated and performed laser capture microdissection (LCM) to isolate regions corresponding to distinct histopathological states^27^ (**Figure 1**), including non-intestinalized columnar metaplasia (COLME), non-dysplastic BE (NDBE), BE indefinite for dysplasia (IND), BE with low-grade dysplasia (LGD) or high-grade dysplasia (HGD), and intramucosal (IMEAC) or early EAC (**Extended Data Figure 1**). We further isolated normal tissue from benign FFPE regions that was used as germline reference.

**Figure 1:**
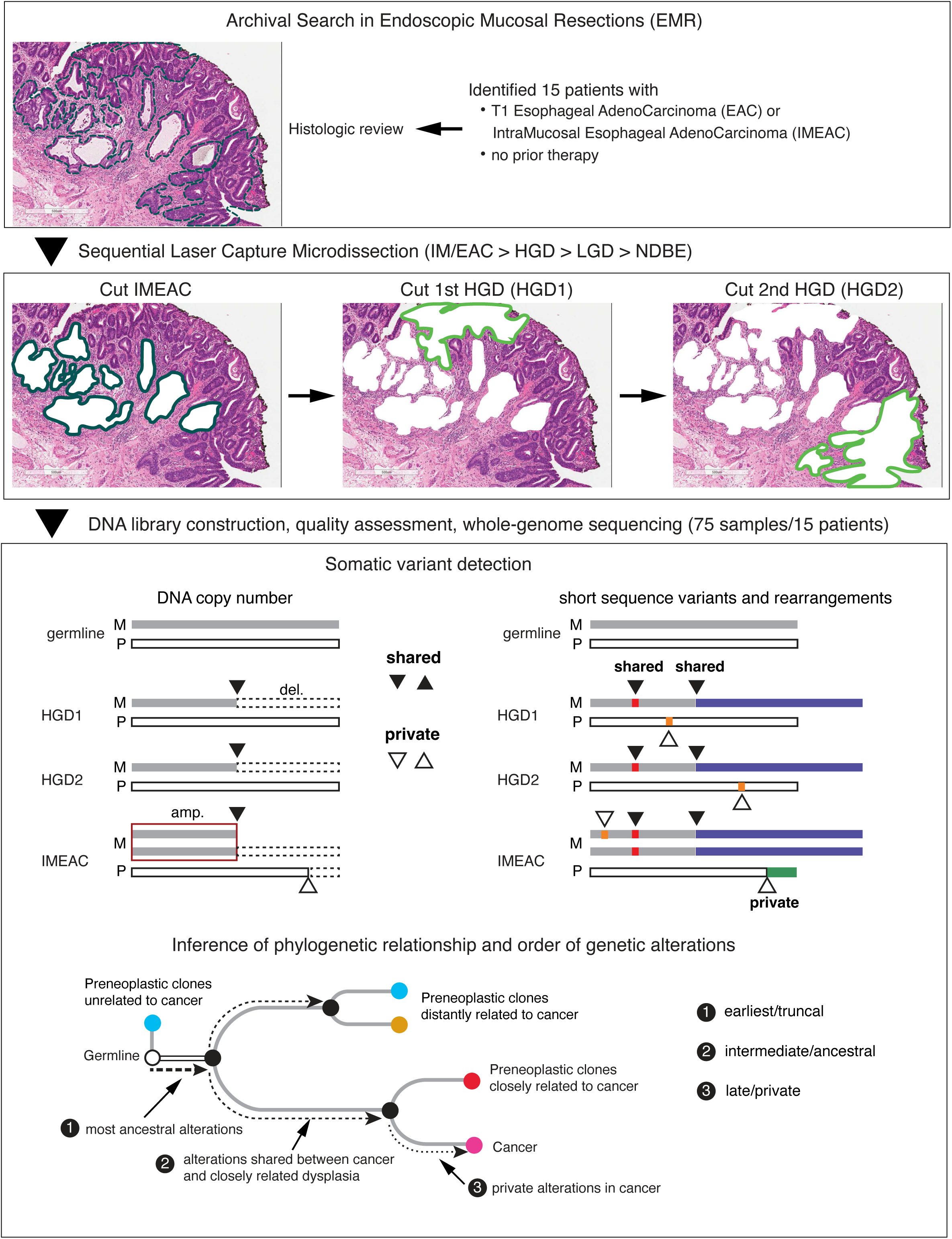
Overview of experimental design and bioinformatic analysis. Fifteen patients whose Barrett’s esophagus tissue samples presented early invasive esophageal adenocarcinomas (EAC) were selected. After histological review, 75 samples of early cancer (EAC) and precancerous lesions, including non-dysplastic Barrett’s Esophagus (NDBE), low-grade dysplasia (LGD) and high-grade dysplasia (HGD), were collected via laser capture microdissection and subjected to whole-genome sequencing. We perform joint variant detection on samples from each patient and then determine their phylogeny based on genetic alterations shared by two or more samples (filled triangles). Based on the phylogeny, we then infer the timing and evolution of copy-number alterations (both shared and private), including distinct copy-number changes on a single parental chromosome in related BE/EAC genomes generated by branching evolution.

Due to the limited quantity of FFPE DNA from small tissue sections and their lesser quality compared to DNA from fresh or frozen cells, we first performed low-pass whole-genome sequencing (WGS) at ∼0.1x mean depth to select libraries with sufficient complexity and then performed deeper sequencing ∼20x. The final cohort consisted of 75 BE/EAC (21 COLME/NDBE/IND, 7 LGD, 23 HGD, and 24 IM/EAC) and 15 reference samples from 15 patients (**Extended Data Table 1**). the variant calls generated by standard tools had both high false positive and high false negative detection rates (**Methods).** For single-nucleotide variants (both somatic and germline), short insertions/deletions, and rearrangements, we performed joint variant detection on all samples from each patient to improve variant detection accuracy (**Figure 1**). Although the joint analysis is sufficient to detect mutations shared by multiple samples, the false negative detection of mutations in individual samples due to sequencing dropout still confounds phylogenetic inference (**Methods**). To bypass this challenge, we focused on somatic copy-number alterations (SCNA) for which better accuracy could be achieved.

We determined chromosome-specific DNA copy number and copy-number changepoints based on haplotype-specific sequence coverage (**Methods, Supplementary Data**). Parental haplotypes were first inferred by statistical phasing using a reference haplotype panel^39^ and then refined based on allelic imbalance across all samples from each patient. We used haplotype-specific sequence coverage to first validate the estimated ploidies and clonal fractions of aneuploid BE/EAC clones and then calculate the integer DNA copy number of parental chromosomes. The determination of long-range parental haplotype both enabled phasing of SCNAs to parental chromosomes and ensured the accuracy of SCNA detection. We further performed segmentation of haplotype-specific DNA copy number and used copy-number changepoints to refine the list of rearrangements. For data presentation clarity, the copy-number plots in the main and extended data figures only show data of the altered homolog, except where stated. The haplotype-specific sequence coverage and copy number of both homologs are provided in **Supplementary Data**.

We determined the phylogenetic tree of samples from each patient (**Figure 2**) based on haplotype-specific copy-number alterations. SCNAs were first identified independently in each sample and then assigned to phylogenetic branches based on their presence or absence in all samples. The branch length (horizontal distance between nodes) approximately reflects the SCNA burden estimated using the number of altered chromosomes. SCNAs on each branch (labelled in **Extended Data Figure 2)** are summarized in **Extended Data Table 2**; SCNAs that affect esophageal cancer genes or identified more than once in the current cohort are annotated in **Figure 2**. In all but two patients (13 and 14), we identified SCNAs in related BE/EAC genomes affecting a single parental homolog but having distinct changepoints that indicate branching evolution of ancestral chromosomes; these chromosomes are labelled with asterisks near the inferred common ancestor. Whole-genome duplication (WGD) was inferred based on the number of homologous chromosomes with more than one copy^40^ and assigned to evolutionary branches based on the WGD status of individual samples (**Methods**). For SCNAs on branches with WGD, their timing relative to WGD was inferred based on the integer copy-number states. Finally, we confirmed the consistency between SCNA-derived phylogenetic trees and genetic similarities estimated from somatic SNVs (**Extended Data Figure 2**). The few instances of discrepancy are discussed in **Methods**.

**Figure 2:**
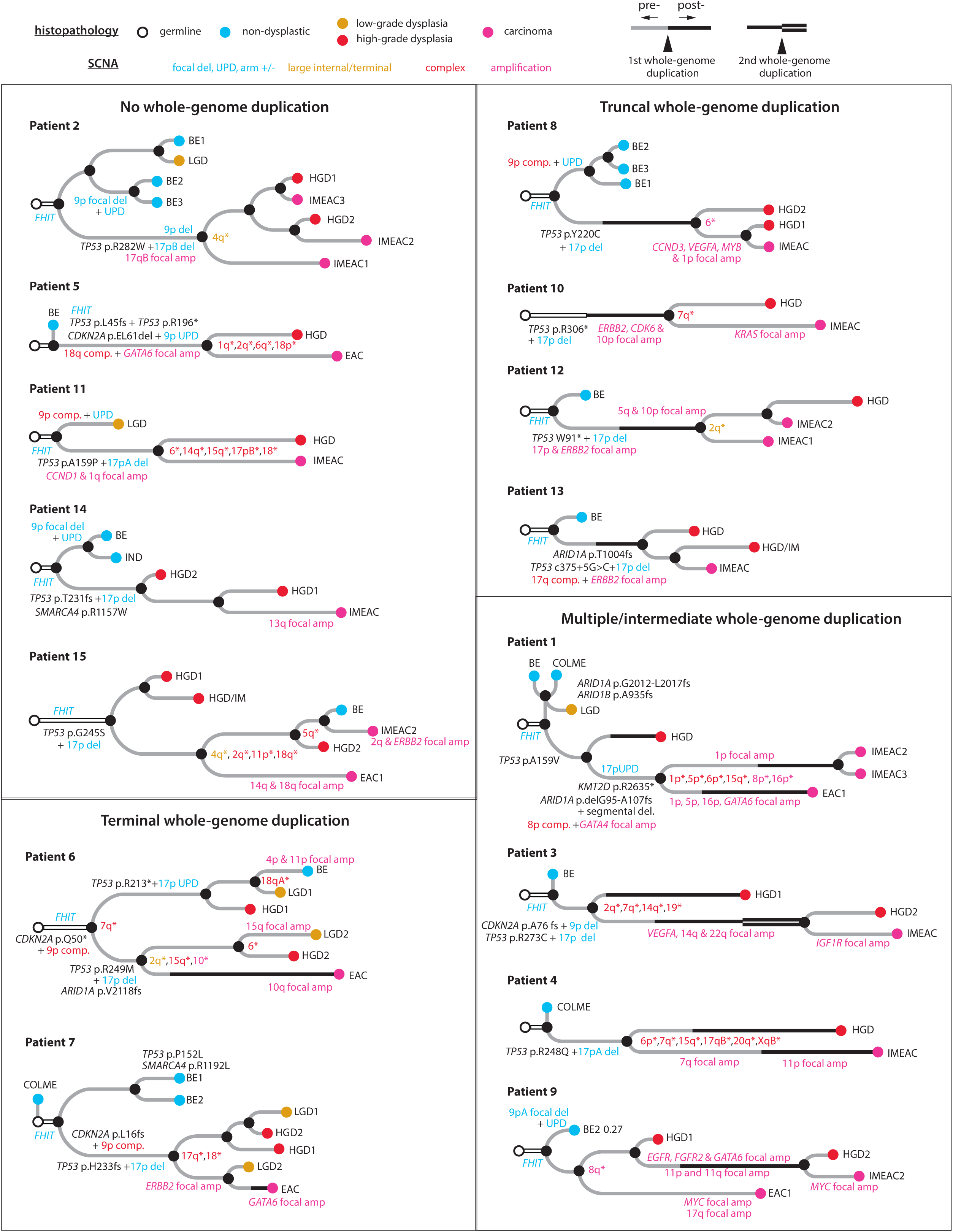
Phylogeny of early EAC and precursor BE lesions determined by haplotype-specific copy-number alterations. Phylogenetic trees are grouped based on the timing of whole-genome duplication (WGD, thick solid line) events. Samples are colored based on their histopathology grading: non-dysplastic (blue), low-grade dysplasia (orange), high-grade dysplasia (red), carcinoma (magenta). The branch length (horizontal distance between nodes) approximately reflects the number of altered chromosomes. For a complete list of alterations along each phylogenetic branch, see **Extended Data Table 2** and **Extended Data Figure 2**. Annotated alterations include: (1) recurrent alterations or those affecting known EAC drivers; (2) focally amplified regions or oncogenes (magenta); (3) chromosomes or chromosome arms (with asterisks) with divergent copy-number alterations in more than one progeny clones. Note that Patient 13 contained a splice-site mutation (c.375+5G>C) in *TP53* that was assessed to produce truncated p53^60^ and also reported to be a recurrent hospot in cancers in a recent study^61^. The colors of annotated chromosomes reflect the complexity of copy-number alterations: simple deletion/duplication, uniparental disomy, arm-level gain/loss (blue), large segmental (terminal or internal) copy-number changes or their combinations (orange), complex copy-number alterations (red), focal amplifications (magenta). For classification of copy-number alterations, see **Extended Data Figure 4**.

The phylogenetic trees of EAC and precursor BE lesions show several recurrent patterns. First, biallelic *TP53* inactivation is a truncal event of the evolutionary branches of cancer or high-grade BE lesions (14/15 patients). By contrast, focal deletion near *FHIT* (a common fragile site) is often ancestral to all BE and EAC lesions; bi-allelic inactivation of *CDKN2A* (a frequently inactivated tumor suppressor) can be truncal to either cancer/HGD lesions (Patient 3,5,6,7) or NDBE/LGD lesions (Patient 2,8,9,11,14). Second, evolutionary branches with the highest SCNA burdens are frequently associated with WGD, which is itself also a frequent event (10/15 patients). Third, high-grade dysplastic BE lesions and cancer lesions from the same patient often harbor distinct SCNA breakpoints on single parental chromosomes (13/15 patients) or distinct regions of focal amplification (10/15 patients), indicating copy-number heterogeneity prior to the emergence of aneuploid BE/EAC clones. Finally, we identified more than one early cancer lesion in five patients (Patient 1,2,9,12,15): The distinct cancer foci from each patient often displayed significant genomic divergence but were individually accompanied by precancerous lesions in close proximity (Patient 1,9,12,15) and/or showing more genomic similarity (Patients 2,9,12,15). The last observation strongly suggests that the cancer foci had evolved independently from distinct BE cells within the same BE field, i.e., independent malignant transformation.

The observation of significant SCNA diversity in BE and EAC subclones suggests highly dynamic copy-number evolution in precancerous BE cells and predicts copy-number diversity at the single-cell level. We directly tested this hypothesis by performing whole-genome sequencing analysis of 68 single cells isolated from a patient with known HGD by endoscopic cytology brushing immediately before radiofrequency ablation. We performed haplotype-specific copy-number analysis and phylogenetic inference using the same strategy as for bulk samples (**Methods**). We identified 12 cells with aneuploid genomes and 56 cells with near diploid genomes. Their phylogeny and selected examples of SCNAs in single BE cells or subclones are shown in **Figure 3;** SCNAs in each cell are listed in **Extended Data Table 3** and DNA copy-number plots of all cells are available in **Supplementary Data**. All the aneuploid cells share biallelic *TP53* inactivation through a pathogenic R175H mutation and loss-of-heterozygosity generated by 17p loss, but show significant heterogeneity of chromosomal copy-number changes. The onset of genomic heterogeneity in precancer BE cells following bi-allelic *TP53* inactivation recapitulates the pattern seen in bulk samples and provides direct evidence of dynamic precancer genome evolution driven by chromosomal instability. We next discuss specific patterns of copy-number evolution and their mechanistic implications.

**Figure 3:**
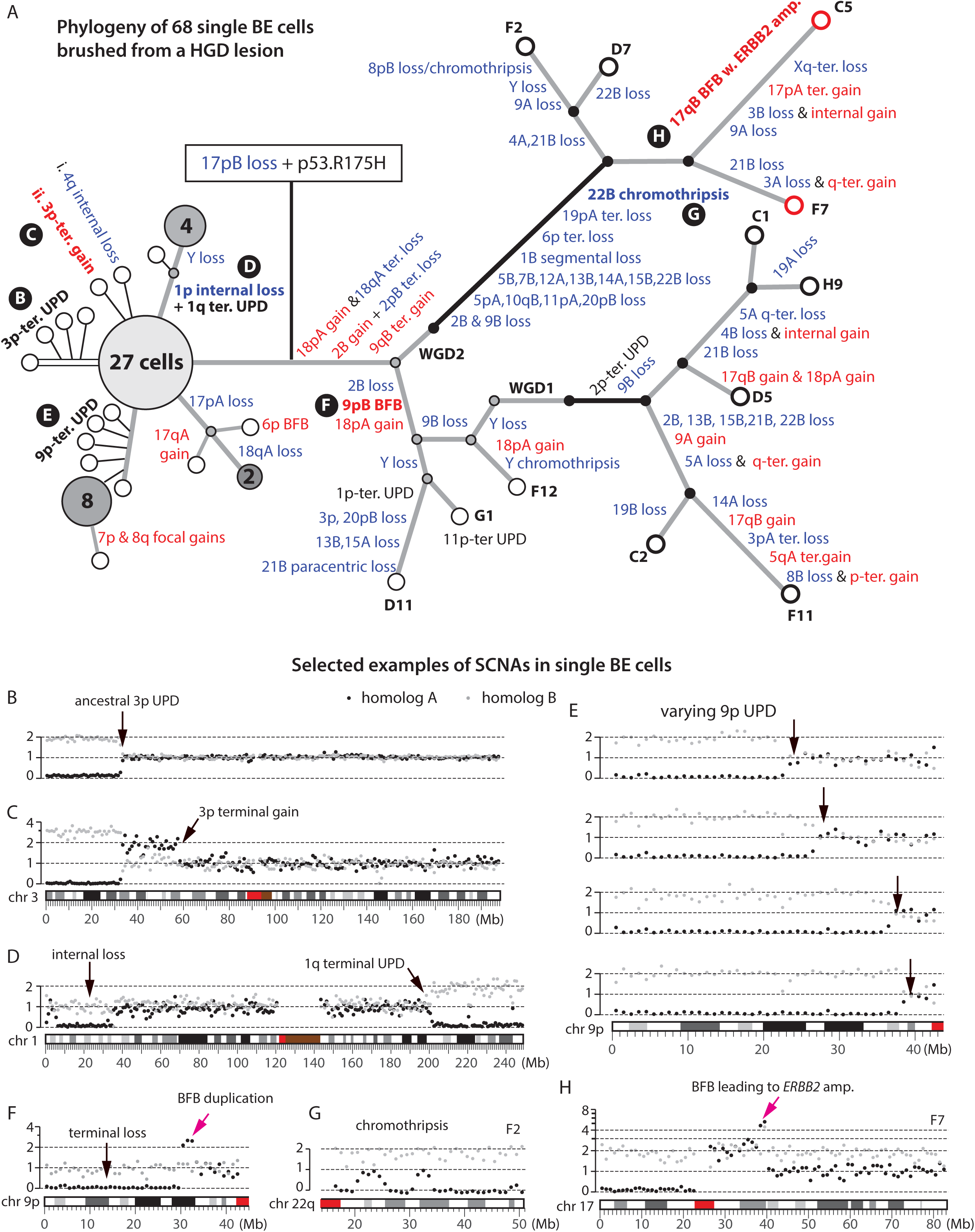
Copy-number evolution in 56 near diploid and 12 aneuploid BE cells from a high-grade dysplastic Barrett’s esophagus determined by single-cell sequencing. **A**. Phylogenetic tree with annotated haplotype-specific copy number alterations (blue for losses, red for gains). Open circles represent single cells; large filled circles represent subclones of cells (with annotated cell counts) with identical copy number; small filled circles represent inferred intermediate states (gray for pre-WGD, black for post-WGD). Aneuploid cells are separated into two branches each inferred to have undergone an independent whole-genome duplication (WGD) event. **B-H**. Examples of copy-number alterations before (B-E) and after (F-H) p53 inactivation. Gray and black dots represent haplotype-specific DNA copy number of parental chromosomes. **B**. Ancestral 3p uniparental disomy (UPD) shared by all but four cells. **C**. Sporadic 3p terminal gain after 3p UPD in one cell. **D**. Large paracentric deletion on 1p and UPD at the 1q-terminus shared by five cells. **E**. Progressive 9p UPD in a subclone of 14 cells. Only four cells are shown, see **Supplementary Data** for the others. **F**. Terminal duplication after terminal deletion on 9p shared by cell G1 and D11 that is consistent with two rounds of breakage-fusion-bridge cycles. **G**. Chromothripsis of Chr.22q shared by cell C5, F2, and F7. **H**. Focal amplification spanning the *ERBB2* gene on Chr.17 (∼40Mb) in cell C5 and F7 (red circles) that displays the signature copy-number pattern of breakage-fusion-bridge cycles. For a detailed list of alterations in each cell, see **Extended Data Table 3**.

### *TP53* inactivation and the onset of genome instability initiates BE genome evolution

We observed increasing SCNA burden with disease progression (**Figure 4A**,left; **Extended Data Fig. 3A and 3B**), but this correlation is mostly attributed to *TP53* mutation status. Samples with *TP53* inactivation show significantly higher SCNA burdens than samples without *TP53* inactivation (**Figure 4A**,middle; **Extended Data Fig. 3C**). In particular, two NDBE samples (from Patient 6 and 15) and four LGD samples (from Patient 6 and 7) with bi-allelic *TP53* inactivation show similar SCNA burdens as HGD and EAC samples; by contrast, NDBE and LGD samples without *TP53* inactivation show significantly fewer SCNAs (**Extended Data Fig. 3A**). These data and the contrasting SCNA burdens in single BE cells with and without intact p53 (**Figure 3A**) both reinforce the association between p53 loss and SCNA evolution^11, 31^.

**Figure 4:**
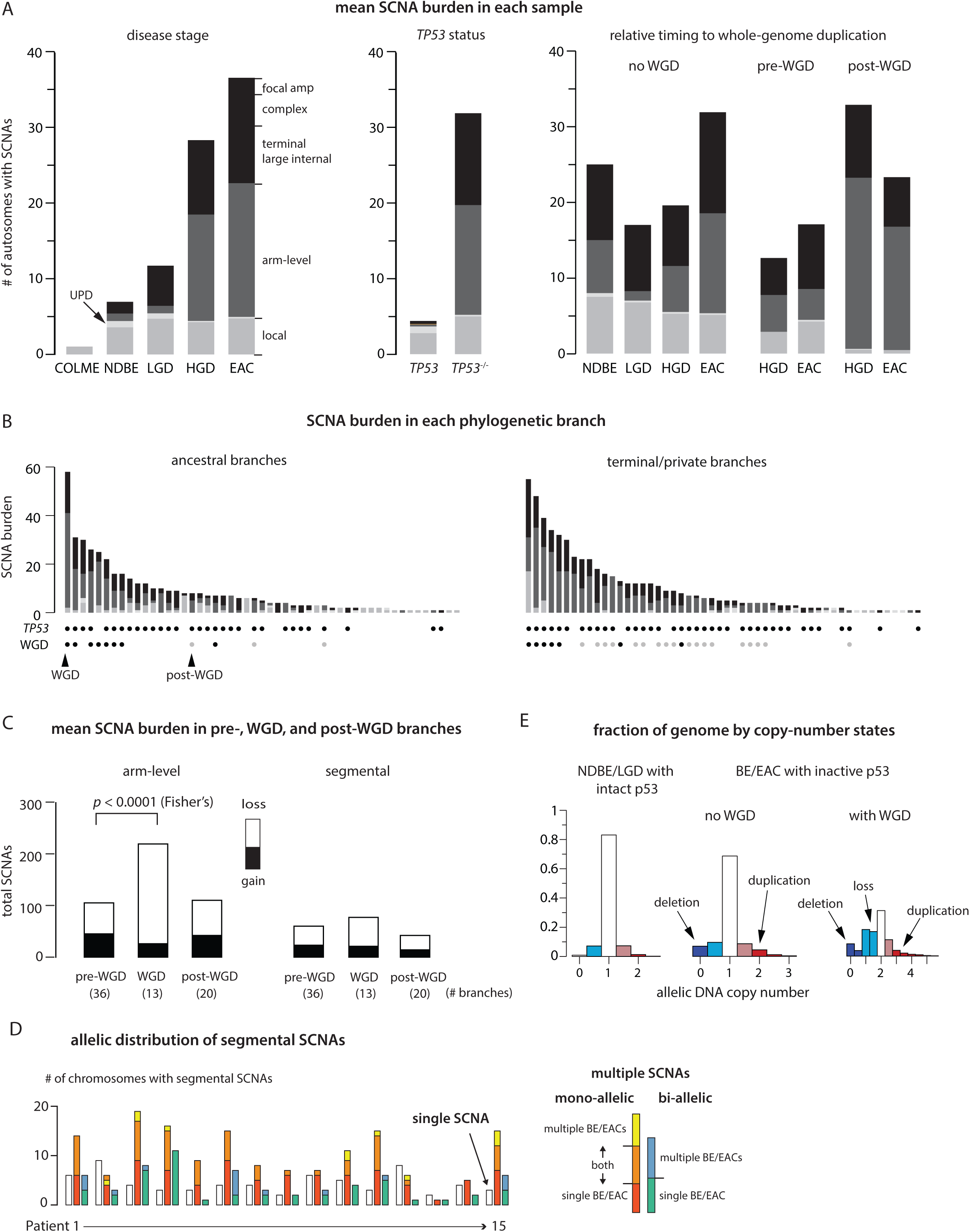
Landscape of somatic copy-number alterations (SCNA) in BE and EAC clones. **A**. Mean SCNA burden in samples grouped by disease stage (*left*), *TP53* mutation status (*middle*), and timing relative to whole-genome duplication (*right*). The SCNA burden is measured by the total number of altered autosomes (both parental homologs, maximum 44) and subdivided into local deletions or duplications (gray), uniparental disomies (light gray), arm-level SCNAs (dark gray), and segmental SCNAs (black). In the middle panel, the ‘intact’ *TP53* group (“*TP53*”) only includes NDBE/LGD samples without detectable *TP53* alterations, but not HGD/EAC samples. See **Extended Data Figure 3** for additional information including the SCNA burden in each sample. **B**. SCNA burden along ancestral (having more than one progeny clone) and terminal (only one progeny clone) phylogenetic branches. The bottom shows the *TP53* mutation status and the relative timing to WGD of each branch. **C**. Total counts of arm-level (left) and segmental (right) SCNAs (filled bars for gains, open bars for losses) in evolutionary branches preceding, concurrent with, or after WGD. Segmental SCNAs only include large internal/terminal SCNAs but not complex SCNAs that can generate both DNA gain and loss. The significantly higher burden of arm-level SCNAs in WGD-concurrent branches than pre-WGD branches (*p* < 10^-4^, Mann-Whitney) that is dominated by chromosome losses (*p* < 10^-4^, two-sided Fisher’s exact test) is consistent with episodic chromosome losses after tetraploidization. WGD is also associated with a modest but significant increase of segmental SCNA burden (*p* = 0.01, Mann-Whitney; WGD-concurrent vs pre-WGD) and of post-WGD arm-level SCNAs relative to pre-WGD branches (*p* = 0.01, Mann-Whitney). These can be explained by elevated rates of chromosome missegregation after tetraploidization that may also lead to complex segmental changes (e.g., from micronucleation). **D**. Allelic distribution of segmental SCNAs identified in all samples from each patient. Shown are the number of chromosomes (Chrs.1-22 and X) with single SCNAs (open bars), multiple SCNAs on a single parental homolog (‘mono-allelic’), and multiple SCNAs affecting both homologs (‘bi-allelic’). Mono-allelic and bi-allelic SCNAs with multiple breakpoints are further divided into subcategories based on whether SCNA breakpoints are found in a single BE/EAC genome, or in multiple related BE/EAC genomes. SCNAs (or SCNA breakpoints) concentrating on single parental chromosomes (mono-allelic) is consistent with either single catastrophic events (e.g., chromothripsis) or successive SCNA acquisition on single unstable chromosomes, whereas SCNAs affecting both parental chromosomes (bi-allelic) are consistent with independent SCNA acquisition. **E**. Fraction of the germline genome at different copy-number states (from 100kb-level allelic copy number). Deletion (dark blue), subclonal deletion/loss (light blue), subclonal gain (light red), or duplication (dark red). There is a marked increase in the fractions of both deleted (CN = 0) and duplicated DNA (CN ≥ 2) in BE/EAC genomes with inactive p53 compared to BE genomes with intact p53. Samples with WGD also have a larger fraction of the genome at the single-copy state reflecting DNA loss after WGD.

Prior analyses of ageing esophageal tissues^9, 10^ by bulk sequencing revealed uniparental disomy (UPD), or copy-neutral loss of heterozygosity, as the only large segmental SCNA. Consistent with this observation, we observed frequent UPDs in both single BE cells (**Extended Data Table 3**) and clones (**Extended Data Table 4**) prior to p53 loss, but only sporadic segmental gains or losses in single BE cells (**Figure 3C,D**) and almost none in BE clones. Remarkably, we identified UPDs on the 9p terminus with varying boundaries in a subclone of 14 single BE cells (**Figure 3E** and **Supplementary Data**). As this variation does not alter total DNA copy number, it can only be revealed by haplotype-resolved copy-number analysis. The varying boundaries of terminal UPD in different cells (arrows in **Figure 3E**) bear an intriguing similarity to our prior observation of varying terminal deletions attributed to ongoing breakage-fusion-bridge cycles^38^ (see **Extended Data Figure 7** that will be discussed later). The similarity between varying terminal UPDs and varying terminal deletions suggests a plausible common origin from broken chromosomes generated by breakage-fusion-bridge cycles^23^, with deletions resulting from translocations involving other broken ends and UPDs resulting from homology-dependent invasion of broken ends into the intact homolog followed by a half crossover resolution^41^ (**Extended Data Fig. 4**, top).

In contrast to the simple SCNA landscape in BE cells with intact p53 is the prevalence of arm-level and complex SCNAs in BE cells and clones after p53 loss. Loss of p53 does not directly cause aneuploidy or chromosomal instability in human cells^42^, but abolishes p53-dependent arrest after DNA damage^43^ or prolonged mitosis^44^. The burst of SCNA complexity after p53 loss is therefore more reflective of an increased frequency of SCNA clonal expansion than an increased rate of SCNA acquisition. Moreover, the observation of sporadic large SCNAs, especially UPDs, in single BE cells with intact p53 indicates that BE cells do acquire DNA breaks, but these breaks do not lead to complex copy-number alterations as seen in BE cells or clones with inactive p53. We next focus on BE cells or clones with inactive p53 and provide evidence supporting that the accumulation of SCNA complexity reflects multigenerational chromosomal instability that is precipitated by sporadic cell division errors but only propagated after p53 inactivation.

### Whole-genome duplication triggers rapid accumulation of arm-level copy-number changes

The most dramatic change in BE cells is whole-genome duplication (WGD). WGD is inferred to be a frequent event in many epithelial cancers^45, 46^ and thought to define a particular EAC evolution trajectory^31^. We inferred 15 WGD events in bulk BE/EAC lesions from 10/15 patients, including independent WGD occurrences in distinct HGD/EACs from Patient 1,3, and 4 (**Figure 2**). We further inferred two independent WGDs in single BE cells without presence of cancer (**Figure 3A**). These observations suggest that WGD may occur frequently during BE progression before the appearance of cancer.

Despite the prevalence of WGD in human cancers^45, 46^ and its tumor-promoting capacity^47, 48^, how WGD impacts tumorigenesis remains incompletely understood. One proposal is that tetraploidization (the event that causes WGD) can precipitate additional genome instability including multipolar cell division or chromosome missegregation^6, 32, 33^ that leads to aneuploidy. Consistent with this model, we inferred that more SCNAs in BE/EAC genomes were acquired after WGD than before WGD (**Figure 4A**, right), and evolution branches with WGD acquisition had significantly higher SCNA burdens (30 events/branch) than non-WGD branches (pre-WGD: 7.5/branch; post-WGD: 8.8/branch) (**Figure 4B, Extended Data Table 2**). Moreover, a majority of post-WGD SCNAs are arm-level changes (302 out of 428 events) and dominated by losses (256) (**Figure 4C**), a pattern also seen in single aneuploid BE cells (**Figure 3A**).

The preponderance of chromosome losses after WGD has two implications. First, this pattern cannot be solely explained by increased rates of random chromosome missegregation^32^ that generates reciprocal gain and loss in a pair of daughter cells. This pattern could reflect a lower fitness of cells with larger chromosome number due to more frequent mitotic delays and defects^46^. It could arise from multipolar cell divisions that generate three or more progeny cells with predominantly chromosome losses^33^ (**Extended Data Fig. 5A**). Future work is needed to test these hypotheses. Second, extensive chromosome losses after WGD may significantly reduce the number of duplicated chromosomes and cause underestimation of WGD incidence in cancer development, especially in cancers with highly aneuploid genomes. Together, our analysis of arm-level SCNAs in BE cells both confirms WGD as a precursor to aneuploidy^49–51^ and highlights the diversity of copy-number outcomes^5^ generated by post-WGD events including multipolar cell division^33^.

### Segmental copy-number alterations display signatures of dicentric chromosome evolution

In contrast to the prevalence of post-WGD arm-level SCNAs, we inferred a similar number of segmental SCNAs in BE/EAC genomes to have occurred prior to (135) and after WGD (126) in samples with WGD acquisition. The fractions of segmental DNA loss and DNA gain are also comparable among pre-, post-, and WGD branches (**Figure 4C**, right), although branches with WGD acquisition have a higher average SCNA burden (5.9 events) than pre- (1.6) or post-WGD (2.1) branches. These observations indicate that segmental SCNA acquisition is promoted by WGD but also occurs independent of WGD.

Segmental SCNAs in BE genomes further display two features of non-randomness. First, SCNA breakpoints are often concentrated on a few chromosomes with complex deletions (chromothripsis) or duplications. Second, distinct SCNAs in related BE/EAC genomes more frequently originate from a single parental chromosome (‘mono-allelic’) than affect both parental chromosomes (‘bi-allelic’) (**Figure 4D** and **Extended Data Fig. 3E**). Both features are consistent with one-off or successive SCNA acquisition on individual unstable chromosomes instead of independent SCNA acquisition across the genome. The connection between segmental SCNA acquisition and chromosomal instability is further supported by the observation of larger fractions of deletions (allelic copy number = 0) or duplications (allelic copy number ≥2 in non-WGD samples and ≥3 in WGD samples) in samples with inactive p53 than in samples with intact p53 (**Figure 4E**). Finally, we recognized that many segmental SCNA patterns in BE/EAC genomes are consistent with the outcomes of chromosomal instability from abnormal nuclear structures including micronuclei^34^ (**Extended Data Fig. 5B**) and chromosome bridges (**Extended Data Fig. 5C**)^38^. We sought to use the genomic signatures of *in vitro* chromosomal instability to deconvolute segmental copy-number complexity in BE/EAC genomes.

The most frequent SCNAs in BE/EAC genomes are gain or loss of large terminal (i.e., spanning a telomere) or internal (with two non-telomeric breakpoints) segments; these alterations are consistent with the outcomes of dicentric chromosome breakage (**Figure 5**). Dicentric chromosomes can result from either end-to-end chromosome fusion or incomplete decatenation of sister chromatids^38^ and lead to a ‘bridge’ between daughter nuclei when the two centromeres segregate to different daughter nuclei. Although dicentric chromosomes can be generated by a variety of mechanisms, the genomic consequences are primarily determined by the formation and breakage of chromosome bridges^37, 38^. Breakage of a single dicentric chromosome (‘chromatid-type’ bridges) will generate reciprocal gain and loss of a telomeric segment (‘terminal’ SCNAs) (**Figure 5A**). If both sister dicentric chromatids are part of the bridge (‘chromosome-type’ bridges), their breakage can give rise to large segmental gain or loss within a chromosome arm, hereafter referred to as ‘paracentric’ SCNAs (**Figure 5B**). Both of these outcomes were directly demonstrated in single-cell experiments^38^ but originally described by McClintock (summarized in Ref.^52^) We further observed large SCNAs spanning centromeres (‘pericentric’ SCNAs) that can result from broken ring chromosomes (**Figure 5C**, first described by McClintock in Ref.^53^) or multicentric chromosomes. The instances of terminal and large internal SCNAs in our BE/EAC cohort are summarized in **Figure 5D** and listed in **Extended Data Table 5:Tab 1**. In total, these events account for ∼50% of segmental SCNAs.

**Figure 5:**
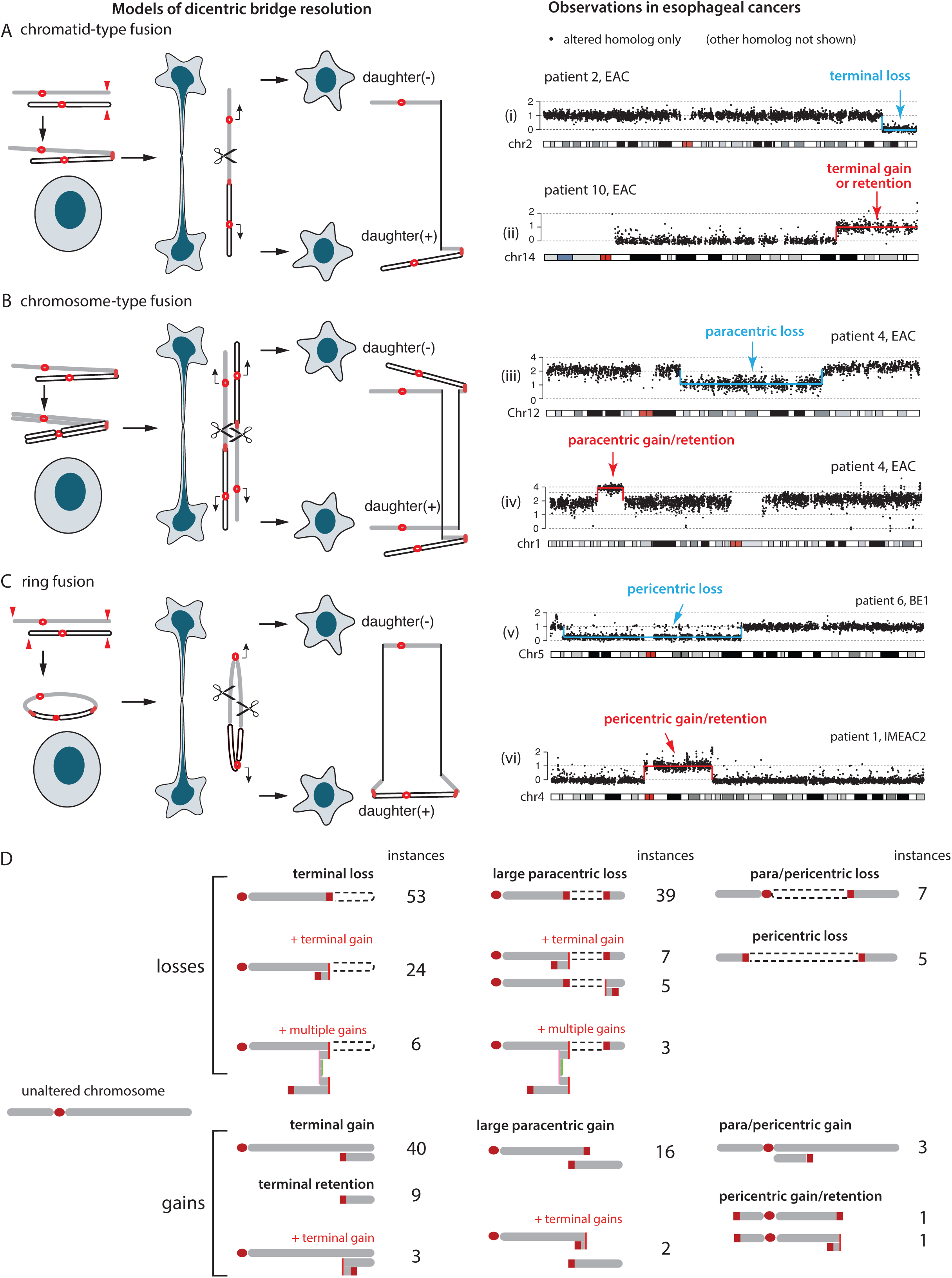
Segmental copy-number alterations in BE/EAC genomes that match the outcomes of dicentric chromosome bridge resolution. **A-C.** (*Left*) Different types of dicentric chromosome breakage and their copy-number outcomes: (A) terminal; (B) paracentric; or (C) pericentric segmental copy number changes. The open and filled chromatids may be sister chromatids or different chromosomes. Both A and B were demonstrated *in vitro* in Umbreit et al. (2020). The model that pericentric copy-number changes may arise from broken dicentric ring chromosomes (C) or multicentric chromosomes (not shown) has not been demonstrated *in vitro* but is plausible as telomere crisis may lead to multiple critically shortened telomeres. Examples of ring chromosomes (Umbreit et al., 2020) or rearrangements involving multiple chromosomes (Maciejowski et al., 2015) were also observed in the progeny populations of cells that have undergone telomere crisis or bridge induction. (*Right*) Examples from BE/EAC genomes that recapitulate the copy-number outcomes of bridge resolution. The allelic copy-number plots (25kb bins) show the DNA copy number of the altered chromosome; the intact homolog is not shown. Examples of gain and loss in each group are unrelated. See **Downloadable Supplementary Data** for the copy-number plots of both homologs in each sample. **D.** Summary of terminal/internal SCNAs in BE/EAC genomes. The number of instances is shown next to the copy-number pattern generated by different copy-number outcomes of BFB cycles. See **Extended Data Table 5** for the complete list.

Although chromosome bridge resolution provides a simple mechanism for single-copy gain or loss of large segments, similar copy-number outcomes may be generated by other processes. For example, terminal deletion or duplication could result from simple chromosomal translocations followed by whole-chromosome losses or gains (**Extended Data Fig. 6A**). This model, however, produces an equal number of terminal gains (including retentions) and losses, and cannot explain the disparity between terminal gains and losses seen in most samples (**Extended Data Fig. 6B**). Moreover, as broken bridge chromosomes can form new dicentrics and undergo breakage-fusion-bridge (BFB) cycles that generate a variety of compound copy-number outcomes, the identification of these compound copy-number patterns in BE/EAC genomes provides stronger evidence of chromosome bridges being involved in BE copy-number evolution.

The most common outcome of BFB cycles is the presence of DNA duplications near the boundaries of large segmental deletions (**Figure 6A,B**) or large segmental gains. Instances of these patterns in BE/EAC genomes are listed in **Extended Data Table 5:Tab 2** and also summarized in **Figure 5D**. The identification of interchromosomal rearrangements between both simple and compound SCNA breakpoints (**Figure 6A,B** and **Extended Data Figure 6C,D**) also suggests that these broken ends were generated simultaneously, most likely from the resolution of multichromosomal bridges as seen in experimental models of telomere crisis^37^ or chromosome bridge resolution^38^.

**Figure 6:**
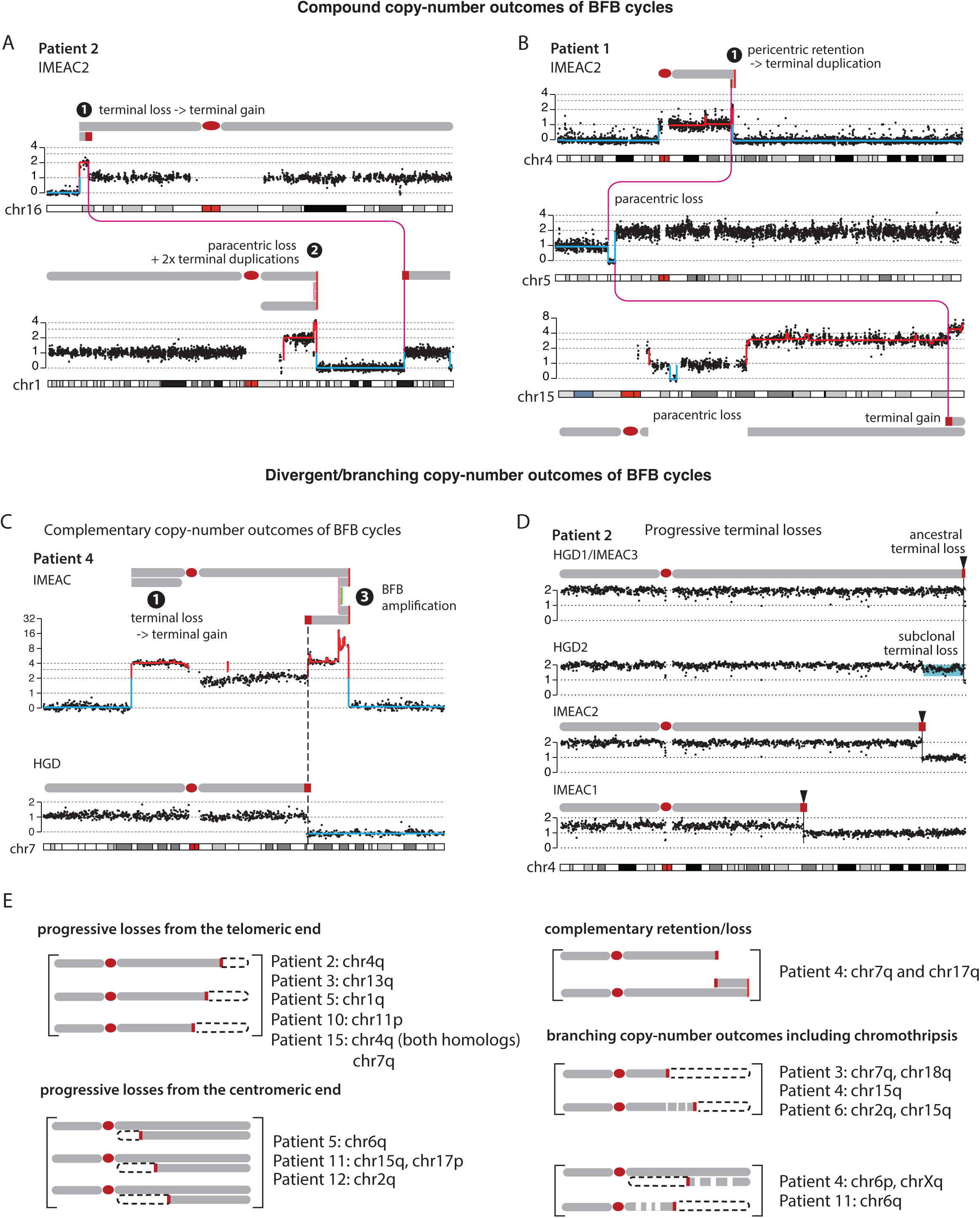
Segmental copy-number patterns consistent with multigenerational breakage-fusion-bridge cycles. Arabic numbers represent different BFB outcomes that are also labelled in **Extended Data Fig. 5D**. Schematic diagrams of altered chromosomes are drawn according to the segmental DNA copy number. **A**. (*Top*) Terminal deletion -> terminal duplication; (*bottom*) paracentric deletion -> two duplications near the centromeric break end. **B**. (*Top*) Pericentric retention -> terminal duplication at the q-terminus; (*middle*) paracentric deletion -> whole-chromosome duplication of the centromeric segment; (*bottom*) terminal gain or pericentric deletion after whole-chromosome gain. Magenta lines represent translocations between broken fragments. See **Extended Data Figure 6** for more examples. **C**. Complementary copy-number gain and loss at a single breakpoint (dashed line) in HGD and IMEAC reflect two broken pieces of a single dicentric chromosome. The focally amplified region on the telomeric end in IMEAC is consistent with preceding BFB amplifications. **D**. A series of terminal deletions on the same parental chromosome seen in five lesions from Patient 2. The proximal boundaries of the subclonal DNA loss near the 4q-terminus in HGD2 and clonal DNA loss in IMEAC2 suggest that IMEAC2 may have evolved from a subclone in HGD2. See **Extended Data Figure 7** for an example of the same pattern revealed in experimental BFB evolution. **E**. Summary of SCNAs in related BE/EAC genomes reflecting divergent/branching BFB outcomes. See **Downloadable Supplementary Data** for the copy-number plots of each instance.

Successive DNA duplications at the broken ends of chromosomes can generate focal amplifications (**Figure 6C**, top). Remarkably, the amplification on 7q in IMEAC shares a common SCNA boundary with the terminal deletion in HGD. (The same pattern of reciprocal DNA retention and loss is also seen in 17q of these two clones.) This pattern of reciprocal DNA retention and deletion directly recapitulates the outcome of broken bridge chromosomes between daughter nuclei (**Figure 5A**) that is only visible by multiregional sequencing. Based on this observation, we inferred that the HGD and the IMEAC clones were independently derived from sibling cells each having inherited a broken piece of a dicentric Chr.7 with amplified DNA that was present in their common ancestor.

Besides DNA duplications at broken termini, BFB cycles can also generate progressive DNA losses from either sequential breakage or deficient replication of bridge chromatin^38^. As each new deletion erases the boundary of preceding deletions, progressive DNA losses can only be revealed in different progeny clones (**Extended Data Figure 7**) but not in a single clone. We observed 11 instances of terminal or paracentric SCNAs with distinct breakpoints in different BE/EAC lesions from the same patient that are consistent with progressive DNA losses (**Extended Data Table 6:Tab1**). One example of varying 4q-terminal losses (boundaries marked by black arrows) in five lesions from Patient 2 is shown in **Figure 6D**.

In summary, we identified frequent duplications or deletions of large terminal, paracentric, and pericentric segments in BE genomes and attributed them to the formation and breakage of dicentric chromosomes (**Figure 5**). This mechanistic association is further supported by the observation of (1) additional duplications or progressive DNA losses at SCNA boundaries (**Figure 6**) reflecting successive BFB cycles (**Extended Data Fig. 7**); and (2) interchromosomal translocations between SCNA boundaries indicating simultaneous generation of broken chromosome ends. In particular, the observation of reciprocal DNA loss and gain in distinct BE/EAC clones from the same patient that directly recapitulate the outcome of dicentric bridge resolution between daughter cells (**Figure 6C**) provides the most compelling evidence of BFB cycles during BE evolution.

### Contemporaneous chromothripsis and BFB cycles generate EAC copy-number complexity

Besides simple DNA loss and gain, dicentric chromosomes can also undergo DNA fragmentation^37, 38^ either from chromosome bridge resolution or in micronuclei from chromosome missegregation. These processes generate chromothripsis with distinct oscillating DNA copy number patterns. For chromothripsis from bridge resolution, fragmentation of the bridge chromatin creates oscillating copy number in a fraction of the chromosome arm that was in the bridge, and the region with oscillating copy number is usually adjacent to the boundaries of large terminal or internal SCNAs corresponding to termini of broken bridge chromosomes (**Extended Data Fig. 8A**). We inferred that 35 instances of chromothripsis were consistent with this pattern (**Extended Data Table 7:Tab1**, ‘direct’ in Column N) and show representative examples in **Extended Data Fig. 8B-D**. For chromothripsis resulting from fragmentation of dicentric chromosomes partitioned into micronuclei, the oscillating copy-number pattern should span whole chromosome arms (“chromosome/arm”) (**Extended Data Fig. 8E**). We inferred that 25 instances of chromothripsis were consistent with this evolution sequence (**Extended Data Table 7:Tab1**, ‘downstream’ in Column N). The second scenario is best demonstrated in the example shown in **Extended Data Fig. 8F**, where the three-state oscillating copy-number pattern (CN=0,1,2) spanning both Chr.17q and 18p together with inter-chromosomal rearrangements indicated chromothripsis of a dicentric translocated chromosome. We additionally identified 40 instances of chromothripsis spanning entire chromosomes or arms that are consistent with micronucleation and 7 instances of regional chromothripsis without a clear relationship to large terminal/internal SCNAs.

We further analyzed DNA rearrangements related to chromothripsis but restricted this analysis to ancestral chromothripsis shared by three or more samples for which joint rearrangement detection can achieve good accuracy (see **Methods**). We identified two examples of chromothripsis involving subchromosomal regions (including arms) from multiple chromosomes (**Extended Data Fig. 8F,G**) that are consistent with multichromosomal bridge resolution. In two instances of chromothripsis, we further identified clustered rearrangement breakpoints near single SCNA boundaries (**Extended Data Fig. 8D,G**) that resemble the tandem-short-templates rearrangement pattern observed in chromothripsis from bridge resolution^38^ and micronucleation^34^. These rearrangement patterns provide additional evidence supporting the connection between chromothripsis and chromosomal bridges or subsequent micronuclei.

The comparison of SCNAs in related BE/EAC genomes provides further evidence for BFB cycles in BE genome evolution. In the example shown in **Figure 7A**, the ancestral paracentric deletion shared by all three genomes (LGD2/HGD3/EAC) was followed by regional chromothripsis and BFB amplifications near the centromeric break end in the LGD2 clone and a terminal duplication near the telomeric break end in the EAC clone; both downstream alterations likely arose from secondary BFB cycles after the ancestral paracentric deletion. In the example shown in **Figure 7B**, the (mostly) non-overlapping segments retained by the HGD and IMEAC genomes is consistent with a random distribution of DNA fragments from a single micronuclear chromosome into a pair of daughter cells^34^. Other examples of chromothripsis as one of the branching outcomes of BFB cycles are listed in **Extended Data Table 6** and **Figure 6E**.

**Figure 7:**
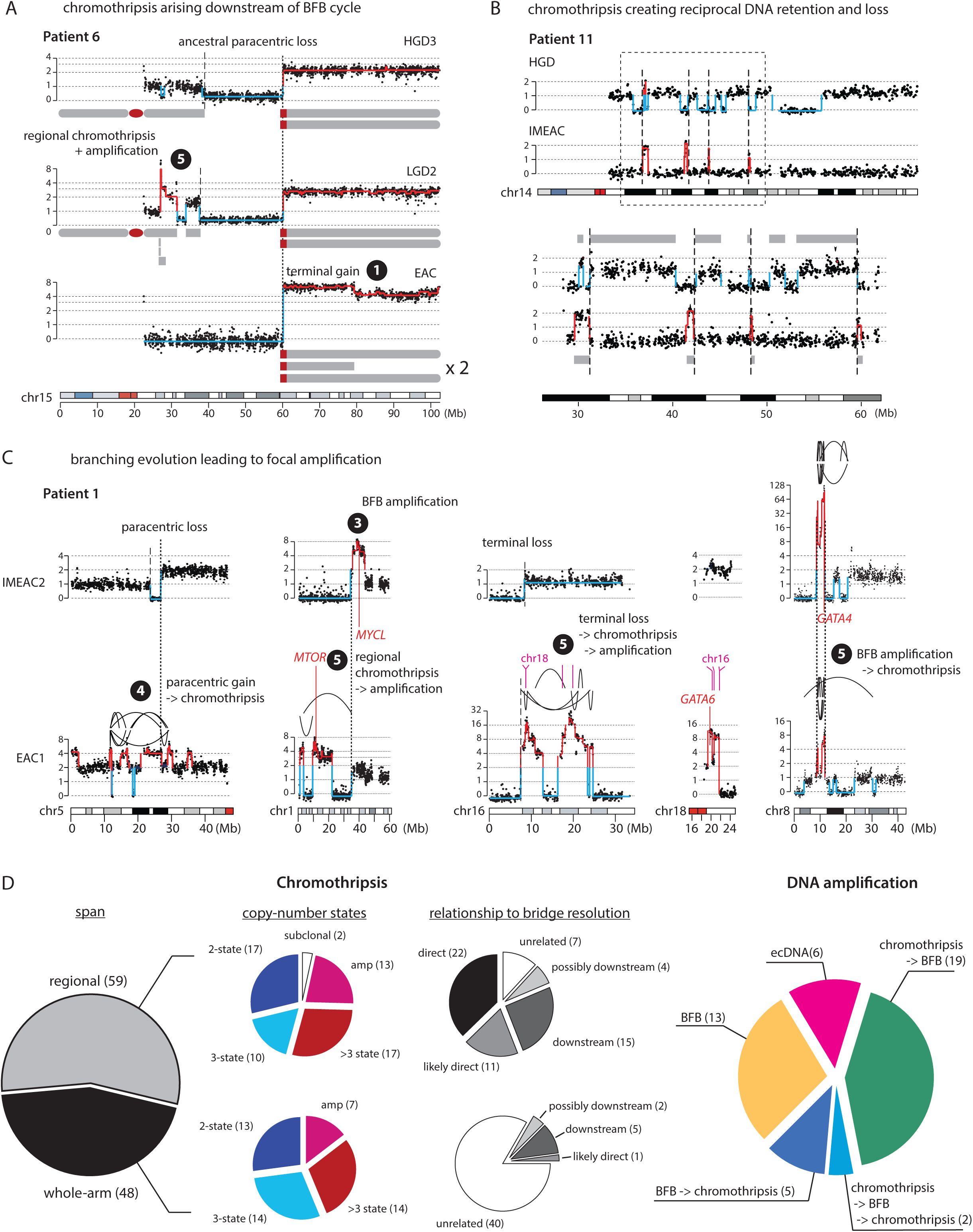
Complex SCNAs in BE/EAC genomes indicating successive chromothripsis and BFB cycles. Arabic numbers represent different BFB outcomes that are also labelled in **Extended Data Fig. 5D**. **A**. Divergent chromothripsis (in LGD2) and terminal duplication (EAC) occurring downstream of an ancestral paracentric deletion in Patient 6. The dotted line represents the ancestral breakpoint shared by all three genomes; dashed lines represent private SCNA breakpoints. **B**. Reciprocal distribution of Chr.14q in HGD and IMEAC lesions from Patient 11. The bottom shows an enlarged view of the outlined region (dashed box). Except for a small segment near 30Mb, all the other segments retained in the IMEAC genome are lost from the HGD genome. Dashed lines denote SCNA breakpoints with opposite retention and loss in the two genomes. **C**. Five subchromosomal regions with distinct copy-number patterns in two cancer lesions from Patient 1. For regions on 5p, 1p, and 8p, we infer the SCNAs evolved from a single unstable ancestor chromosome based on shared SCNA breakpoints (dotted lines). For 16p, the SCNAs are related by a common region of terminal deletion with adjacent boundaries (dashed lines). The amplified regions on 16p in EAC1 are joined to the amplified region on 18q spanning *GATA6*. The order of chromothripsis and amplification is determined based on whether the amplified regions are interrupted by deletions (indicating chromothripsis before amplification) or peppered with DNA losses (indicating chromothripsis after amplification). **D**. Summary of chromothripsis and DNA amplification instances grouped by copy-number features and the inferred evolutionary sequences. The inference of chromothripsis arising either directly from or downstream of dicentric chromosome breakage is based on the span of oscillating copy-number pattern relative to entire chromosomes; instances with less certainty are annotated accordingly (“possibly downstream” or “likely direct”).

The examples in **Figure 7A** and **7B** illustrate how copy-number breakpoints with either identical (**Figure 7A**, dotted line) or complementary (**Figure 7B**, dashed lines) DNA retention and loss in related genomes can inform about the evolutionary sequence of the observed copy-number alterations. This is further demonstrated in the Chr5 example in **Figure 7C**. The shared copy-number breakpoint (dotted line) with complementary DNA retention and deletion in IMEAC2 and EAC1 indicates a reciprocal distribution of broken chromosome fragments into their ancestors; the paracentric loss in IMEAC2 further suggests a chromosome-type bridge breakage event (**Figure 5B**). Therefore, the chromothripsis alteration with three oscillating copy-number states in EAC1 must have arisen downstream of the ancestral breakage event.

The combination of chromothripsis and successive DNA duplications in BFB cycles can explain complex segmental gains and amplifications. Whereas simple BFB cycles generate duplications flanked by large segmental deletions (**Figure 5D, Extended Data Fig. 8H**), BFB cycles following chromothripsis generate segmental gains or amplifications with interspersed DNA deletions (**Extended Data Fig. 8I**). Several copy-number patterns in Patient 1 indicate contemporaneous chromothripsis and BFB amplifications (**Figure 7C**). On both Chr.1p and Chr.16p, the oscillation between DNA deletion and amplification in EAC1 suggests an evolution sequence of ancestral chromothripsis followed by downstream BFB amplifications; the same regions in IMEAC2 display terminal duplications (Chr.1p) and a simple terminal deletion (Chr.16p). The presence of a shared copy-number breakpoint on Chr.1p and a common region of terminal deletion on Chr.16p between the EAC1 and IMEAC2 genomes suggests that the distinct copy-number patterns reflect divergent evolutionary outcomes of a single ancestral broken chromosome. Interestingly, the amplified regions on 16p in the EAC1 genome do not contain known oncogenes but are co-amplified with a region on 18q containing *GATA6*, a recurrently amplified EAC oncogene. By contrast, the IMEAC2 genome harbors neither amplification but has more amplified *GATA4* on Chr.8p. Moreover, the shared boundaries of amplified regions on 8p in both EAC1 and IMEAC2 indicates that the *GATA4* amplification was ancestral to both genomes but underwent different downstream evolution. The distinct *GATA4* and *GATA6* amplifications in these two genomes, likely reflective of positive selection for their combined expression^54^, highlights how persistent chromosomal instability rapidly generates copy-number heterogeneity and fuels the acquisition of oncogenic amplifications.

As DNA amplification is only one out of many possible outcomes of multigenerational copy-number evolution (we operationally defined focally amplified regions to have allelic copy number ≥ 8 that can be attained with at least three rounds of duplications), clonally fixated amplifications are likely reflective of positive selection and expected to contain oncogenes. Among 45 focally amplified regions each spanning one or multiple loci on a chromosome (**Extended Data Table 7:Tab2**), 24 encompass putative oncogenes and 29 overlap with regions that are recurrently amplified in cancer. The significance of focal amplification as a mechanism of oncogenic activation during EAC transformation^30, 31^ is further supported by the observation of both recurrent amplifications of EAC oncogenes, including *ERBB2* on 17q (5/15 patients) (**Extended Data Fig. 8H,I**) and *GATA6* on 18q (4/15 patients), and sporadic oncogene amplifications that are exclusive to cancer lesions but not their precursors, including *IGF1R* (Patient 3), *MET* (Patient 4), and *KRAS* (Patient 10).

In summary, we found that many complex segmental copy-number alterations in BE/EAC genomes, including focal amplifications, can be deconvoluted into different evolution sequences of sequence duplications generated by BFB cycles and chromothripsis from DNA fragmentation (**Figure 7D**). Together with observations of terminal/internal SCNAs reflecting simple copy-number outcomes of BFB cycles, these data provide *in vivo* evidence for the involvement of abnormal nuclear structures including micronuclei^34–36^ and chromosome bridges^37, 38^ in the generation of EAC genome complexity.

### Chromosomal instability generates continuous copy-number variation prior to discrete changes

Our analysis of BE/EAC genomes reveals both copy-number complexity and copy-number heterogeneity in BE subclones that indicate multigenerational evolution of unstable chromosomes. Importantly, chromosomal instability first generates copy-number variation in single BE cells. We wondered whether such instability in single BE cells can be discerned prior to copy-number heterogeneity or complexity in BE subclones.

If chromosome breakage only generates reciprocal DNA retention and loss between sibling cells, such changes are not visible at the clonal level as there is not net DNA gain or loss. However, we previously demonstrated that chromosomes in both micronuclei and bridges undergo deficient DNA replication leading to net DNA losses^34, 38^. If broken chromosomes remain mitotically unstable for multiple generations, successive under-replication of the broken termini can generate varying terminal losses in the progeny population (**Figure 8A**) that lead to ‘sloping’ copy number variation (**Extended Data Figure 7**). We identified sloping copy-number variation on three chromosomes in the HGD sample from Patient10 (**Figure 8B**). The constant DNA copy number of the intact homolog (gray) establishes that the sloping copy-number pattern reflects genetic variation instead of technical variability (e.g., due to FFPE DNA degradation). Moreover, the observation of clonal (‘discrete’) copy-number changes on both Chr.9 and Chr.11 in the IMEAC genome within the same regions of sloping copy number in HGD suggests that the IMEAC ancestor was a subclone of HGD. Remarkably, the IMEAC genome does not show clonal copy-number alterations on 12q that would have been derived from an HGD subclone with varying 12q loss, but contains a high-level amplification spanning *KRAS* on the 12p arm; the amplification was inferred to have originated from the same parental chromosome with sloping copy number variation on the 12q-terminus in HGD. It is tempting to speculate that the *KRAS* amplification had evolved from an unstable Chr12 missing the q-terminus by chromothripsis and subsequent duplications.

**Figure 8:**
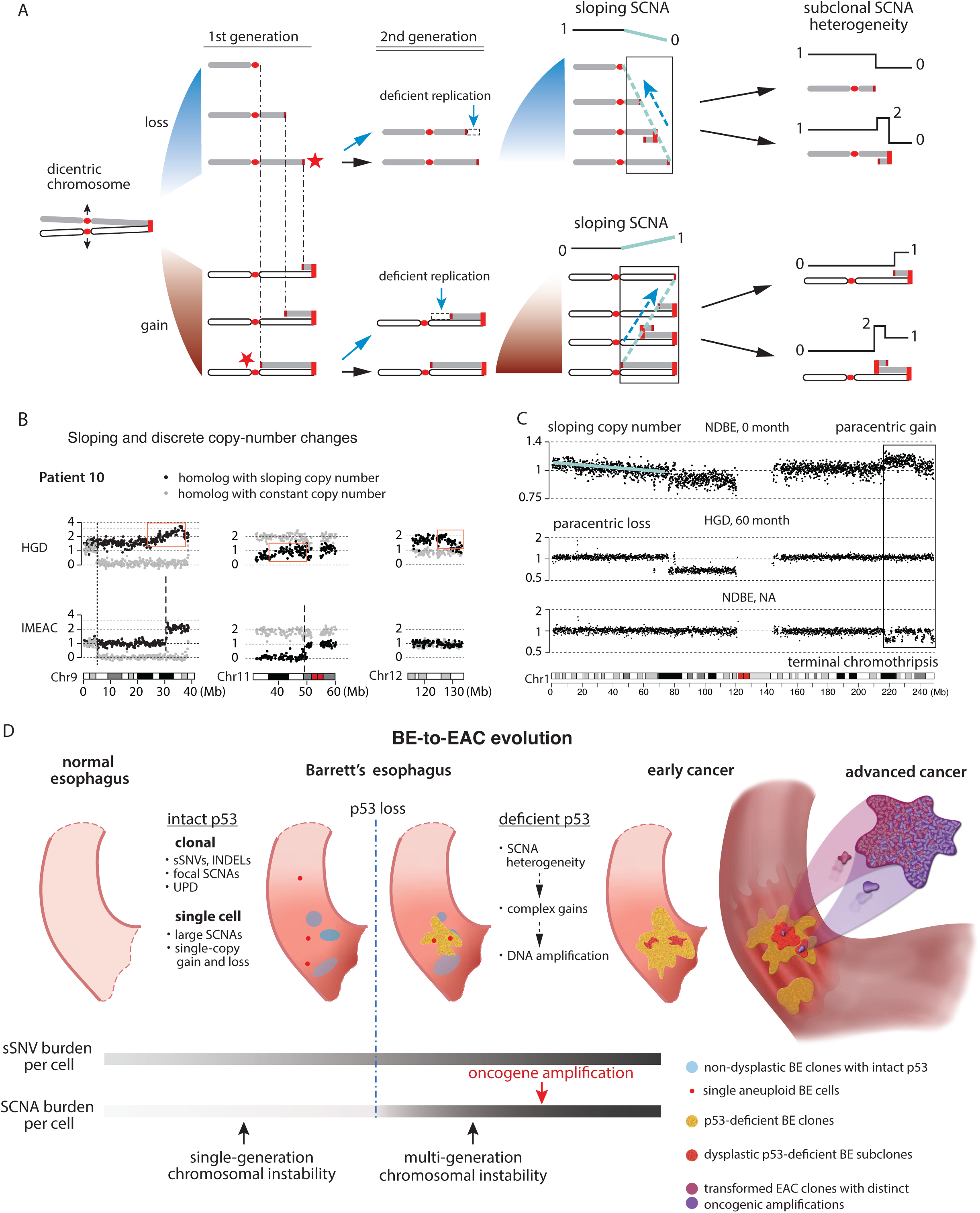
Chromosomal instability creates copy-number heterogeneity prior to copy-number complexity. **A**. Successive BFB cycles can generate progressive DNA losses at the broken ends of chromosomes resulting in a gradual attenuation (sloping) of DNA copy number towards either telomeric (top) or centromeric (bottom) boundaries. Individual broken ends in single cells may acquire terminal duplications that become visible after clonal expansion, but the population average will accrue DNA loss due to deficient DNA replication. **B**. Sloping DNA copy number on Chrs.9, 11, and 12 (black dots) in the HGD sample from Patient 10. The constant DNA copy number of the other homolog is shown in gray. In the regions of sloping copy-number variation on Chrs.9 and 11 in HGD, we observe clonal copy-number changepoints in IMEAC, suggesting clonal expansion of a subclone/single cell in the HGD sample. **C**. BE copy-number evolution revealed in longitudinal BE sequencing data published by Killcoyne et al. (2020). In this patient (Patient 86), the NDBE sample at 0 month displays sloping (1p terminus) and subclonal (1q terminus) copy-number variation. A subsequent HGD lesion (at 60 months) from the same patient shows a (sub)clonal paracentric loss on 1p; another NDBE lesion (timing unspecified) showed chromothripsis at the 1q-terminus in the same region of subclonal copy-number gain in the NDBE lesion at 0 month. Both examples indicate copy-number heterogeneity. See **Extended Data Fig. 10** for additional examples. **D**. Evolutionary dynamics of local sequence changes (single-nucleotide variants, short sequence deletions/duplications) and chromosomal structural aberrations during esophageal cancer evolution. Prior to p53 loss, the suppression of clonal expansion of chromosomal structural alterations implies that only alterations that do not disrupt chromosomal instability (local sequence changes, focal deletions/duplications, or uniparental disomies) are detectable at the clonal level. After p53 loss, there is a rapid increase of SCNA burden per cell that is due to both clonal expansion of ancestral SCNAs and SCNA accumulation during multigenerational evolution of unstable chromosomes, which generates both copy-number heterogeneity and DNA duplications. Although the average mutational burden per cell (of both local and structural alterations) and the total genetic diversity of the tumor clone continue to increase during cell proliferation, the acquisition of cancer drivers can cause clonal dominance or sweep that make minor subclones harder to detect by bulk or even single-cell sequencing. Therefore, analyses of precancer lesions with limited clonal expansion can reveal ancestral genetic heterogeneity that may be undetectable in advanced cancers.

To further explore the possibility that sloping copy-number variation in early-stage BE samples precedes clonal SCNAs in late-stage BE subclones, we analyzed the sequencing data of longitudinal BE samples released in a recent study^55^ (**Extended Data Figure 9A**). We first confirmed the presence of large segmental SCNAs in both non-dysplastic and dysplastic BE samples prior to transformation and the presence of distinct copy-number alterations in aneuploid BE or early cancer clones indicating copy-number evolution (**Extended Data Figure 9B and 10, Extended Data Table 8, Supplementary Data**). The observation of extensive copy-number evolution in longitudinal BE samples provides orthogonal evidence of persistent chromosomal instability in BE cells that complements the observation of widespread copy-number heterogeneity in multifocal BE samples. We further identified sloping copy-number variation in 9 patients. (Due to the limited sequencing depth, this inference was based on total DNA sequence coverage instead of haplotype-specific coverage.) In Patient 86, we observed sloping copy-number variation on the 1q arm in the NDBE sample indicating varying terminal gains (**Figure 8C**, top); the same region shows a clonal terminal retention in a late-stage HGD sample (**Figure 8C**, middle). In contrast to the sloping DNA copy number of 1p, the 1q arm contains a subclonal paracentric gain that may be related to the chromothripsis at the same 1q-terminal region in another NDBE lesion (**Figure 8C**, bottom). Together, the observations in both longitudinal and multifocal BE samples suggest ongoing evolution of unstable BE genomes prior to the emergence of EAC clones. As sloping copy-number variation precedes clonal SCNAs, it may ultimately serve as a prognostic marker of BE progression or ongoing genome instability.

## Discussion

We here studied precancer genome evolution in a unique sample set of incipient esophageal adenocarcinomas and adjacent Barrett’s esophagus lesions by haplotype-specific copy-number analysis. We identified recurrent copy-number evolutionary patterns related to both gross karyotype changes and complex segmental alterations including focal amplifications that indicate continuous genome instability in BE cells.

We find that arm-level copy-number changes often accumulate in episodic bursts and are consistent with the outcome of whole-genome duplication (WGD) and downstream events including multipolar cell division and micronucleation^32, 33^. WGD is frequently followed by extensive chromosome losses, giving rise to highly aneuploid genomes, but can also generate near complete genome duplication. For example, the EAC genome in Patient 7 is a near complete duplication of the LGD2 genome (with odd copy-number states on 4q, 5, and 9q indicating post-WGD losses); the D5 cell in the single-cell collection is close to a complete duplication of the F12 cell (with odd copy-number states on 2p, 9q and post-WGD gains of 17q and 18p). When and how duplicated genomes re-establish stable karyotypes *in vitro* and *in vivo* require further investigation.

We find several patterns of segmental copy-number alterations in BE/EAC genomes that are consistent with an origin from dicentric chromosome breakage and evolution^38^. These include simple segmental copy-number gains and losses consistent with the outcome of a single BFB cycle (**Figure 5**), compound copy-number gains consistent with successive BFB cycles (**Figure 6A-C**), and distinct copy-number alterations to a single parental chromosome in related BE/EAC genomes that are consistent with copy-number variation generated by multigenerational BFB cycles (**Figure 6C-E**). The mechanistic association between BE/EAC genome complexity and BFB cycles is further supported by the presence of regional or arm-level chromothripsis (**Figure 7A,C** and **Extended Data Fig. 8A-F**), interchromosomal translocations (**Figure 6A,B, Extended Data Fig. 8F,G**), and tandem-short-templates rearrangements (**Extended Data Fig. 8D,G**), all of which were previously identified *in vitro*^37, 38^. Finally, the patterns of progressive DNA deletions (**Figure 6D**) and sloping copy-number variation (**Figure 8B,C**) provide strong evidence for ongoing BFB cycles^38^ in BE cells. The sloping copy-number pattern is most simply explained by the under-replication of a broken chromosome over multiple generations that generates a polyclonal mixture of cells with varying DNA losses. This pattern of polyclonal copy-number variation may be regarded as a signature of ongoing or ‘present’ genome instability that precedes clonal SCNAs that indicate ‘past’ genome instability (**Figure 8A**).

We observe nearly ubiquitous bi-allelic *TP53* inactivation preceding the emergence of aneuploid BE cells or BE clones. This result reinforces prior observations in BE cells^50^ or from comparative studies of BEs and late EACs^11, 30, 31, 56^. However, cells with intact p53 do occasionally acquire large copy-number alterations. This is demonstrated by the observation of infrequent arm-level or large segmental SCNAs in single BE cells (**Figure 3**) and even instances of chromothripsis in BE clones (e.g., on Chr9p in Patient 8 BE1-3, Patient 11 LGD, and Patient 6, all samples) inferred to have occurred prior to *TP53* inactivation. In contrast to BE cells with intact p53, the most distinguishing features of p53-null BE cells include (1) massive aneuploidy including whole-genome duplication; and (2) complex segmental gains (with copy-number states above two) that require multiple generations of chromosome breakage and recombination. This observation suggests that the dominant tumor suppressive mechanism of p53 may be the suppression of cell proliferation after chromosome missegregation^44^.

The abrogation of p53-dependent cell cycle arrest after chromosome missegregation has two implications (**Figure 8D**). First, arm-level or large segmental SCNAs generated by chromosome missegregation events are more likely to undergo clonal expansion and become visible at the clonal level. Second, and more importantly, it allows single cell division errors such as whole-genome duplication or chromosome bridge formation to precipitate multigenerational instability that both generates copy-number heterogeneity and fuels the acquisition of oncogenic amplifications. Therefore, even without an apparent increase in the rate of events that generate unstable chromosomes, p53 loss marks the onset of rapid accumulation of copy-number heterogeneity and complexity that contrasts with continuous SNV accumulation. This explains the significant differences between SCNAs in ageing esophagus or BEs with intact p53 and in BEs with deficient p53. Interestingly, we observed a novel pattern of copy-number variation in BE cells with intact p53 reflecting uniparental disomy (UPD) alterations with varying boundaries (**Figure 3E**). How large segmental UPDs arise in mammalian cells is unknown. The similarity of progressive DNA breakpoints in varying UPDs to those in progressive DNA losses (**Figure 6D**) suggests that these two patterns may reflect different DNA repair outcomes of broken chromosomes generated by successive BFB cycles (**Extended Data Fig. 4**). If this model were true, it further implies that cells with intact p53 do tolerate certain types of chromosomal instability but raises the question of how p53 or other selection factors impact the rearrangement outcomes of such instability.

The early onset of genome instability during BE progression revealed in our analysis challenges the prevailing view that chromosomal aberrations are exclusive to advanced cancers or only arise late during tumor development. Analyses of advanced tumors by either bulk^5^ or single-cell^57^ sequencing usually reveal only truncal or late subclonal alterations, indicating relatively late divergence of different cancer subclones. As late-stage cancers are often dominated by the most aggressive clones, analyses of late-stage cancers cannot reveal copy-number heterogeneity in single cells prior to transformation. By contrast, genetic diversity is more visible in precancerous lesions due to the lack of dominant clones. This explains the observation of significant copy-number differences in multifocal BE clones (**Figure 2**), copy-number evolution in longitudinal BE samples (**Extended Data Fig. 9,10**), and sloping copy-number variation in single BE lesions (**Figure 8B,C**). Moreover, the generation of complex copy-number gains, including focal amplifications, necessitates multigenerational chromosomal instability that invariably creates copy-number heterogeneity (**Figures 3, 6, 7**). Therefore, complex DNA gains in EACs or dysplastic BEs can be regarded as a signature of ‘past’ chromosomal instability in their ancestor cells.

Oncogenic amplifications are a hallmark of advanced EACs. Our analyses demonstrate that these events are frequently present in both early EACs and dysplastic BEs with deficient p53 (**Figures 2** and **3**). We further identified distinct oncogenic amplifications in different dysplastic BEs or early EACs from the same patient (**Figure 2** and **7C**), some of which were associated with independently transformed EAC foci. As independent EAC clones may grow into each other to form a single tumor mass or seed different metastatic lesions, both intratumor and primary/metastasis oncogenic amplification heterogeneity^58^ may be the inherent outcome of chromosomal instability after p53 loss that could have been initiated in precancer BE cells and persist after transformation.

Our model of chromosomal-instability driven copy-number evolution makes several predictions. First, segmental copy-number complexity at the clonal level is preceded by copy-number heterogeneity at the single cell level. This is demonstrated in our study (**Figure 3** and **8**) but should be further tested by single-cell DNA sequencing of precancerous or ageing tissues. Second, p53 loss enables the accumulation of copy-number heterogeneity in precancer lesions that may differ from late-stage cancers due to the lack of clonal sweep. This prediction can be tested in other cancers with early p53 inactivation and precursor conditions, including serous ovarian cancers^16^, basal breast cancers, uterine serous endometrial cancers, pancreatic cancers^59^, and colitis-associated colorectal cancers^15^. Finally, our analysis of SCNAs in BE/EAC genomes suggests a mechanism-based classification of copy-number patterns. Extending this analysis to cancers both with and without *TP53* inactivation will generate new knowledge of tumor evolution dynamics with both diagnostic and therapeutic implications.

## Data availability

All sequencing data generated in the current study were uploaded to Genotypes and Phenotypes (dbGaP) (accession <u>phs002706</u>) with controlled access according to the Protocol approved by the Institutional Review Board of the Brigham and Women’s Hospital. Third-party data that were re-analyzed were obtained from the European Genome-phenome Archive (EGA) through data access agreement approved by the International Cancer Genome Consortium. Processed DNA copy number data and compiled copy-number plots are available at https://github.com/chunyangbao/NG_ESAD75.

## Code availability

Usage of published or public bioinformatic packages is stated in Methods with references to either the publications or the repositories of the software packages. All the algorithms and bioinformatic pipelines implemented in this study are described in Methods; scripts and codes are uploaded to https://github.com/chunyangbao/NG_ESAD75.

## Supporting information

Supplementary Information on the computational method and validation

Methods and Captions for Supplementary Figures and Tables

## Downloadable Supplementary Data

Haplotype-resolved DNA copy number of bulk BE/EAC samples

Haplotype-resolved DNA copy number of single BE cells from HGD brushing

Total DNA copy number of longitudinal BE sequencing data (re-analysis)

## Acknowledgements

We would like to thank all patients who were willing to contribute samples to this study, the DFCI Center for Cancer Genomics and the Genomics Platform at the Broad Institute of MIT and Harvard for assistance with library preparation and sequencing, and Dr. David Pellman for a critical reading of the manuscript.

## Author contributions

Conception of study: MDS, AJB, CB, CZZ; Patient selection and clinical data collection: KW, JMD, KKM, MDS, JK; Histologic review: AA, RO, MDS; Sequencing experimental design and data generation: MDS, LS; Fluorescence in-situ hybridization analysis: HB, MW, YI; Bioinformatic analysis: CB, MDS, CZZ with help from RT, GB, CS, GG; Manuscript preparation: CZZ, CB, MDS, AJB; Manuscript review: KKW, YI

---

**FIGURE S1.**
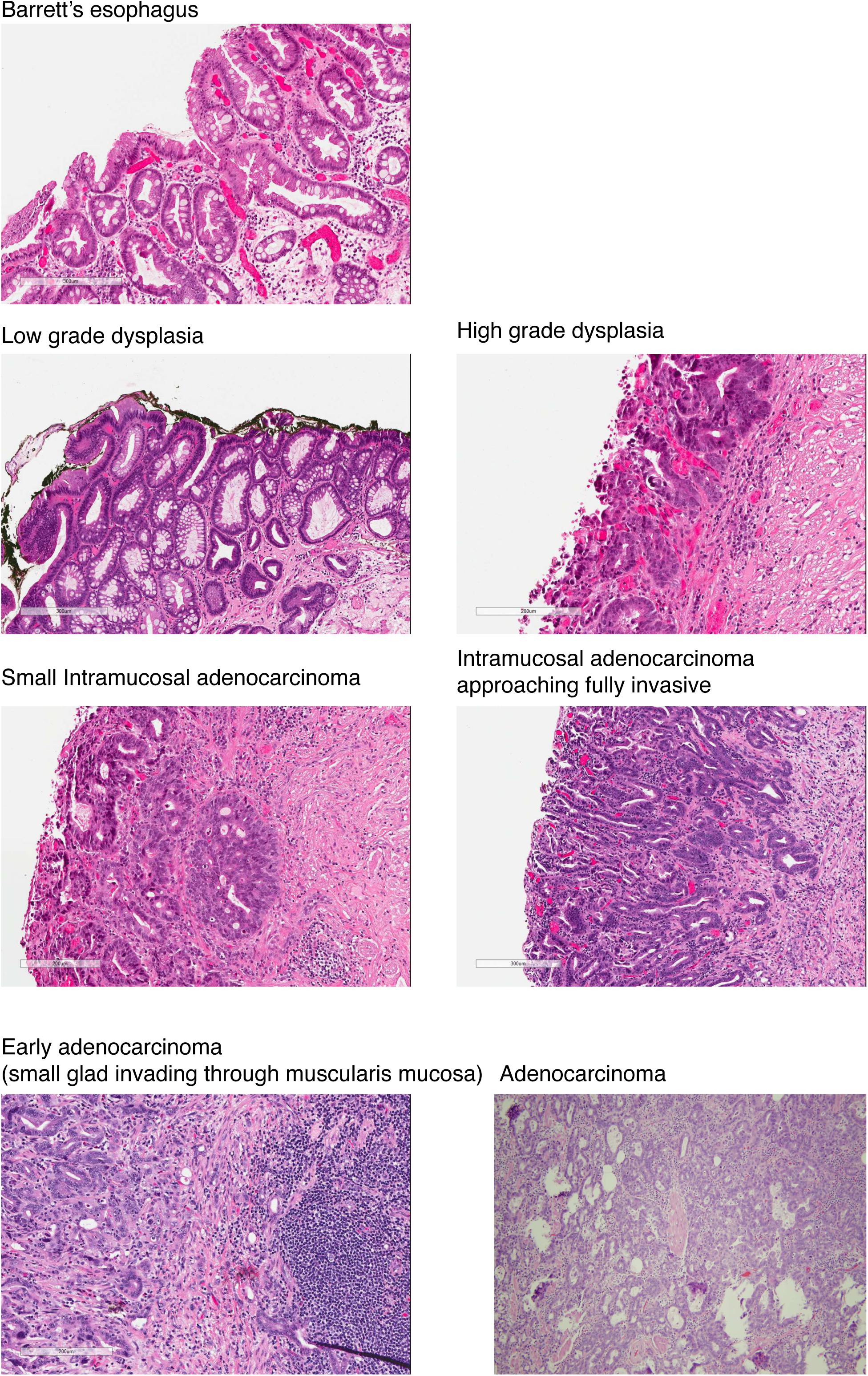

**FIGURE S2.**
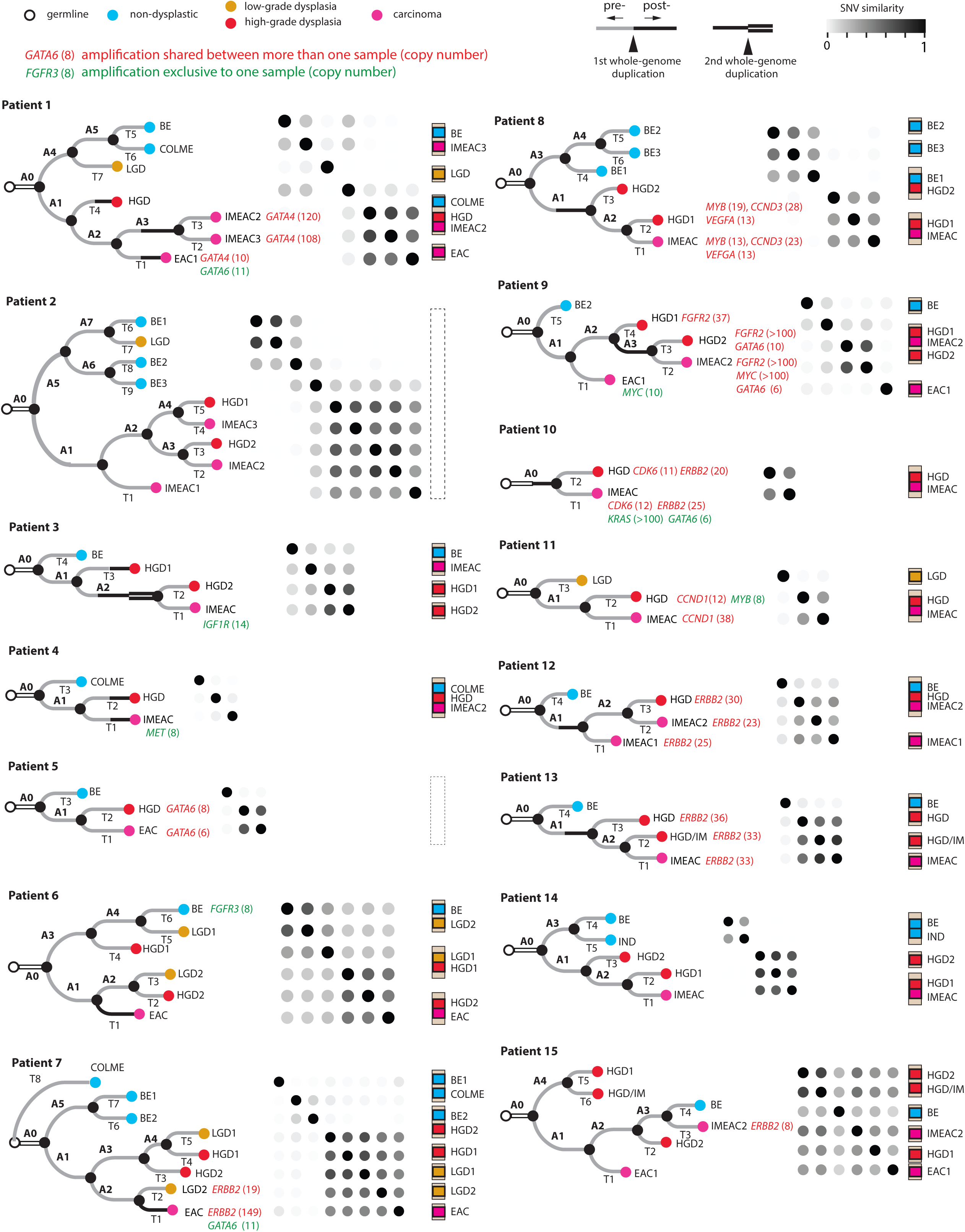

**FIGURE S3.**
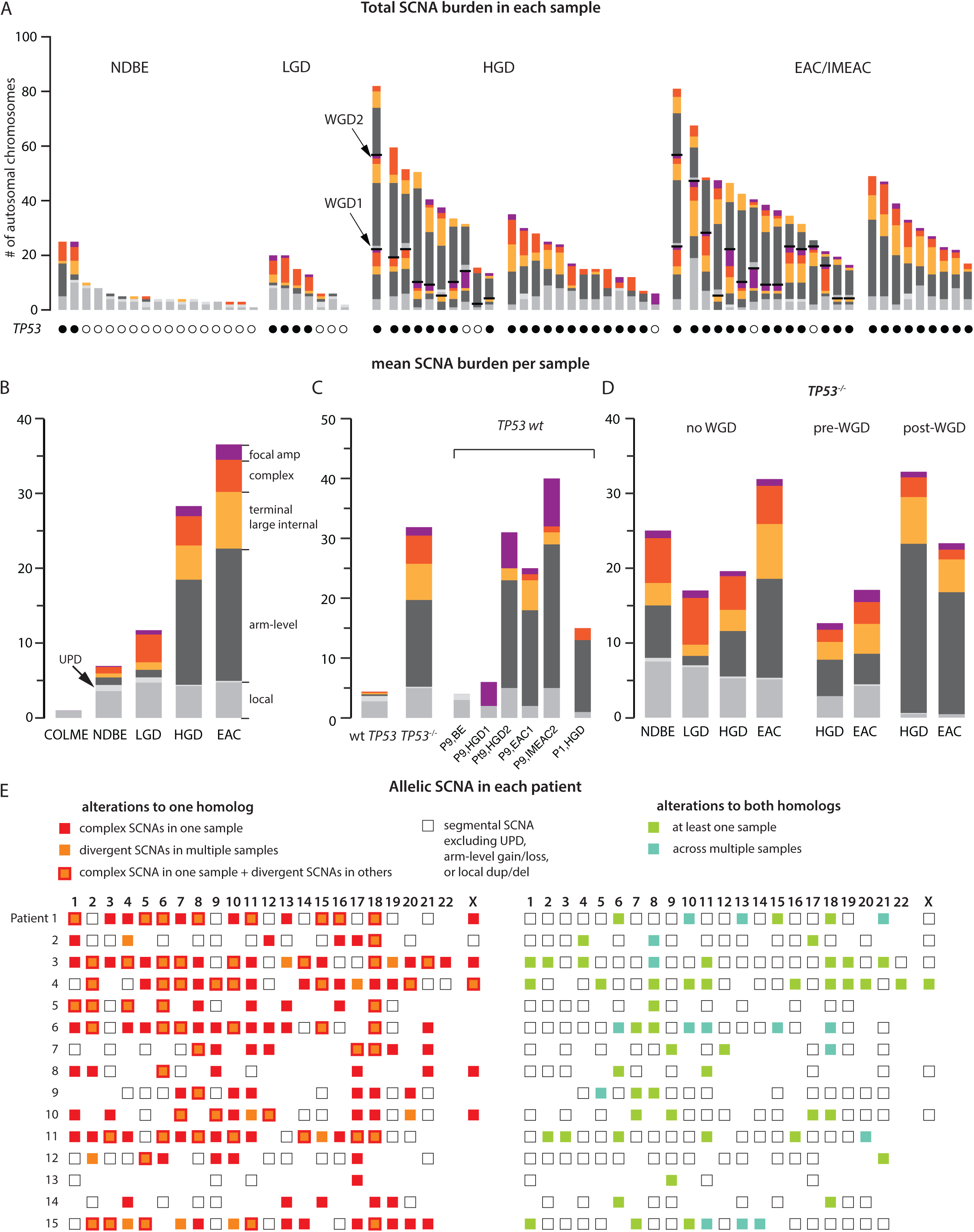

**FIGURE S4.**
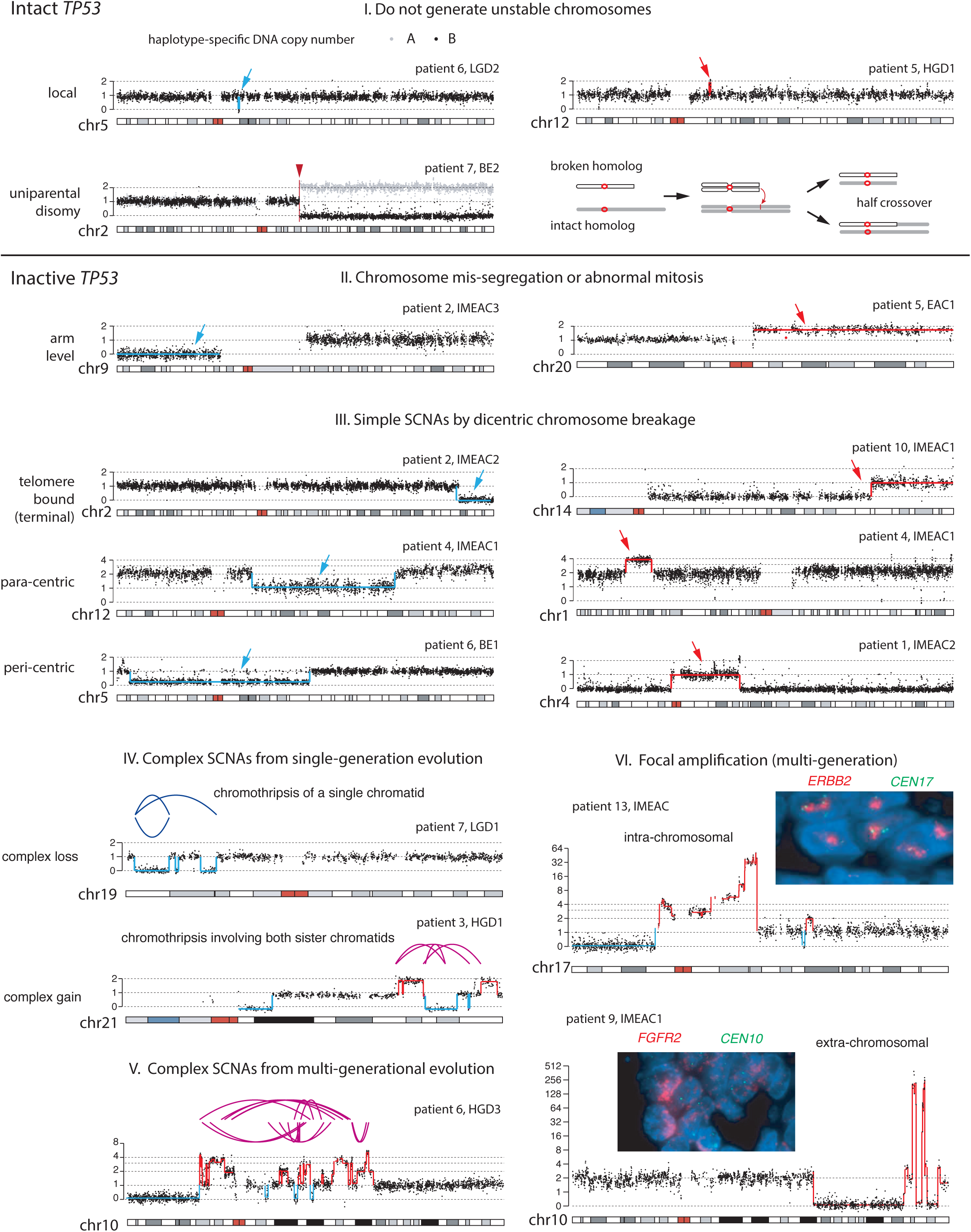

**FIGURE S5.**
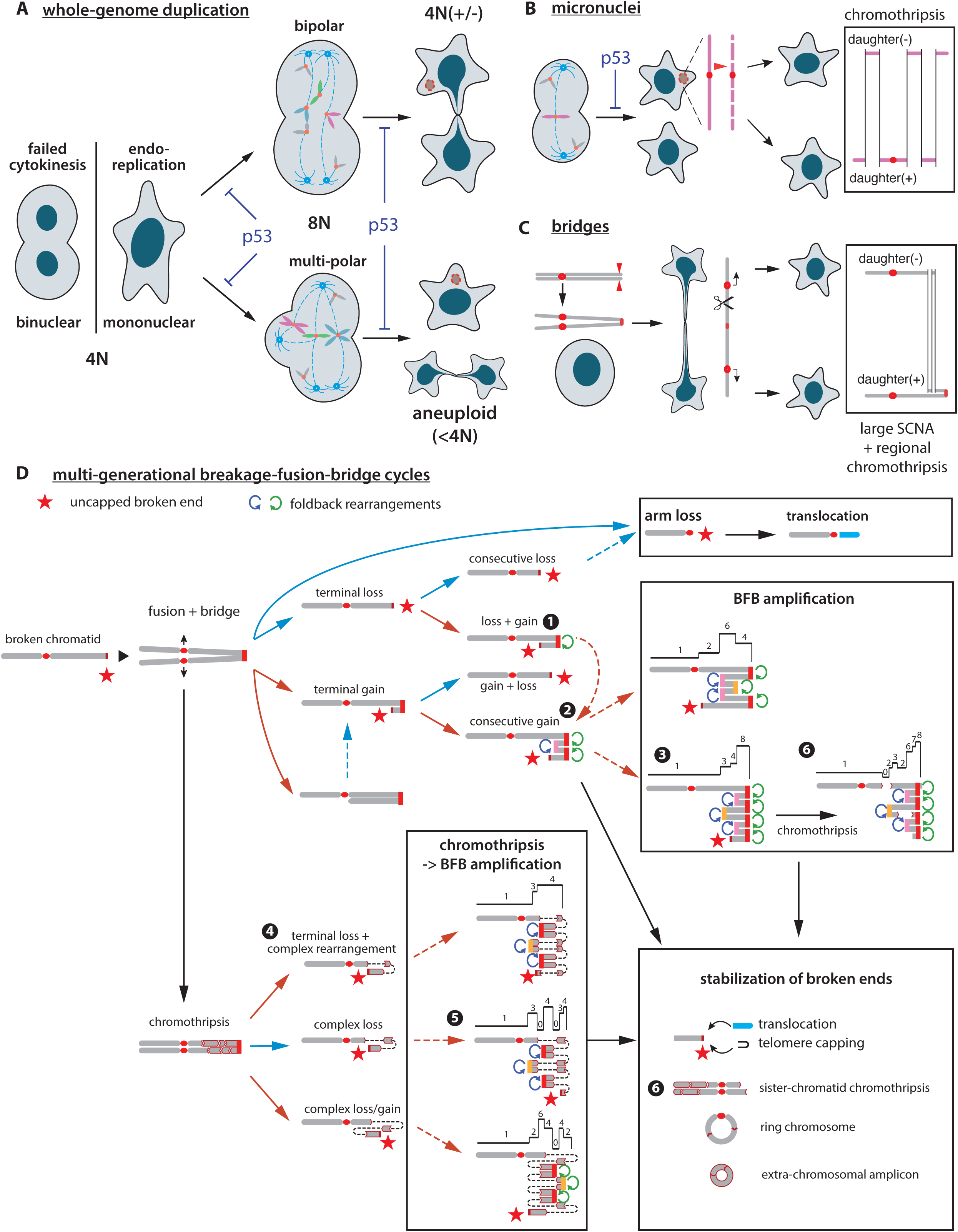

**FIGURE S6.**
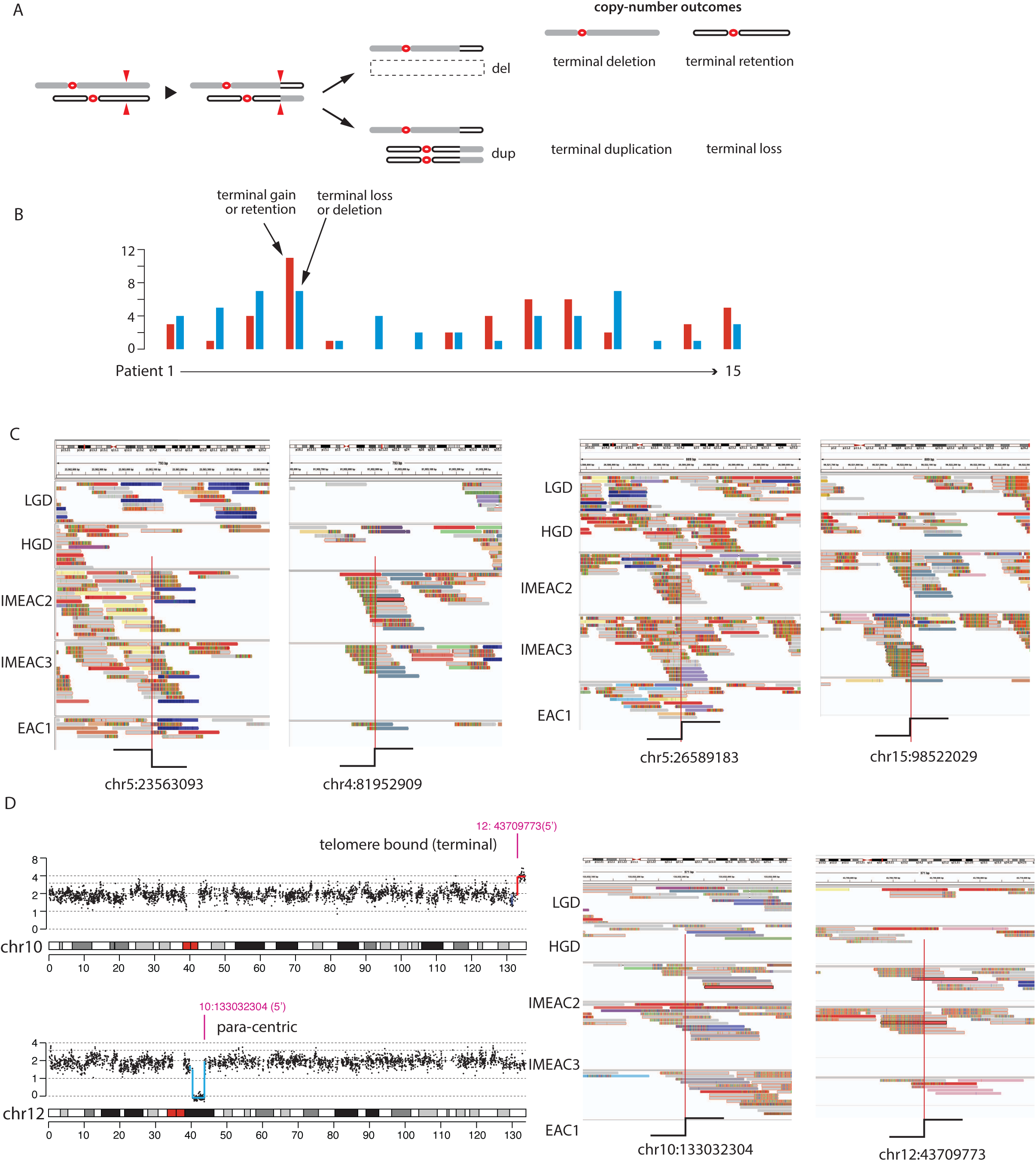

**FIGURE S7.**
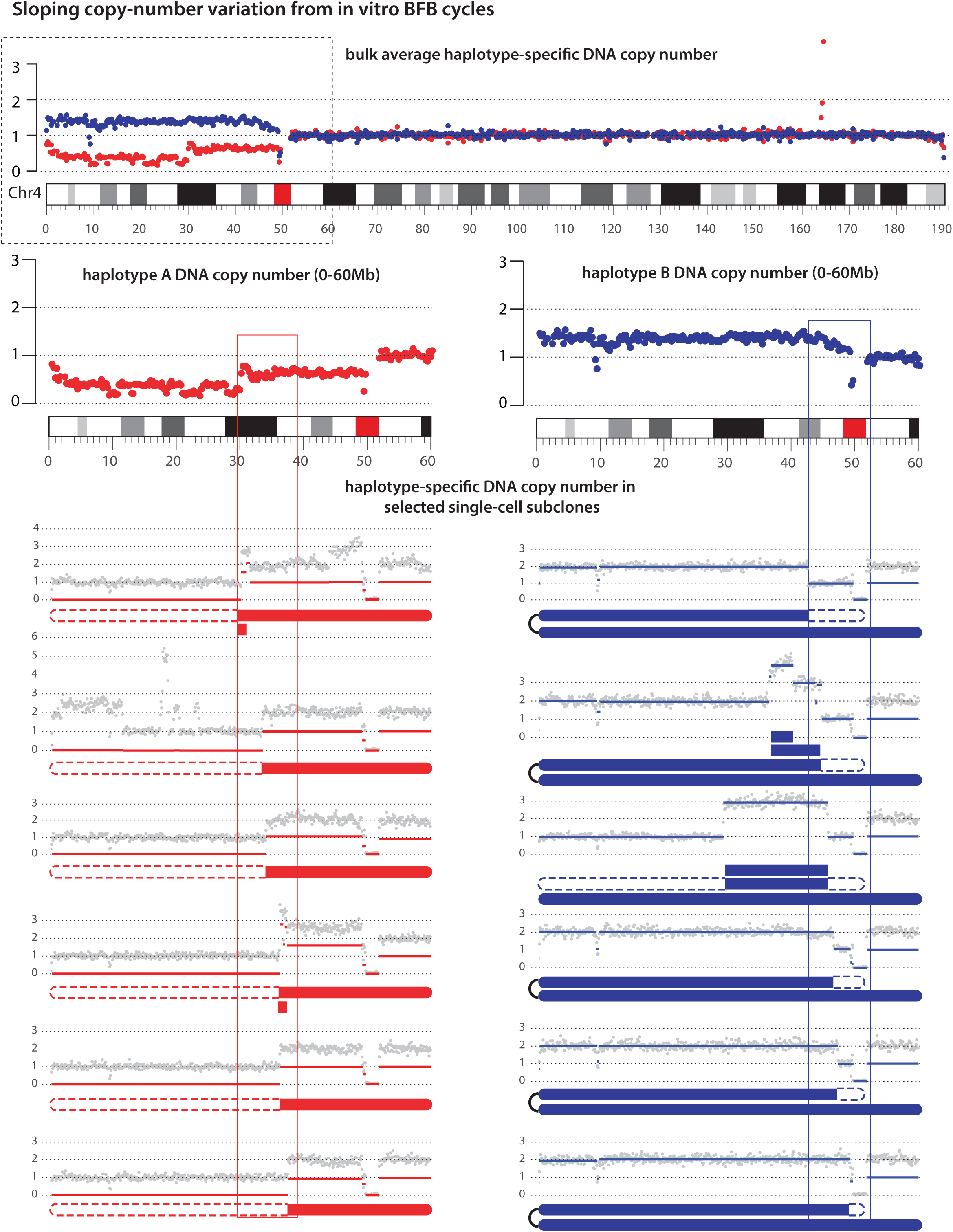

**FIGURE S8.**
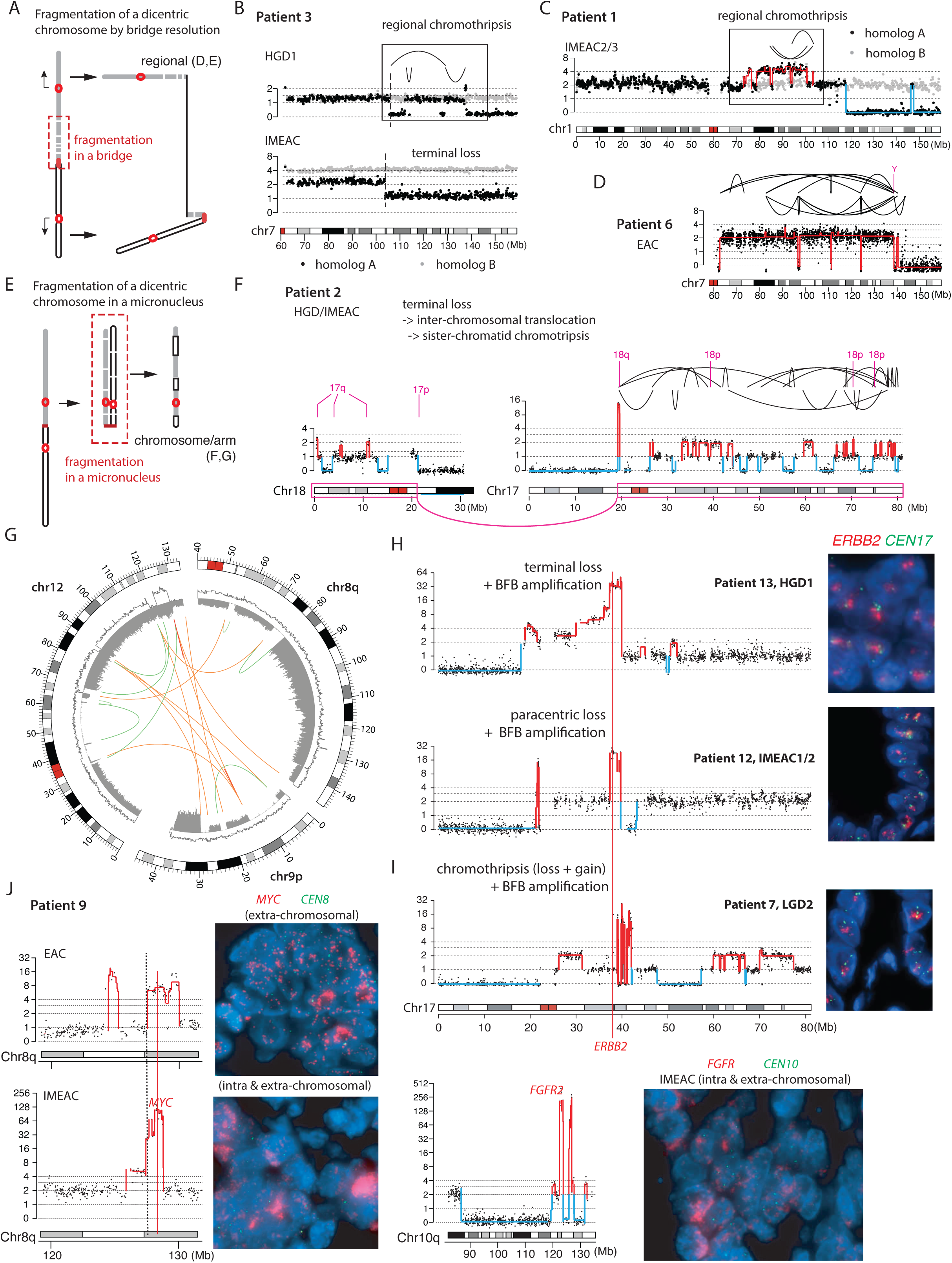

**FIGURE S9.**
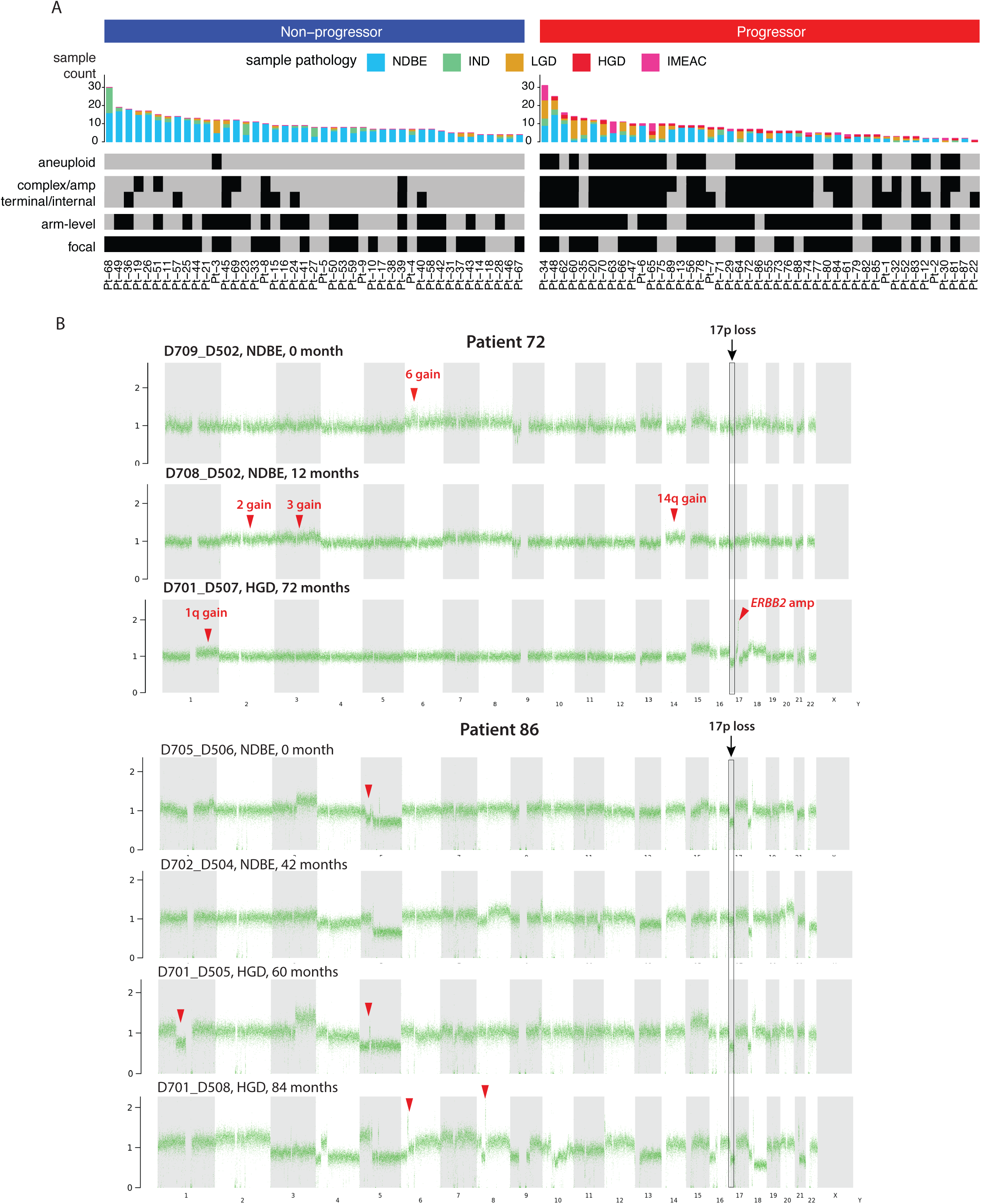

**FIGURE S10.**
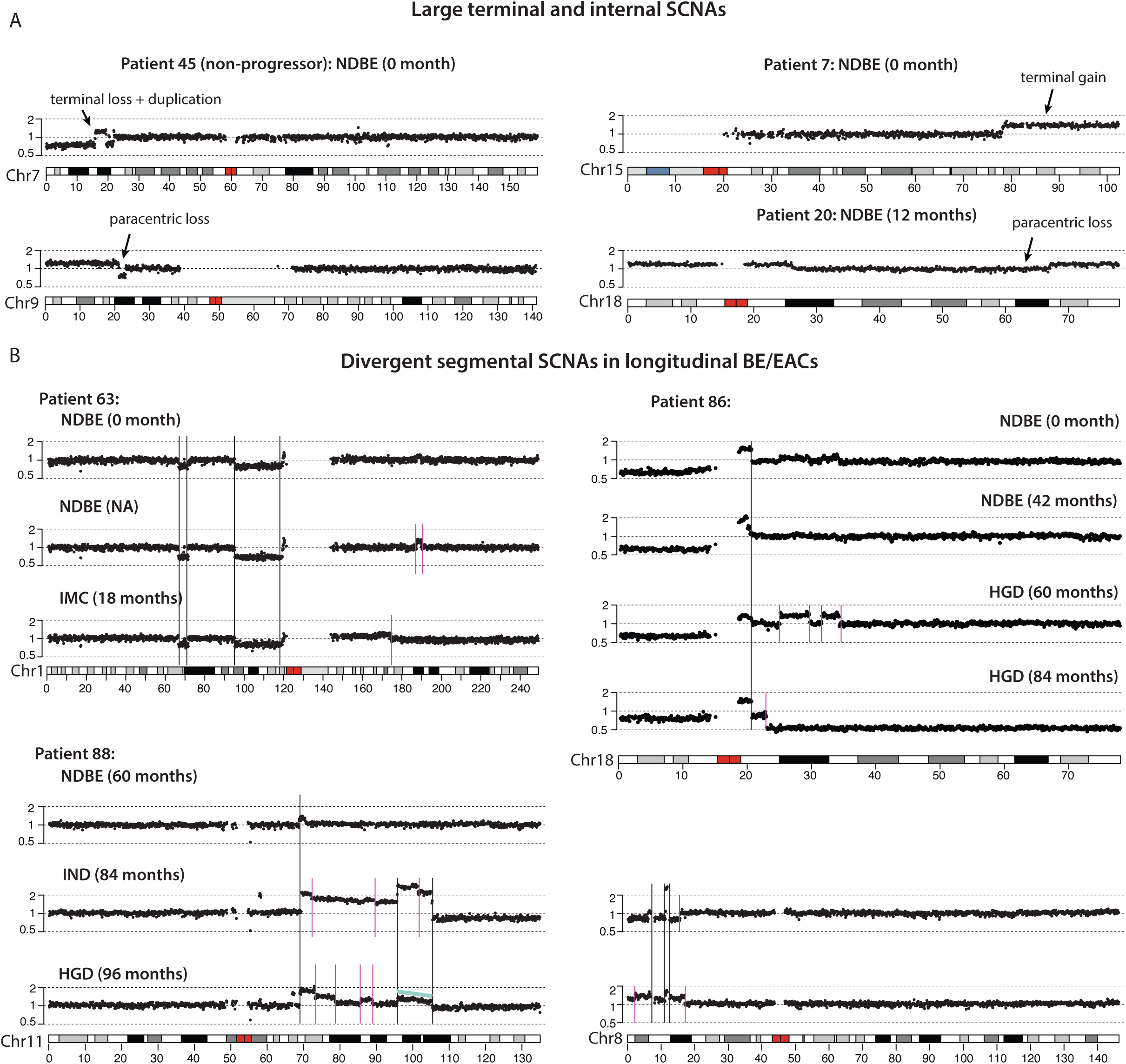

**Additional Figure 1.**
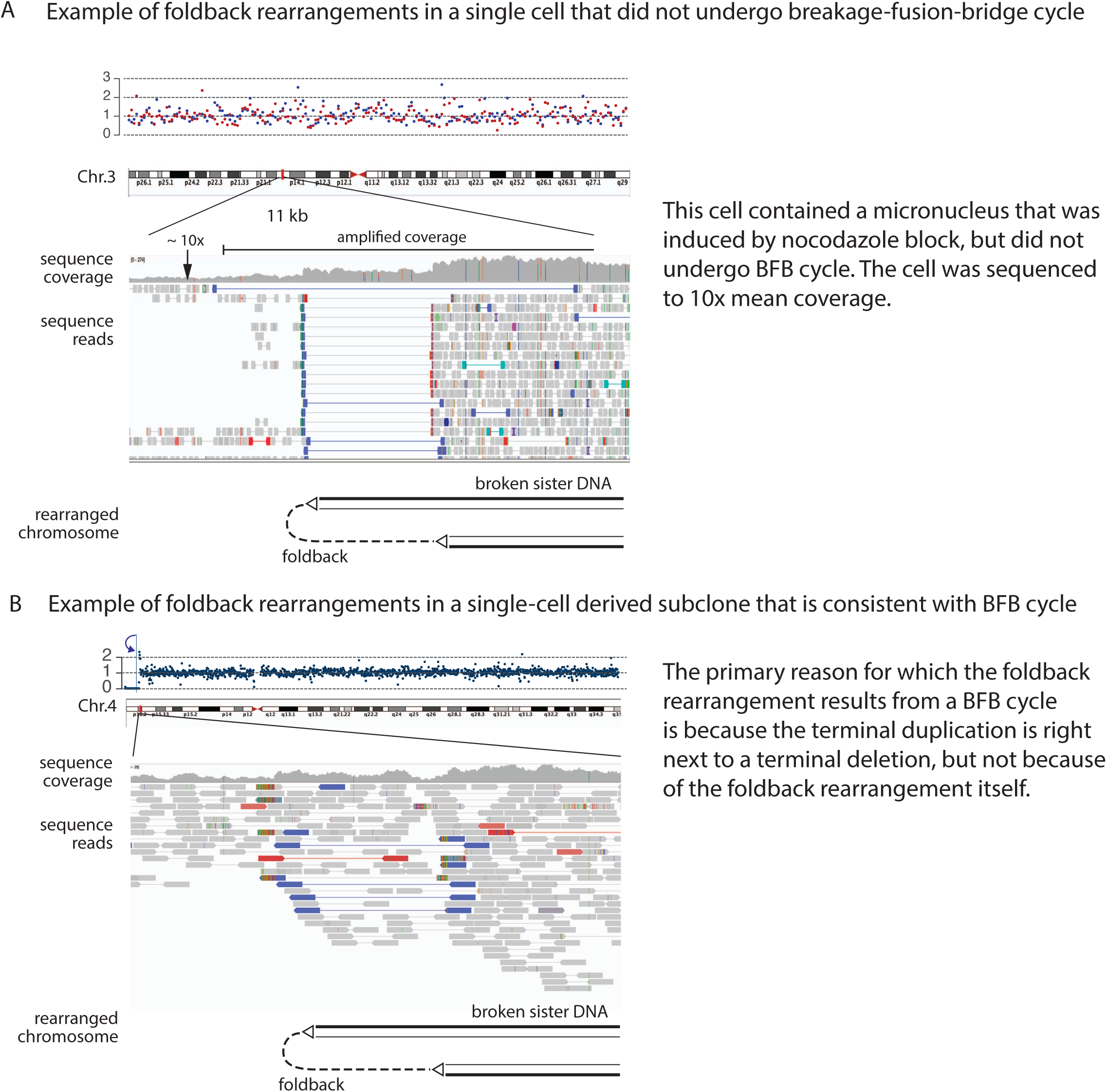

**Additional Figure 2.**
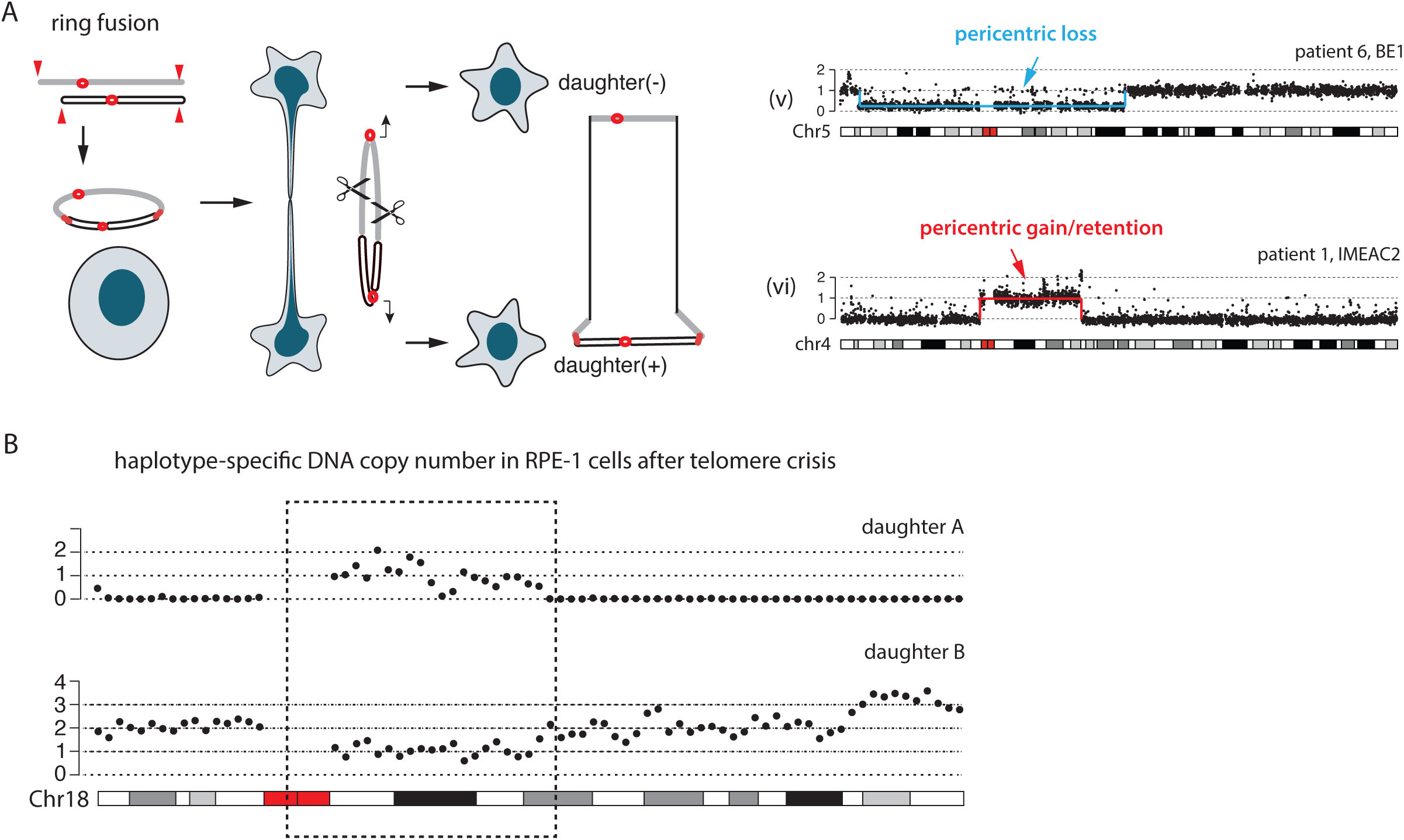

