## Supplementary Information on the computational method and validation for "Genomic signatures of past and present chromosomal instability in the evolution of Barrett’s esophagus to esophageal adenocarcinoma"

#### This PDF file includes:

Supplementary text
Figs. S1 to S11
Captions for Dataset S1 to S2
References for SI reference citations

#### Other supplementary materials for this manuscript include the following:

Datasets S1 to S2

### Supporting Information Text

### Analysis of somatic genome evolution in paired cancer and precancer samples

**General goals of the analysis.** The genomic analysis of paired BE and esophageal cancer lesions, or more generally of multi-regional or longitudinal cancer samples, needs to accomplish three goals. First, we need to identify somatic alterations in these lesions. Second, we want to determine the phylogenetic relationship between these lesions (or different clones in these lesions) based on genetic alterations that are either shared or related between these lesions. (The difference between "shared" and "related" alterations will be discussed later). Finally, based on the phylogenetic relationship, we want to infer the evolutionary sequences of genetic changes, including their timing relative to disease progression. Below we discuss the bioinformatic and computational challenges related to each goal and strategies to overcome these challenges. We use examples from data in the current study to illustrate these challenges.

**Technical challenges in the detection of genetic variants from FFPE DNA sequencing.** The detection of genetic variants in FFPE DNA libraries is confounded by sequence artifacts generated during DNA extraction and library construction due to DNA degradation. These technical artifacts include: (1) base substitution errors introduced during amplification of damaged DNA (including abasic or damaged bases) [1], which introduce false single-nucleotide (SNV) or insertion/deletion (INDEL) variants; (2) chimeric DNA (often consisting of sequences from opposite strands) generated by template-switching events during polymerase extension from nicked or single-stranded DNA templates [2], which may be falsely identified as sequence rearrangements; (3) uneven sequence coverage due to varying DNA recovery (degradation, chromatinization etc.), which can be falsely identified as copy-number alterations. Uneven sequence coverage also contributes to false negative detection of local sequence alterations including single-nucleotide variants, insertions/deletions, and rearrangements.

Besides sequencing artifacts, the limited amount of input DNA from micro-dissected tissue samples also results in low DNA library complexity that also causes low variant detection sensitivity. Although the technical limitations of FFPE libraries can be ameliorated by modified experimental protocols of DNA fixation or extraction, they cannot be eliminated and will pose challenges to both variant detection and downstream analysis.

**Analytical strategies to cope with technical limitations of FFPE sequencing.** For sequence alterations including single-nucleotide variants (SNV), insertion/deletion (INDEL) events, and rearrangements, the specificity of their detection can be improved either by setting a more stringent threshold of variant read support or by using independent replicates (i.e., separate tissue specimens from the same tumor): The requirement for more variant read support may remove artificial sequence changes that are generated only on a small fraction of input DNA molecules; however, this requirement also leads to lower detection sensitivity due to uneven sequence coverage or low sequencing depth of tumor DNA.

In contrast to requiring more variant read support or increasing the mean sequencing depth, using biological replicates is a more efficient strategy for controlling sequence artifacts in our sample cohort as we are most interested in alterations that are present in more than one BE or EAC lesions and therefore informative about their phylogenetic relationship. We have therefore performed joint detection of both local variants (SNVs and INDELs) and rearrangements from all BE/EAC lesions from each patient instead of the standard tumor/normal pair analysis. (See **Joint variant detection by HaplotypeCaller** and **Somatic rearrangement detection** sections in **Online Methods** for more details.)

Although the joint variant detection strategy can suppress false positive variants in individual samples and achieve better sensitivity for shared variants, it has two limitations. First, it cannot distinguish between false variants and true private mutations that are only present in a single sample. It is therefore impossible to perform comprehensive genetic variant discovery in individual BE or EAC lesions or benchmark detection accuracy. Second, joint variant detection cannot eliminate false negative variant detection (including genotyping of true variants) in individual BE/EAC lesions as the detection sensitivity depends on the quality of input DNA, the depth of sequencing, as well as tumor purity.

As will be discussed in the next two subsections, the inadequate detection of SNVs, INDELs, and rearrangements can confound tumor phylogenetic inference based on these variants. By contrast, one can achieve more robust phylogenetic inference based on somatic copy-number alterations (SCNA) due to the better detection accuracy and the ability to phase SCNAs to parental chromosomes.

**Computational and bioinformatic challenges related to tumor phylogenetic inference.** The general rationale for tumor phylogenetic inference is that tumor clones with a more recent common ancestor should share more genetic alterations than tumor clones with an earlier ancestor. Therefore, the phylogenetic distance ("relatedness") between two tumor genomes is inversely correlated with the number of shared genetic alterations. A simple and commonly used approach is to estimate phylogenetic distance based on the number of genetic variants, including SNVs/INDELs, rearrangements, and SCNAs, that are shared between two tumor samples. However, this approach has several limitations.

First, because early/truncal mutations may be lost in progeny clones due to downstream **DNA deletions or loss-of-heterozygosity alterations**, the number of genetic variants shared between two tumor clones is not equivalent to the number of genetic alterations in their common ancestor. Consider an example of three tumor clones **A**, **B**, and **C**: **A** descended from the common ancestor, whereas **B** and **C** are expanded from sibling cells with reciprocal DNA gain and loss (such as shown in Fig. 6C and Fig. 7B). Although **B** and **C** share a more recent ancestor than A, there are actually more mutations shared between **A** and **B** or between **A** and **C** than between **B** and **C**, because **B** and **C** each have lost mutually exclusive subsets of ancestral mutations on the broken chromosome. This example offers a possible explanation for the discordance between phylogenetic trees determined from point mutations and from copy-number changes (see [3] for an example).

*Note: We use this example only to illustrate the ambiguity of shared genetic variants as a measure of genetic relatedness when used for clustering-based phylogenetic inference. In principle, one can still use shared genetic variants to rank different phylogenetic trees (e.g., using a maximum likelihood approach) as long as variants in deleted regions are excluded from the calculation. However, the latter requires phasing of somatic variants that is usually not possible with shotgun sequencing data.*

The second limitation of using the number of shared variants for phylogenetic relatedness is due to the presence of different subclones in a single tumor sample. Mutations detected in a polyclonal tumor sample may include mutually exclusive variants in different subclones. Therefore, the number of mutations shared between two tumor samples may not correspond to the number of mutations shared between specific tumor clones in each sample. Consider the following example: tumor sample 1 consists of 40% of tumor clone **A**, 10% of tumor clone **B**, and 50% stroma, and tumor sample 2 consists of 30% of tumor clone **B**, 5% of tumor clone **C**, and 65% stroma. The number of mutations shared between the two samples has no clear relationship to the genetic similarity between the tumor clones. To overcome this ambiguity, one should only consider mutations in the dominant subclone in each sample, i.e., **A** in tumor sample 1 and **B** in tumor sample 2. However, such analysis requires accurate mutant allele fraction calculation and subclonal inference that cannot be achieved with standard (30×) whole-genome sequencing.

Besides these two challenges, tumor phylogenetic inference is further confounded by false negative variant detection. Consider three tumor clones **A**, **B**, and **C** with 10,000 mutations in their common ancestor and an additional 1,000 mutations in the common ancestor of **B** and **C** (i.e., **B** and **C** diverged more recently from each other than from **A**). Assuming a false negative detection rate of 10%, we expect to detect $10,000 \times (1 - 0.1) = 9,000$ ancestral mutations in each clone, $10,000 \times 0.9^2 = 8,100$ mutations shared in two clones, and $10,000 \times 0.9^3 = 7,290$ mutations shared by all three clones. For the additional 1,000 mutations shared between **B** and **C**, we expect to detect $1,000 \times 0.9^2 = 810$ mutations in both. Adding these together, we expect to observe 7,290 truncal mutations in all three clones (not informative about their phylogeny), 810 ancestral mutations shared between **A** and **B**, and 810 + 810 = 1620 mutations shared between **B** and **C**. Therefore, the additional 1,000 mutations shared by clone **B** and clone **C** is diluted by ancestral mutations that are missed in **B** or **C**, which appear as mutations shared by either **B** or **C** with **A**. If sample **C** has a lower detection sensitivity of 0.5 (e.g., due to low tumor cell fraction), then the observed number of mutations shared by **B** and **C** is approximately

$$\underbrace{11,000 \times 0.9 \times 0.5}_{\text{detected mutations shared by } \mathbf{B} \text{ and } \mathbf{C}} - \underbrace{10,000 \times 0.9^2 \times 0.5}_{\text{detected mutations shared by } \mathbf{A}, \mathbf{B}, \text{ and } \mathbf{C}} = 900.$$

This number is comparable to the number of detected mutations shared by A and B (= 810). This calculation indicates that the number of shared mutations detected in tumor samples can be severely confounded by false negative detection of ancestral mutations.

As tumor samples (including many samples in our cohort) often contain less than 50% tumor cells (causing false negative detection of ancestral mutations) but frequently consist of more than one subclones (causing false assignment of mutations to tumor clones), we expect significant uncertainty in the phylogenetic distance estimated from the number of shared point mutations. This uncertainty is further aggravated by the prevalence of DNA losses in aneuploid BE and EAC genomes (causing irreversible loss of ancestral mutations), the presence of whole-genome duplication (further decreasing the detection sensitivity of post WGD mutations), the high frequency of base substitution errors (causing false positive variant detection), and sequence coverage non-uniformity due to FFPE DNA degradation (causing false negative variant detection). Because of these limitations, we seek an alternative strategy of phylogenetic inference based on somatic copy-number alterations.

**Advantages of SCNA-based phylogenetic inference.** The first advantage of SCNA-based phylogenetic inference is their better detection accuracy from FFPE sequencing data than local sequence variants (SNVs, INDELs, and rearrangements). As discussed previously, both the non-uniformity of sequence coverage and the limited sequencing depth of FFPE DNA result in lower detection sensitivity of local sequence variants. As SCNAs affect large segments (including whole-chromosome arms) and alter both the total read depth and the allelic ratio between parental chromosomes, SCNA detection does not require deep sequencing and is insensitive to sequence dropout. Moreover, as individual SCNAs usually affect one of two parental chromosomes (homologs), they can be distinguished from sequence coverage non-uniformity due to technical artifacts (e.g., FFPE DNA degradation) that affect both homologs equally. By contrast, there is no strategy to "correct" false SNVs, INDELs, or rearrangements due to sequence artifacts in FFPE libraries (causing false positive detection).

The ability to detect SCNAs based on allelic coverage not only ensures the accuracy of SCNA detection but further enables phasing of SCNAs to parental chromosomes. By contrast, the haplotype phase of SNVs or rearrangements is generally not available from shotgun sequencing. The phasing of SCNAs to parental chromosomes enables the identification of ancestral SCNAs that are lost due to downstream DNA deletions when the ancestral SCNA and the downstream deletion are phased to the same homolog. This is not feasible for SNVs or rearrangements as one cannot be certain whether these variants were present on the deleted homolog or the intact homolog.

Finally, the phasing information of SCNAs enables us to draw evolutionary inferences about SCNAs or SCNA breakpoints in related BE/EAC samples. When SCNAs or SCNA breakpoints are phased to different parental chromosomes, they arise from independent alterations regardless of the distance between SCNA breakpoints. By contrast, when SCNAs/breakpoints are phased to the same parental chromosome, they could have descended from a single ancestral unstable chromosome with divergent secondary events. This capability enables us to identify mechanistically related chromosomal breakpoints, including reciprocal DNA retention/loss (**Fig. 6C** and **Fig. 7B**), progressive deletions (**Fig. 6D**), and other divergent outcomes of breakage-fusion-bridge cycles (**Fig. 7A** and **7C**).

We note that whole-chromosome or arm-level SCNAs are less informative than segmental SCNAs. This is because segmental SCNAs are non-reiterative (i.e., arising only once) but gain or loss of a chromosome or chromosome arm can arise multiple times independently. Therefore, SCNA-based phylogenetic inference is best suited for tumors with high segmental SCNA burdens. However, this strategy does not require a large number of segmental SCNAs. This is because SCNAs breakpoints are relatively rare (10-100 in BE/EAC clones) and shared SCNA breakpoints impose a strong constraint on the phylogenetic relationship between different tumor clones. The specificity of SCNA breakpoints as lineage markers contrasts with the ambiguity of SNVs due to DNA losses or false negative detection. Using haplotype-specific breakpoints for tumor phylogenetic inference is conceptually similar to using rearrangement breakpoints to determine the phylogeny of animal genomes [4]. A key difference between the two is that the analysis of organismal genome evolution only needs to consider breakpoints in a haploid genome ('gametes'), whereas the analysis of somatic genome evolution needs to consider breakpoints on each parental chromosome separately (i.e., haplotype-specific breakpoints) as the two homologs evolve independently.

### Technical analysis of haplotype-specific copy-number calculation

The haplotype-specific coverage or allelic depth is calculated as the total sequence coverage $R(x)$ multiplied by the average allelic fraction of each parental homolog $F_A(x)$ and $F_B(x)$, where $x$ is the genomic coordinate. For 10× mean sequencing coverage, the average number of reads/fragments in 10kb intervals is

$$10 \times 10^4/\text{read length} = 300 - 500.$$

The standard deviation of read count is approximately proportional to the square root of the read count; therefore, the statistical error of normalized coverage $\delta R(x)/\overline{R(x)}$ is approximately $1/\sqrt{300} \approx 0.06$. This estimation implies that 10× mean sequencing depth is sufficient for the calculation of average sequence coverage $R(x)$ in 10kb or bigger intervals.

In **Online Methods**, we showed that the statistical error of the phased average allelic fraction is approximately

$$\sqrt{\frac{f(1-f)}{n}\left\langle\frac{1}{D}\right\rangle},$$

where $f$ is the haplotype fraction, $n$ is the number of phased variants, and $\langle 1/D\rangle$ is the mean of the inverse of sequence coverage at the phased variant sites. Both $n$ and $D$ depends on the mean sequence coverage. We therefore first assessed the dependence of allelic depths on the sequencing depth.

**Dependence on the sequencing depth.** To determine the minimum sequencing depth required for the accurate calculation of allelic depths, we calculated allelic depths from down-sampled allelic coverage in several BE/EAC samples and compared the results calculated from down-sampled coverage with those derived from the original. To generate down-sampled allelic coverage, we randomly selected reads at each variant site based on the down-sampling factor. The down-sampling was only applied to allelic coverage and therefore only affected the calculation of allelic fractions.

Results from this analysis demonstrate that single-copy DNA deletions with ≥50% clonality can be reliably detected at 10× or even 5× mean sequencing depth. An example of this analysis is presented in Fig. S7 and additional examples can be found in Pages 1-4 of Additional Dataset S1: DownSamplePlots.pdf. Below we will verify this result based on quantitative estimates of the allelic depth change and its statistical variation.

The statistical error of allelic depth can be estimated as

$$\delta\left(R(x)F(x)\right) \approx \overline{R(x)}\cdot\delta F(x) + \delta R(x)\cdot\overline{F(x)},$$

where $\overline{R(x)}$ and $\overline{F(x)}$ are the expected local sequence coverage and allelic fraction, and $\delta R(x)$ and $\delta F(x)$ are the statistical errors. At 10× mean sequencing depth, the statistical error of the phased average allelic fraction (restricted to intervals with 5 or more variants with coverage) is approximately

$$\delta F(x) \approx \sqrt{\frac{f(1-f)}{n}\left\langle\frac{1}{D}\right\rangle} \approx \sqrt{\frac{0.5\cdot 0.5}{5}\cdot\frac{1}{10}} \approx 0.07.$$

The statistical error of the total sequence coverage is approximately

$$\frac{\delta R(x)}{\overline{R(x)}} \approx \frac{1}{\sqrt{300}} \approx 0.06$$

based on 300 unique fragments in 10kb intervals. Therefore, the combined statistical error of allelic depth is approximately

$$\overline{R(x)}\cdot 0.07 + \overline{R(x)}\cdot 0.06\cdot\overline{F(x)}.$$

In disomic regions, $\overline{R(x)} \approx 2$ and $\overline{F(x)} \approx 1/2$, and the statistical error is

$$\delta\left(R(x)F(x)\right) \approx 0.1\cdot\overline{R(x)} \approx 0.2.$$

Based on Eq. (1) in **Online Methods**, the allelic depths are given by

$$A(x) = \frac{\alpha c_{\mathrm{A}}(x) + (1-\alpha)}{\alpha\tau/2 + (1-\alpha)}, \quad B(x) = \frac{\alpha c_{\mathrm{B}}(x) + (1-\alpha)}{\alpha\tau/2 + (1-\alpha)}$$

where $\alpha$ is tumor purity, $\tau$ is tumor ploidy, $c_{\mathrm{A,B}}$ are the integer allelic copy-number states. For near diploid genomes, $\tau \approx 2$, a single copy change $\Delta c_{\mathrm{A,B}} = \pm 1$ will result in a change in the normalized allelic depth of

$$\delta = \frac{\alpha}{\alpha\tau/2 + (1-\alpha)} \approx \alpha.$$

For near tetraploid genomes, $\tau \approx 4$, a single copy change will result in a change in the normalized allelic depth of

$$\delta = \frac{\alpha}{2\alpha + (1-\alpha)} \approx \frac{\alpha}{1+\alpha}.$$

Therefore, if we require the allelic depth change $\delta$ to be at least twice the standard error, then the minimum clonality of detectable single-copy changes in a near diploid genome at 10× depth is $\alpha = 0.2 \times 2 = 0.4$ and the minimum clonality of detectable single-copy changes in a near tetraploid genome is $\alpha = 2/3 \approx 0.67$.

We note that these thresholds are derived based on the allelic depths in 25kb intervals. In our copy-number workflow, we considered multiple consecutive 25kb intervals to identify regions of allelic imbalance and further performed joint haplotype inference based on allelic imbalance in all related BE/EAC samples from the same patient. The joint haplotype inference from allelic imbalance in multiple samples enables long-range haplotype phasing. With long-range haplotype information, we can perform phased averaging of allelic fractions in larger intervals to improve the accuracy of allelic depth calculation. In the final haplotype copy-number calculation, we chose 100kb intervals that have on average ~ 30 phased variants with coverage. This reduced the statistical error of allelic fraction to

$$\delta F(x) \approx \sqrt{\frac{f(1-f)}{n}\left\langle\frac{1}{D}\right\rangle} \approx \sqrt{\frac{0.5\cdot 0.5}{30}\cdot\frac{1}{5}} \approx 0.04$$

even at 5× mean sequencing depth. The statistical error of allelic depth is approximately

$$\delta\left(R(x)F(x)\right) \approx \overline{R(x)}\cdot(0.04+0.03) \approx 0.14$$

which is much lower than the change in allelic depth due to single-copy changes with clonality $\alpha = 40\%$ in either near diploid ($\delta \approx \alpha = 0.4$) or near tetraploid samples ($\delta \approx \alpha/(1+\alpha) = 0.29$).

In our cohort, samples from Patient 8 had the lowest sequencing depth (5 − 8×). Moreover, the three samples with the lowest depths: HGD1 (5.5×), HGD2 (6.3×), and IMEAC (7.2×) had high ploidy (HGD1:3.83; HGD2:2.83; IMEAC:2.9). The purity estimates for these clones were around 0.4. For the HGD1 sample with ploidy 3.83 and purity 0.39, the allelic depth difference due to single-copy difference is approximately

$$\delta = \frac{0.39}{3.83/2\cdot 0.39 + (1-0.39)} \approx 0.28.$$

The above estimation establishes that this change is significantly higher than the statistical error of allelic depth calculated for 100kb intervals (~ 0.14).

**Dependence on SCNA clonality.** We next analyzed the dependence of allelic depth difference on the clonality of copy-number alterations. In this analysis, we calculated the allelic depths in simulated tumor/normal mixtures with 20× mean sequence coverage and projected tumor cell fractions ("tumor purity") in the range of 10-40%. We first calculated the fractional depths of tumor reads and normal reads based on the estimated tumor cell fraction of the original sample, the projected tumor cell fractions, and the final sequencing depth of 20×. We then calculated the total sequence coverage $R(x)$ as a linear mixture:

$$R(x) = \alpha R_{\mathrm{tumor}}(x) + (1-\alpha)R_{\mathrm{normal}}(x).$$

For the allelic coverage, we performed separate down-sampling on the tumor allelic coverage and the normal allelic coverage based on the fractional sequencing depths and combined the allelic coverage to represent the final allelic coverage.

Results from this analysis suggest that the minimum clonality of single-copy changes that is resolvable from 25kb-interval allelic coverage is 18-20%. An example of this analysis is presented in Fig. S8 and additional examples can be found in Pages 5-8 of Additional Dataset S1: DownSamplePlots.pdf.

We note that most samples in our collection have purity above 30%, which is well above the detection limit (18%). The following samples have less than 30% purity: EAC from Patient 6 (27%), BE from Patient 9 (27%), HGD from Patient 13 (27%), BE (24%) and IND (16%) from Patient 14, and BE (28%), HGD1 (27%), HGD2 (27%), and IMEAC2 (24%) from Patient 15.

Below we discuss each sample separately.

EAC from Patient 6: This sample had estimated tumor purity of 27% and ploidy of 3.38, and was sequenced to 25× mean coverage. The average number of phased variants in 25kb intervals is about 10. Based on these numbers, we estimate the statistical error of allelic fractions in disomic regions to be

$$\delta F(x) \approx \sqrt{\frac{f(1-f)}{n}\left\langle\frac{1}{D}\right\rangle} \approx \sqrt{\frac{0.5\cdot 0.5}{10}\cdot\frac{1}{25}} \approx 0.03.$$

The statistical error of allelic depths is $\sim 0.03\times 2 = 0.06$, which is significantly lower than the allelic depth difference expected for single-copy changes

$$\delta = \frac{0.27}{0.27\cdot 3.38/2 + 1 - 0.27} = 0.26.$$

BE from Patient 9: The only SCNA in this sample was 9p uniparental disomy (UPD). UPDs cause reciprocal single-copy gain and loss to both homologs, resulting in an allelic depth difference of

$$A(x) - B(x) = 2\times\frac{\alpha}{\alpha\tau/2 + (1-\alpha)}.$$

In this sample, the estimated ploidy is 1.96 and the estimated purity is 27%. Therefore the allelic depth difference on 9p is

$$A(x) - B(x) = \frac{2\cdot 0.27}{0.27\cdot 1.96/2 + 1 - 0.27} \approx 0.54,$$

which is significantly higher than the statistical error ($\sim 0.14$). Moreover, the resolution of Chr.9 haplotype in this patient based on allelic imbalance in HGD2, EAC1, and IMEAC2 samples ensured the accuracy of 9p allelic depth calculation in the BE sample (10× sequencing depth, near diploid).

HGD from Patient 13: This sample had estimated tumor purity of 27% and ploidy of 3.62, and was sequenced to 23× mean coverage. The statistical error of allelic depths and the allelic depth difference for single-copy changes are both similar to the EAC sample from Patient 6. The sequencing coverage is therefore sufficient for the detection of single-copy changes.

BE and IND from Patient 14: Both samples were near diploid and shared 9p UPD, whose detection sensitivity is higher because of the two-copy difference between the two homologs. The IND sample (21.7×) also contained a single-copy gain of Chr.8. The detection of Chr.8 gain in this sample was enabled by the determination of Chr.8 haplotype phase based on allelic imbalance in samples HGD1 and IMEAC from the same patient.

BE, HGD1, HGD2, and IMEAC2 from Patient 15: These samples were sequenced to at least 23.2× (HGD2). The statistical error of allelic fractions is approximately $\delta F(x) \sim 0.03$. Because all the samples have near diploid genomes, the allelic depth difference due to single-copy changes is approximately equal to the purity (0.24-0.28). Therefore, the sequencing depth is sufficient to detect clonal single-copy changes (e.g., Chr.12q and Chr.17p). The biggest challenge with samples from this patient is the presence of many subclonal copy-number changes. This is overcome by joint haplotype inference based on allelic imbalance in all samples. These include completely resolved haplotypes on Chr.1 and Chr.2 (HGD1,HGD/IM), Chr.4 (HGD1, HGD/IM, IMEAC2), Chr.5 (HGD/IM, BE, IMEAC2, HGD2, EAC1), Chr.6 (BE, IMEAC2, HGD2, EAC1), Chr.7 and Chr.8 (IMEAC2, EAC1), Chr.9 (HGD1,HGD/IM, EAC1), Chr.10 (IMEAC2, EAC1), Chr.11p (BE, IMEAC2, HGD2), Chr.13q (HGD1, HGD/IM), Chr.15q (HGD2, EAC1), Chr.18 (HGD/IM, IMEAC2), Chr.19 (HGD/IM, BE, IMEAC2, HGD2), Chr.20 (HGD1, HGD/IM, IMEAC2), Chr.21q (HGD/IM, BE, IMEAC2, HGD2), Chr.22q (BE, IMEAC2, HGD2). The phylogeny of these samples was determined based on segmental copy-number alterations on Chr.2q and Chr.11p (see Fig. S6 and caption for more details).

In summary, the tumor/normal mixture benchmarking and the statistical estimation demonstrate the capability to reliably detect single-copy changes with 20% or higher clonality at 20× sequencing depth through a combination of phased average allelic fraction and joint haplotype inference from multiple samples with allelic imbalance. *We note that in a single tumor sample, the minimum clonality of detectable SCNAs will depend on the uniformity of sequence coverage in the DNA library and may not reach 20% for FFPE libraries with significant variation in the sequence coverage.*

**Validation of allelic-imbalance based haplotype inference.** In the last subsection of this document, we provide validation of the method of haplotype inference based on allelic imbalance. For this validation, we used the whole-genome sequencing data of clones expanded from single post-crisis retina pigmental epithelium (RPE-1) cells generated in Ref. 37 together with truth haplotype data of RPE-1 cells generated in a previous study from us (Tourdot et al., Genome Biology 2021). The truth haplotype was determined by computational haplotype inference from linked-reads and Hi-C sequencing of RPE-1 cells and validated by the sequencing of monosomic RPE-1 cells.

We had previously used the RPE-1 haplotype data to calculate haplotype-specific DNA copy number of both parental chromosomes in these clones and resolved all segmental copy-number alterations. For this validation, we combined statistical haplotype phasing and allelic-imbalance based haplotype inference to calculate haplotype-specific DNA copy number. These results were then compared against results derived using the truth haplotype for validation.

The alignment and post-alignment processing of the sequencing reads were described in our previous paper (Tourdot et al., 2021). Reads were aligned to the GRCh38 reference. We performed genotyping and statistical phasing using the newly generated reference haplotype data from the 1000 Genomes project (available from http://ftp.1000genomes.ebi.ac.uk/vol1/ftp/data_collections/1000G_2504_high_coverage/working/20201028_3202_phased/). These data were generated by variant calling and phasing (with `shapeit2`) of sequencing reads that were aligned to GRCh38, which may have some differences from the liftover of reference haplotypes from GRCh37 to GRCh38. Haplotype phasing and haplotype-specific copy-number calculation were performed using the same algorithm as described in ***Haplotype refinement and phasing of somatic copy-number alterations using allelic imbalance*** in **Online Method**. Results from this analysis are presented in Additional Dataset S2: AllelicDepthPhasing.pdf and selected examples are presented in Figs. S9-S11.

In comparison to the WGS data of BE/EAC samples, the RPE-1 WGS data had better quality as they were derived from live cells instead of degraded FFPE DNA. The RPE-1 libraries were also sequenced deeper than the BE/EAC samples (30-40× mean depth). Because of these advantages, the RPE-1 data had both better variant coverage and better uniformity of total sequence coverage. To make a fair comparison, we chose to perform haplotype inference using allelic depths calculated for 10kb intervals, which have similar allelic coverage as 25kb allelic depths in the BE/EAC data. Importantly, *many copy-number alterations in the RPE-1 data were inferred to be subclonal*. This enabled us to evaluate the capability to detect subclonal SCNAs and perform haplotype inference based on subclonal allelic imbalance.

In Additional Dataset S2, we first present genome-wide copy-number plots of all RPE-1 clones (Pages 1-2), with red and blue dots correspond to the 1Mb-level DNA copy number of each parental haplotype. On Page 3, we show the results of haplotype phasing from copy-number gains of 10q and 12p that are shared by all RPE-1 clones. (The 10q example is included in Fig. S9.) These examples demonstrate that the phasing algorithm can accurately resolve the long-range haplotype phase in large regions of allelic imbalance that is indicated by the agreement between the allelic-imbalanced derived haplotype and the truth haplotype. In pages 4-20 of Additional Dataset S2, we show the results of haplotype phasing in all chromosomes with allelic imbalance (determined using the truth haplotype) in all post-crisis RPE-1 clones. For each clone, the first plot shows the genome-wide DNA copy number and each subsequent panel shows the results of haplotype phasing on a single chromosome with either arm-level or segmental allelic imbalance.

We manually reviewed all chromosomes with partial or complete allelic imbalance identified with the complete haplotype data (40 chromosomes total). The results are summarized below.

- X-29 (one instance): Chr.11 (no error)
- X-32 (one instance): Chr.18 (one switching error at the centromere)
- 24-141 (two instances): Chr.7 (no error), Chr.12 (no error, opposite haplotype assignment of iso-12p is unrelated to alterations to 12q; **sloping copy-number variation at the 12q-terminus**)
- 24-144 (two instances): Chr.3 (no error; minor subclonal SCNAs on 3p affect both homologs and are unresolvable), Chr.6 (unable to resolve subclonal gains on 6p with allelic depth difference ≈ 0.1, 6p-terminus was phased correctly).
- X-33 (four instances): Chr.1 (unable to resolve subclonal 1q loss with allelic depth difference ≈ 0.09), Chr.8 (one switching error on 8p, no error in complex segmental gains on 8q), Chr.18 (no error; **note the presence of subclonal 18q-terminal loss with very low clonal fractions**), Chr.X (no error).

- I-dox-1 (five instances): Chr.2 (two switching errors on 2q near the centromere), Chr.7 (one switching error on 7q), Chr.8 (one switching error on 8p), Chr.11 (no error), Chr.15 (one switching error on 15q).
- X-25 (five instances): Chr.4 (no error; **SCNAs on both homologs**), Chr.8 (no error), Chr.12 (no error), Chr.13 (no error; **partial retention near the 13q terminus with very low clonal fractions**), Chr.X (no error)
- X-37 (six instances): Chr.1 (one switching error of the 1q-terminal deletion), Chr.14 (no error), Chr.15 (no error; **sloping copy-number variation**), Chr.17 (one switching error on 17p), Chr.18 (no error; **sloping copy-number variation on 18q**), Chr.22 (unable to resolve subclonal 22q loss with allelic depth difference 0.09 (q-ter) – 0.13 (pericentric)).
- X-36 (six instances): Chr.2 (one switching error on 2p **with sloping copy-number variation**; 2q haplotype derived from subclonal loss with allelic depth difference ∼ 0.16 had a few local errors), Chr.4 (one switching error at the centromere), Chr.6 (a few local errors at the 6q terminus with allelic depth difference ∼ 0.20; **this region has sloping copy-number variation**), Chr.7 (no error), Chr.10 (no error), Chr.18 (one switching error at the centromere)
- X-35 (eight instances): Chr.1 (no error), Chr.5 (a few local errors on 5q:50-100Mb with allelic depth difference ≈ 0.16; **sloping copy-number variation**), Chr.6 (no error; **sloping copy-number variation near 6q-terminus**), Chr.8p (unable to resolve subclonal gain of 8p with allelic depth difference ≈ 0.08), Chr.10 (two switching errors on 10q near the centromere), Chr.12p (no error), Chr.16 (two switching errors on 16p), Chr.18 (one switching error at the centromere; **sloping copy-number variation on 18p and 18q**).

Except for four SCNAs with clonality <14% (24-144:Chr.6; X-33:Chr.1; X-37:Chr.22; X-35:Chr.8) that can only be resolved with the truth haplotype data, all the other SCNAs can be resolved using 10kb allelic depth data. For most chromosomes, the haplotype inferred by allelic imbalance has either no (20/40) or one long-range switching error (10/40) that is often at the centromere or near regions with low complexity or variant density. Three chromosomes had two switching errors. These switching errors are easily identifiable. Three copy-number alterations with clonality of 15-20% are resolvable but contain some local phasing errors (X-36:Chr.2q; X-36:Chr.6; X-35:Chr.5). In particular, our phasing algorithm can generate correct haplotype inference even in regions where both homologs are altered and in regions with complex segmental gains and losses (Fig. S10). Together, these results demonstrate the robustness of our algorithm for allelic-imbalance based haplotype inference.

We note that this analysis also provides *independent evidence about the copy-number outcomes of breakage-fusion-bridge cycles or telomere crisis* in addition to the results from our prior work (Ref. 38) as the copy-number alterations in post-crisis RPE-1 clones are initiated by telomere de-protection. In particular, we identified multiple examples of sloping copy number variation, two of which are shown in Fig. S11. These results demonstrate the robustness of our haplotype-inference algorithm even when one homolog has non-constant copy number.

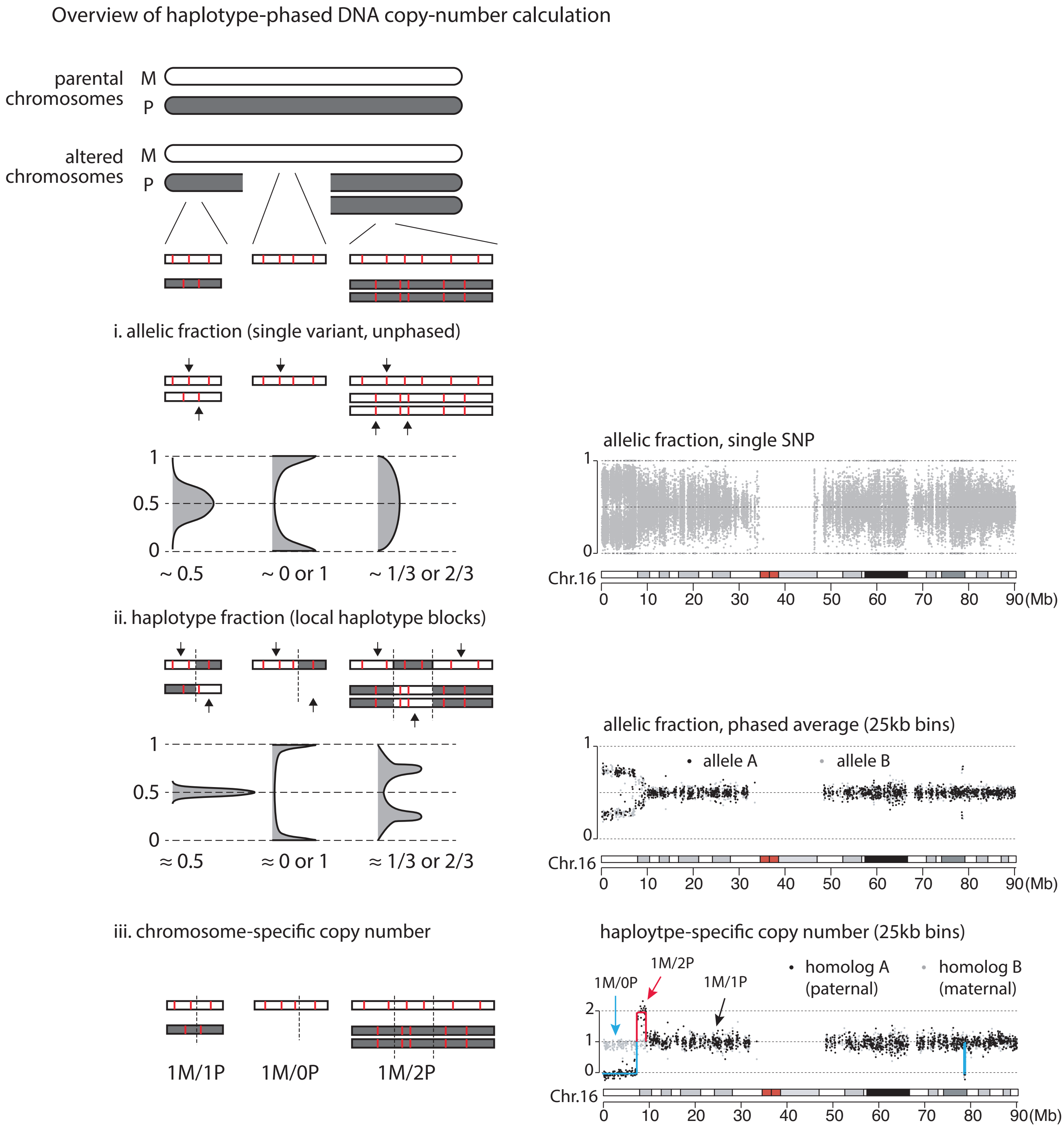


**Fig. S1.** General idea of haplotype-specific DNA copy-number calculation based on (i) allelic fractions at single polymorphic sites; (ii) phased average allelic fractions in local haplotype blocks; and (iii) chromosome-specific allelic fractions. (Left) Schematic illustration of expected results; (right) real data from Chr.16 in the HGD2 sample from Patient 4.

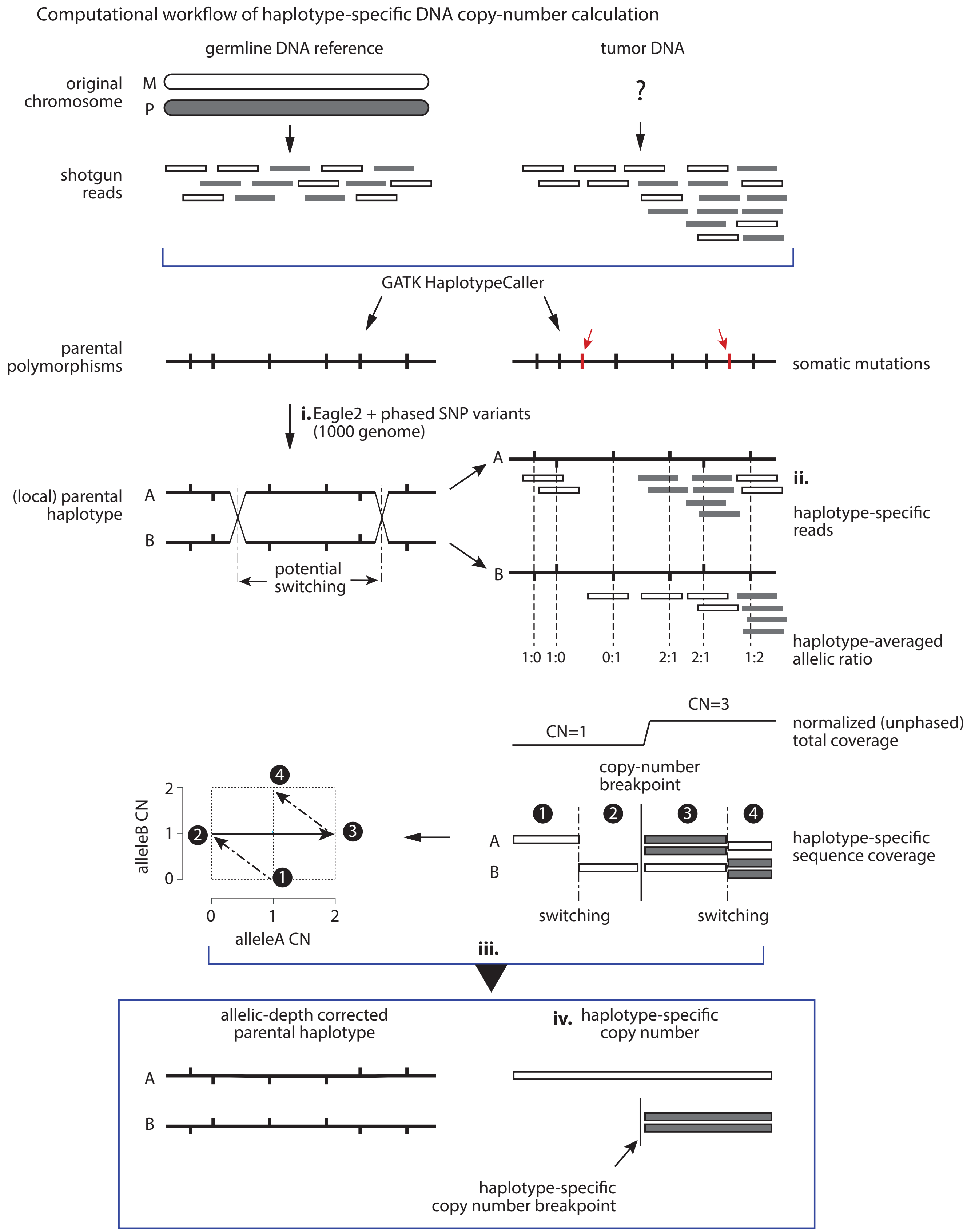


**Fig. S2.** Overview of the computational workflow of haplotype-specific copy-number calculation: (i) statistical haplotype phasing (middle, left); (ii) calculation of local allelic fractions and allelic coverage (middle, right); (iii) correction of haplotype switching errors by allelic imbalance (lower middle); (iv) final haplotype copy-number calculation (bottom). See **Online Methods** for a detailed description of each step.

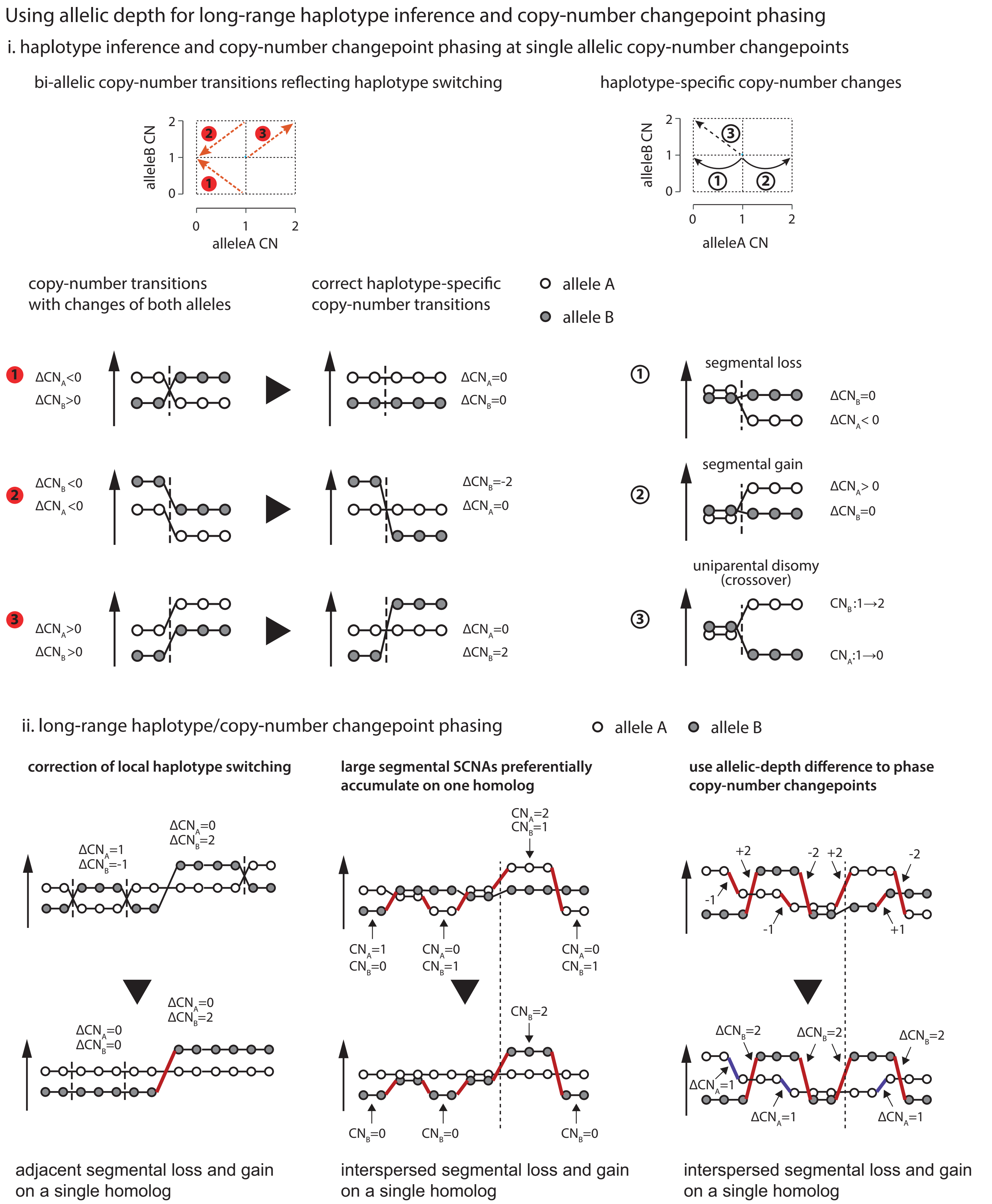


**Fig. S3.** Strategies of long-range haplotype inference by allelic imbalance: i. allelic copy-number transitions due to haplotype switching errors (left, changes in both allelic depths) and reflecting true somatic copy-number alterations (right, changes in only one allelic depth, except uniparental disomy); ii. long-range phasing of copy-number alterations/changepoints based on continuous allelic imbalance (left), allelic imbalance with interspersed allelic balance (middle), and allelic imbalance with different allelic ratios (right).

Examples of long-range haplotype inference and copy-number changepoint phasing in single samples

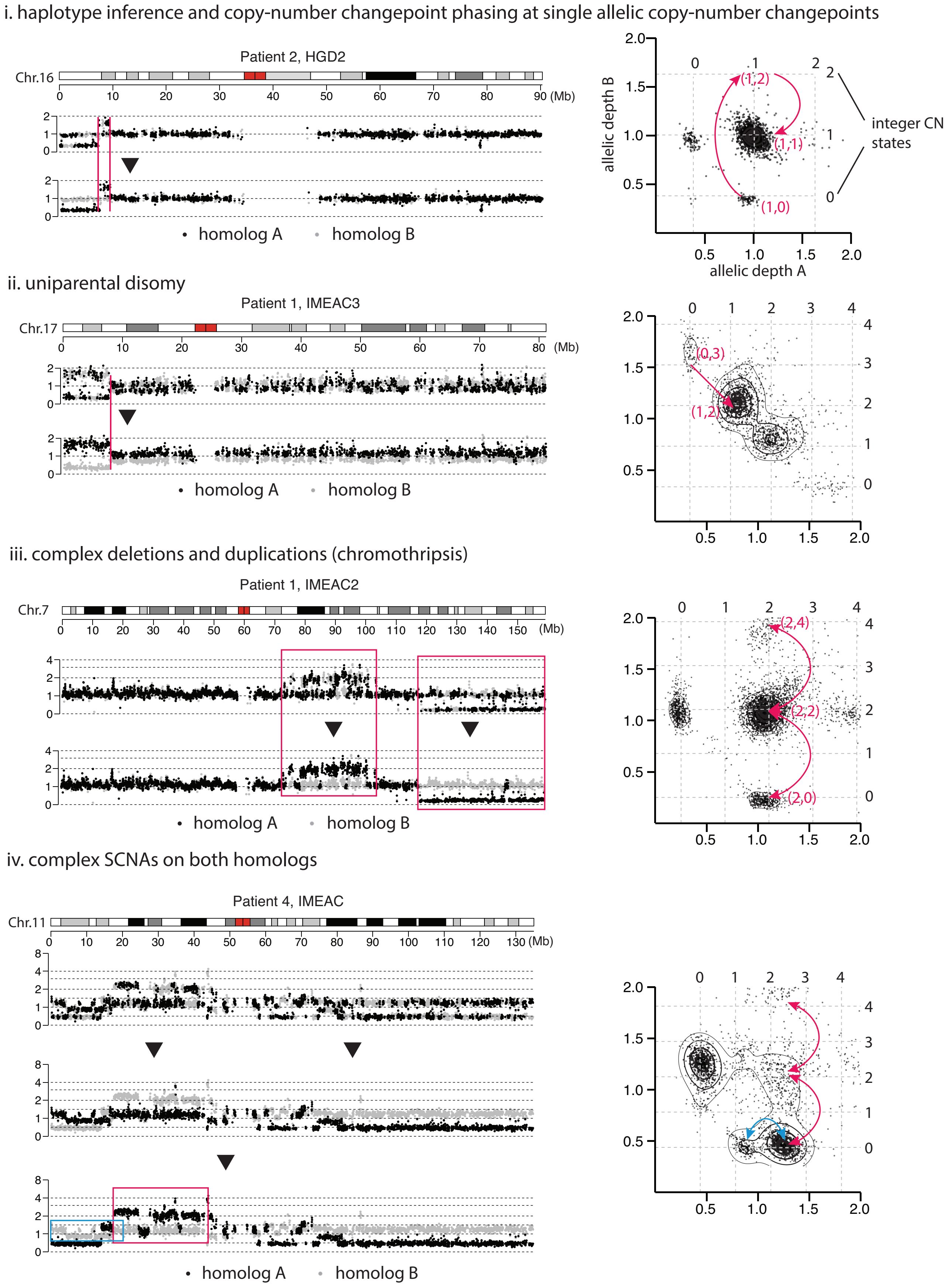

**Fig. S4.** Examples of haplotype phasing by allelic imbalance in a single sample. (i) contiguous allelic imbalance with adjacent deletion and duplication on one homolog; (ii) uniparental disomy; (iii) two regions of allelic imbalance separated by a region of allelic balance; (iv) complex copy-number alterations on both homologs with different allelic depths changes. (Left) Normalized allelic depths (gray and black dots for opposite homologs) calculated from the statistical haplotype (top) and after correction of haplotype switching (bottom); (right) scatter plots of allelic copy-number states and transitions (arrows); the integer copy-number states determined after purity/ploidy correction are annotated by gray dashed lines. The different allelic copy-number states are annotated in red, with transitions (red arrows) annotated on the left. See Fig. S3 for schematic illustrations of allelic depths and transitions. In (iii), we expect the terminal deletion and the internal duplication to occur on a single homolog in two successive BFB cycles, the first generating the terminal deletion ("chromatid type") with a small region of retention, the second generating the internal duplication ("chromosome type") with interspersed DNA losses. Regions with altered DNA copy number generated by these two events are outlined on the left. See Figure 5 for further information about the different outcomes of breakage-fusion-bridge cycles and Fig. S3ii, middle panel for the incorporation of this mechanistic insight in the computational haplotype inference. In (iv), the final haplotype phase of the p-arm (from the middle to the bottom panel) is inferred based on the different changes in allelic depths: $\Delta$CN $\approx 1.2$ for the black homolog reflecting three-copy changes (red arrows on the right) and $\Delta$CN $\approx 0.4$ for the gray homolog reflecting single-copy changes (cyan arrow). Regions with oscillating copy number on each homolog are outlined on the left with matching colors. As this sample (Patient 4, IMEAC) is inferred to have undergone whole-genome duplication (WGD), alterations to the black homolog with three-copy changes most likely occurred before WGD, whereas alterations to the gray homolog with single-copy changes occurred after WGD. This example underscores the importance of haplotype-specific copy-number analysis for both evolutionary and phylogenetic inferences.

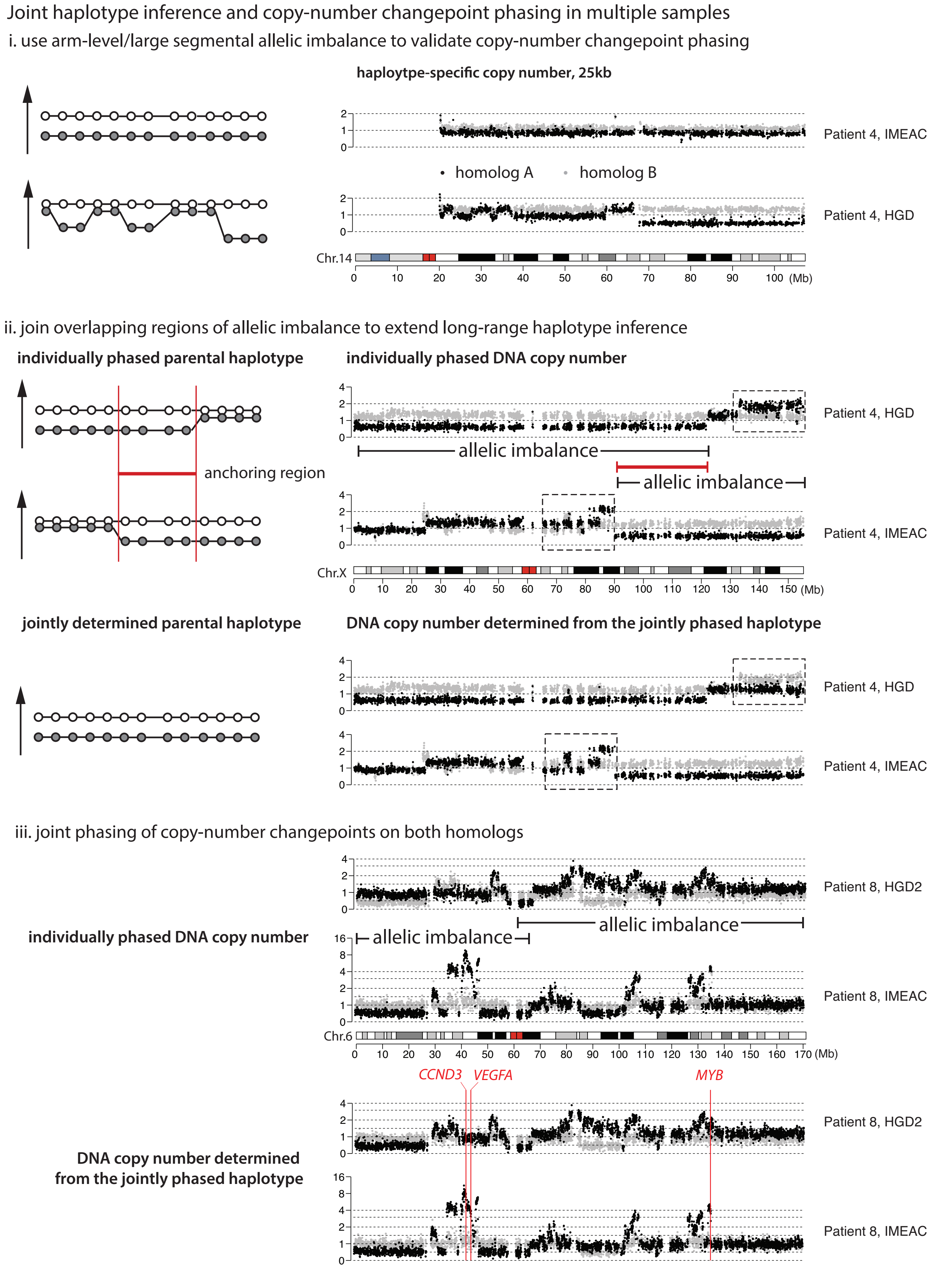
Joint haplotype inference and copy-number changepoint phasing in multiple samples
i. use arm-level/large segmental allelic imbalance to validate copy-number changepoint phasing
haploytpe-specific copy number, 25kb
Patient 4, IMEAC
homolog A
homolog B
Patient 4, HGD
Chr.14
(Mb)
ii. join overlapping regions of allelic imbalance to extend long-range haplotype inference
individually phased parental haplotype
individually phased DNA copy number
Patient 4, HGD
allelic imbalance
anchoring region
allelic imbalance
Patient 4, IMEAC
Chr.X
jointly determined parental haplotype
DNA copy number determined from the jointly phased haplotype
Patient 4, HGD
Patient 4, IMEAC
iii. joint phasing of copy-number changepoints on both homologs
Patient 8, HGD2
allelic imbalance
individually phased DNA copy number
allelic imbalance
Patient 8, IMEAC
Chr.6
CCND3
VEGFA
MYB
Patient 8, HGD2
DNA copy number determined
from the jointly phased haplotype
Patient 8, IMEAC

**Fig. S5.** Examples of haplotype correction from allelic imbalance in multiple samples from one individual. (i) Allelic imbalance in the IMEAC sample is used to derive the complete chromosomal haplotype that is then used to calculate/validate SCNAs in the HGD sample. (ii) Allelic imbalance in the HGD sample (0-120Mb) and in the IMEAC sample (90-qter) are joined based on the haplotype phase in the overlapping region of allelic imbalance in both samples (90-120Mb). The jointly-inferred haplotype is then used to calculate chromosome-specific DNA copy number (shown on the bottom). (iii) Another example of joint haplotype inference based on overlapping allelic imbalance that resolves complex SCNAs on the black homolog and a large segmental deletion (85-100Mb) of the gray homolog in HGD. There are several noteworthy observations in this example. First, the HGD1/HGD2/IMEAC samples have the lowest sequencing depths among all samples (HGD1:$5.5\times$; HGD2:$6.3\times$; IMEAC:$7.2\times$); the agreement between the 6p haplotypes that are independently derived from each of the three samples demonstrates the accuracy of our computational algorithm for allelic-imbalanced based haplotype correction even when it is applied to individual samples sequenced at $5-7\times$ depth. Second, the distinct patterns of deletions and duplications/amplifications in the HGD2 and IMEAC genomes are inferred to have arisen from a single unstable ancestral chromosome based on multiple shared copy-number breakpoints including the shared boundary of the p-terminal deletion. Moreover, we inferred an ancestral WGD event preceding the divergence of these two samples. Based on the complete deletion at the 6p terminus, the ancestral unstable Chr.6 could have been generated prior to WGD but undergone multiple rounds of alterations both before and after WGD that create focal amplifications. Third, HGD1 and IMEAC have nearly identical copy-number breakpoints but display different copy-number states (see copy-number plots in **Downloadable Supplementary Data**); this pattern is consistent with complex ecDNAs arising from chromothripsis (which generates interspersed DNA deletions between the focally amplified regions) but undergoing asymmetric segregation to create copy-number heterogeneity. This inference is consistent with results from fluorescence in-situ hybridization (FISH) analysis of *VEGFA* that is located at ≈43.7Mb. We further note that the amplified DNA in the HGD1/IMEAC genomes also contains two other oncogenes, *CCND3* at 41.9Mb and *MYB* at 135.2Mb (GRCh37 coordinates), but the amplified DNA in HGD2 contained none. It is plausible that the amplified DNA was generated by an ancestral chromothripsis event but distributed into two daughter cells in a reciprocal manner; the daughter cell that inherited oncogenic DNA fragments subsequently acquired oncogenic amplifications and became the ancestor to HGD2 and IMEAC, whereas the sibling cell was the ancestor of HGD1 that did not transform to EAC.

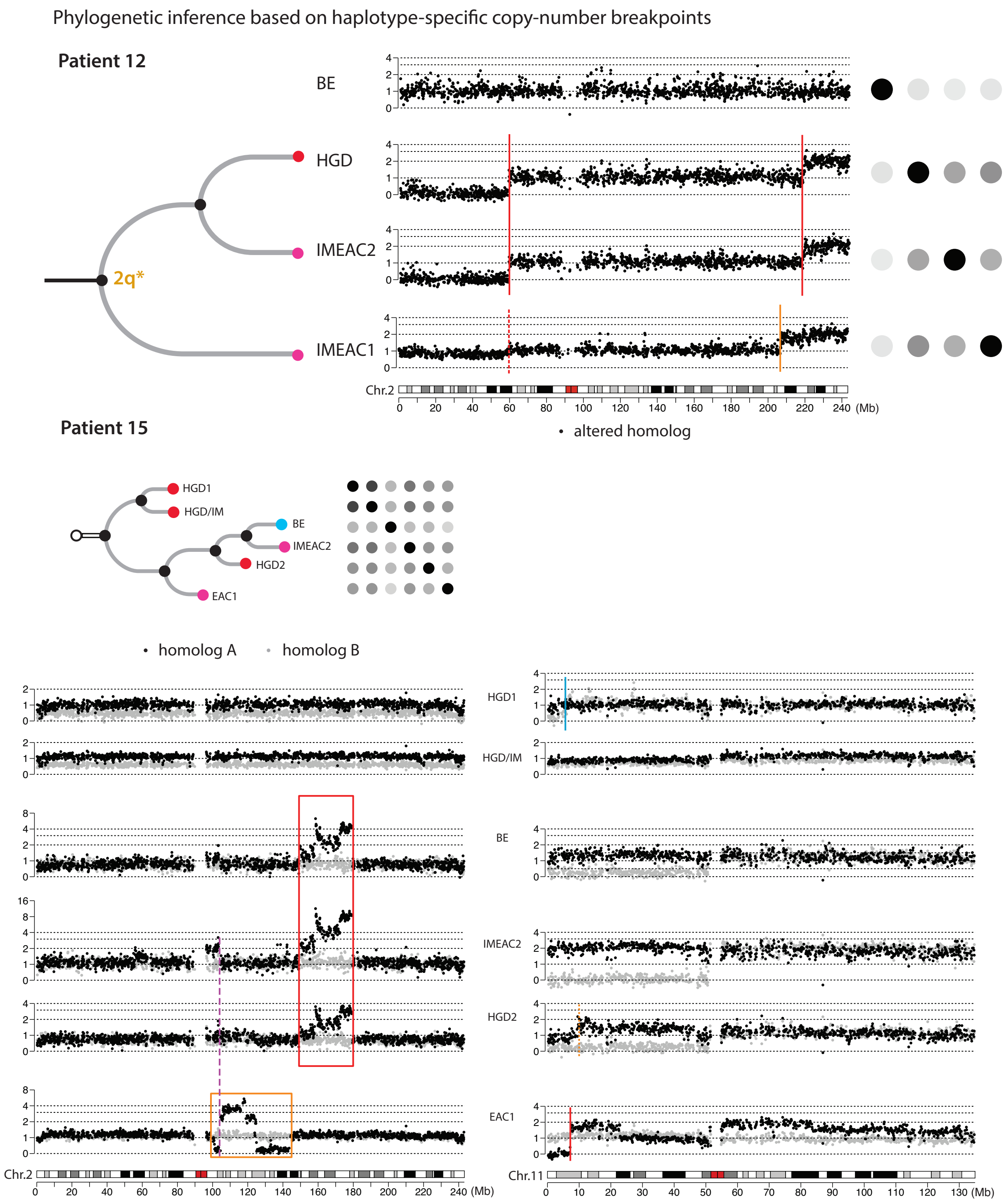


**Fig. S6.** Phylogenetic inference based on haplotype-specific copy-number changepoints. (Top left) Phylogenetic relationship between HGD, IMEAC1, and IMEAC2 samples from Patient 12 derived from copy-number changepoints. (Top middle) Haplotype-specific copy number changepoints on Chr.2 in each genome. Note that the breakpoint at 60Mb shared between HGD and IMEAC2 is also present in a subclonal population in IMEAC1 as indicated by the minor decrease in allelic depth; also note that the breakpoint near 220Mb that is shared between HGD and IMEAC2 and establishes their kindred relationship has a related breakpoint near 208Mb in IMEAC1. (Top right) Pairwise SNV similarity estimated from the fraction of shared sSNVs between these samples. (See Extended Data Figure 2 for the color bar.) Although the SNV similarity between HGD and IMEAC1 is slightly higher than between HGD and IMEAC2, this is most likely due to two technical reasons: (1) the lower sequencing depth of IMEAC2 (14.1×) in comparison to IMEAC1 (26.8×) causes a higher false-negative rate of variant detection in the IMEAC2 sample; (2) the presence of subclonal HGD/IMEAC2 cells in the IMEAC1 sample (as indicated by the subclonal copy-number breakpoint at 60Mb) confounds the calculation of SNV similarity. (Middle) SCNA-derived phylogenetic tree and pairwise sSNV similarity of samples from Patient 15. (Bottom) Evidential support for the SCNA-based phylogenetic inference. The complex SCNA patterns on Chr.2q (black homolog) and 11p loss (gray homolog) shared between BE, IMEAC2, and HGD2 establish their kindred relationship; the shared breakpoint (magenta dashed line) with complementary copy-number changes in IMEAC2 and EAC1 indicates a reciprocal distribution of broken fragments of an ancestral chromosome. The absence of any SCNA breakpoint on Chr.2 in HGD1 and HGD/IM suggests that their common ancestor was distinct from the common ancestor of the other four samples. Note the unrelated 11p-terminal deletion in HGD1 on the gray homolog and the terminal deletion and duplication in EAC1 on the black homolog that will confound phylogenetic inference from unphased SCNA breakpoints. The apparently high sSNV similarity between HGD1, HGD/IM and IMEAC2 is most likely the outcome of false negative variant detection in BE, HGD2, and IMEAC2 genomes due to the low clonal fractions of tumor cells (BE:28%; HGD2:27%; IMEAC2:24%).

**Patient 7, HGD2. Purity: 60%; ploidy: 1.91** | Allelic depths in 25kb intervals calculated from down-sampled allelic coverage on Chr.18 with mean sequencing depths of 30×, 20×, 10×, 5×.

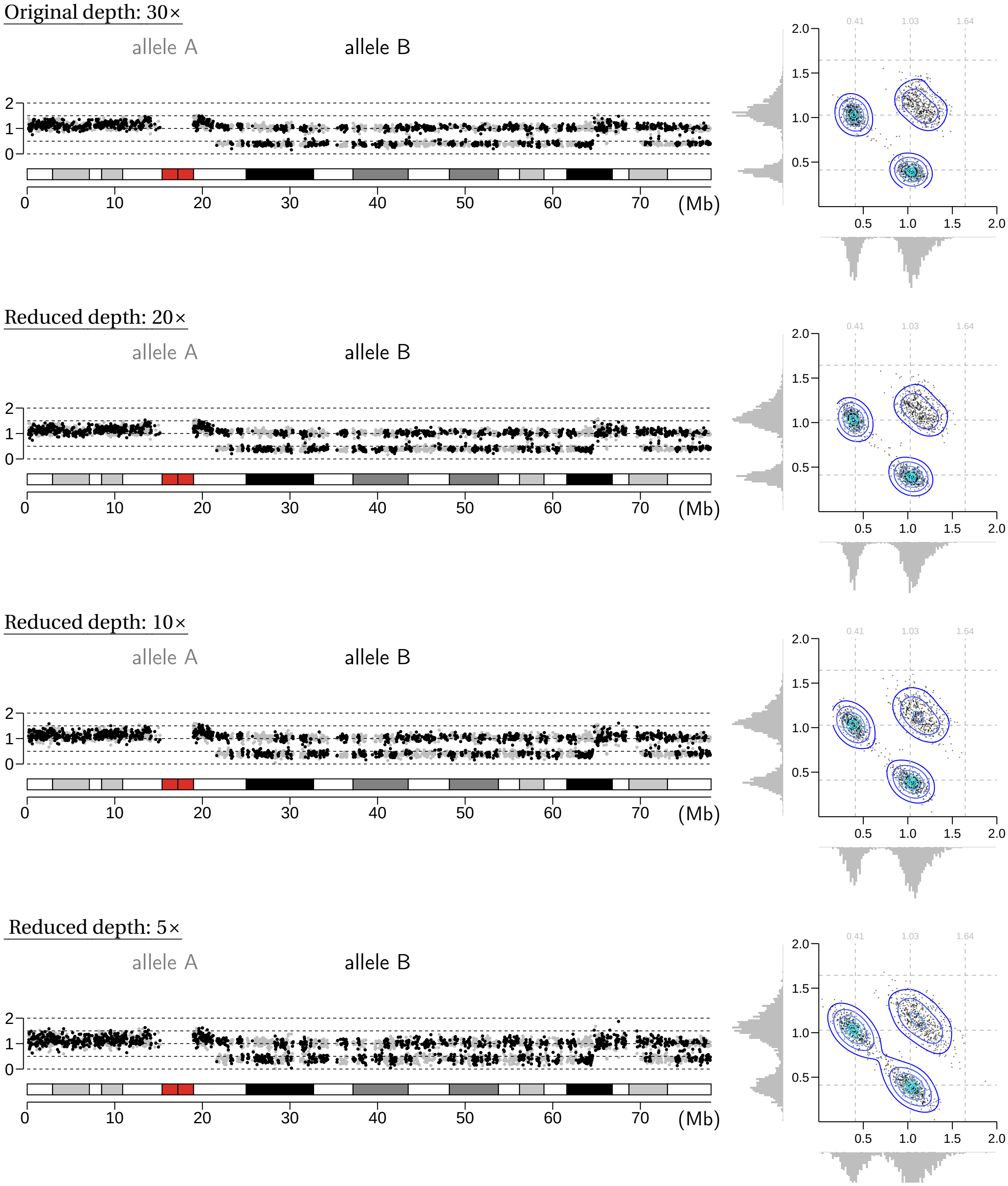


**Fig. S7.** Allelic depths in 25kb intervals calculated from the original (30×) and down-sampled allelic coverage of Chr.18 in sample HGD2 from Patient 7. Left panels show the allelic depths along Chr.18: Black and gray dots represent the normalized depth of each allele ("allele A" and "allele B"). Right panels show the scatter plots and distributions of allelic copy-number states: X- and Y-axis represent allele A and allele B depths; dotted gray lines represent allelic depths (annotated with numbers in gray) corresponding to integer copy-number states $0, 1, 2, \ldots$; the contour map represent the 2D distribution of allelic copy-number states, the histograms on each axis represent the 1D distribution of allelic copy-number states. The reduced allelic depth on 18q is inferred to be a clonal deletion (tumor purity 60%). The separation between different allelic copy-number states (in both 2D and 1D distributions) ensures the accuracy of long-range haplotype phasing based on allelic depth differences. There is a minor increase in the statistical variations of allelic depths when the sequencing depth is reduced from 20× to 10× and a bigger increase as when reduced to 5×. This analysis suggests that this deletion can be reliably identified and phased even at 5× mean sequencing depth.

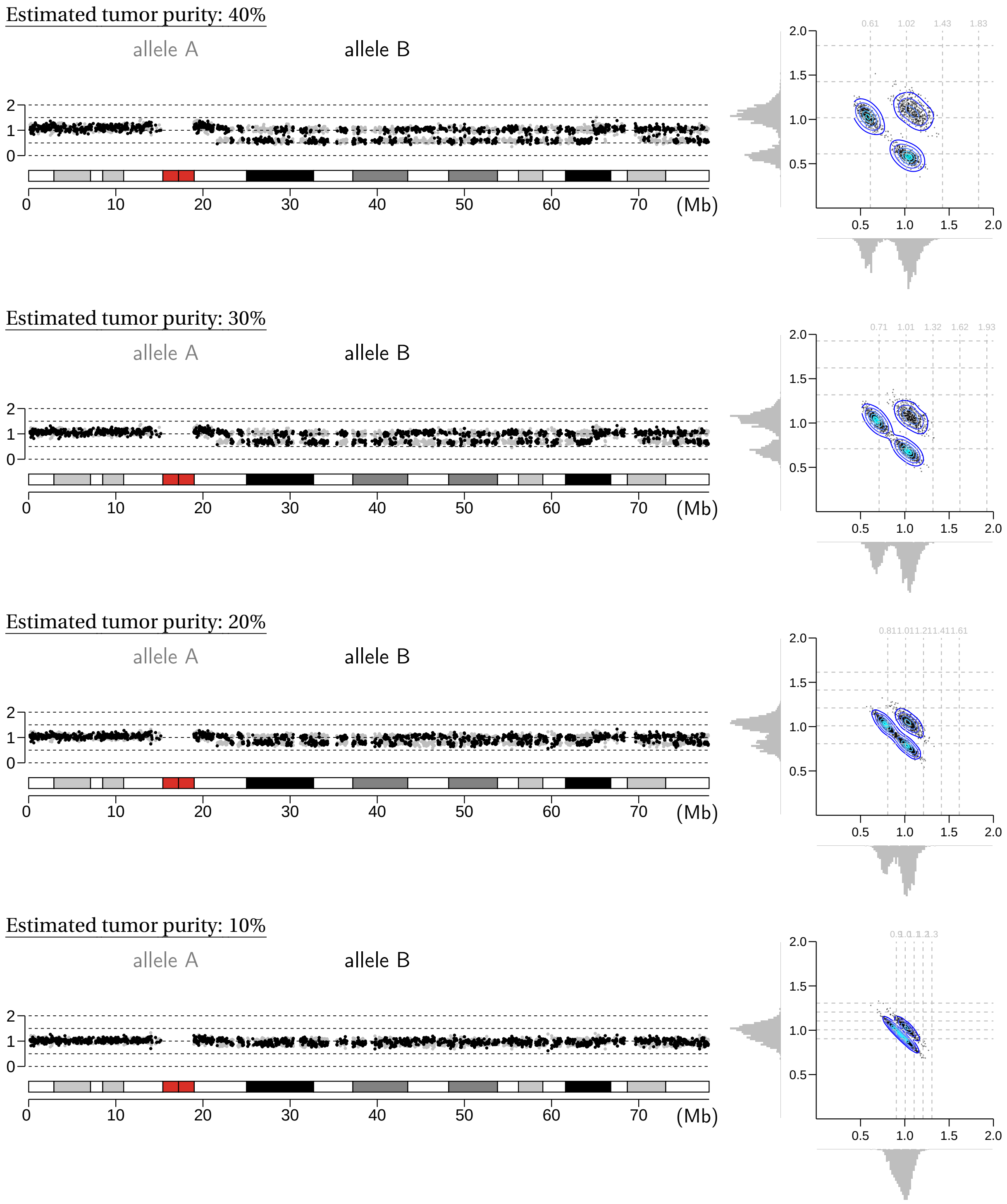
Estimated tumor purity: 40%
allele A
allele B
(Mb)
0.61 1.02 1.43 1.83
Estimated tumor purity: 30%
allele A
allele B
(Mb)
0.71 1.01 1.32 1.62 1.93
Estimated tumor purity: 20%
allele A
allele B
(Mb)
Estimated tumor purity: 10%
allele A
allele B
(Mb)

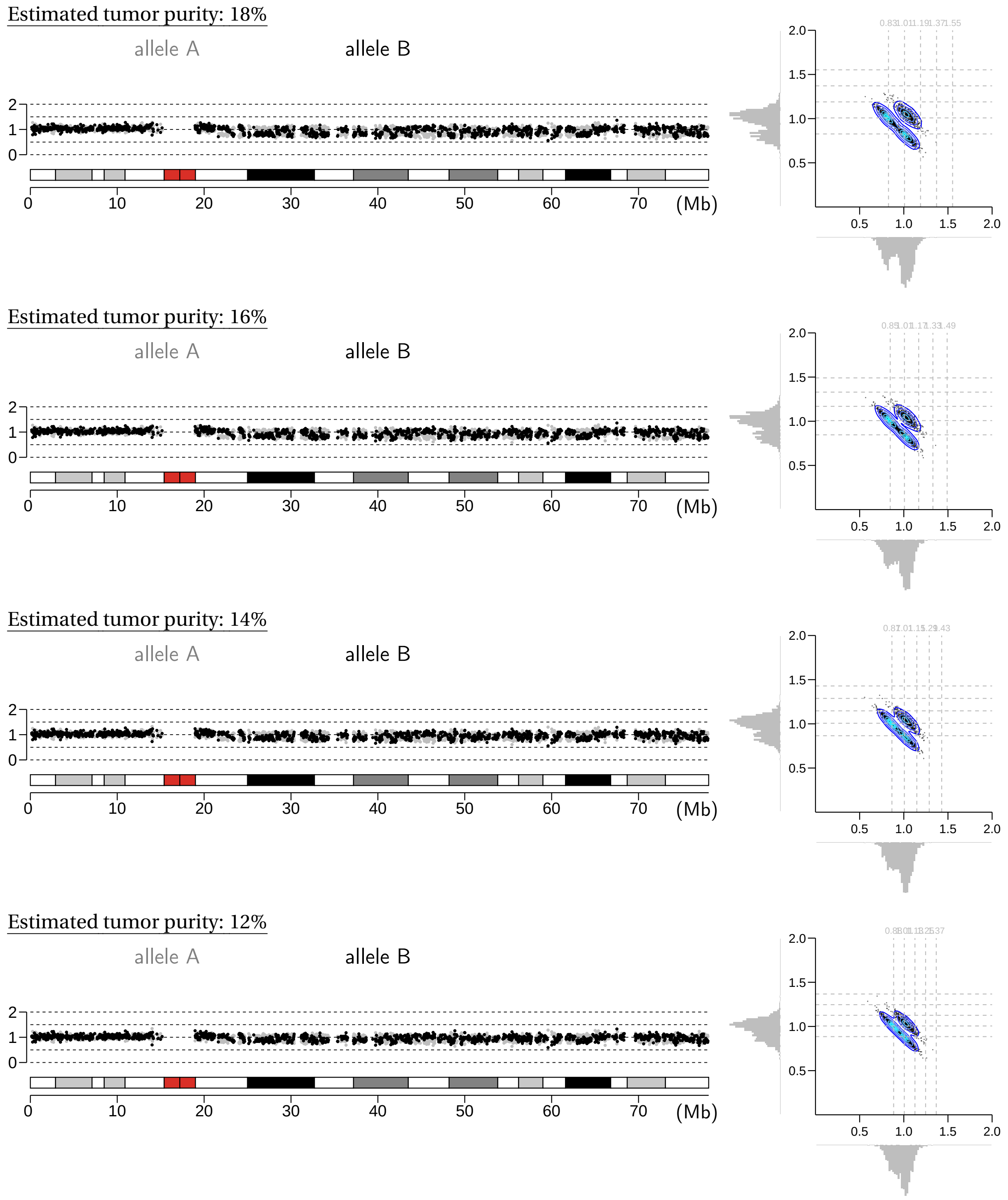


**Fig. S8.** Allelic depths calculated from the allelic coverage of *in silico* mixtures of tumor and matching germline samples. The combined sequencing depth is $20\times$ and the estimated tumor cell fractions in the mixtures are 40%, 30%, 20%, 10%, 18%,16%,14%,12%. Left and right panels are the same as in Fig. S7. Based on the allelic depth distributions, the lowest clonality of single-copy changes that can be resolved from allelic depth differences is 18-20%.

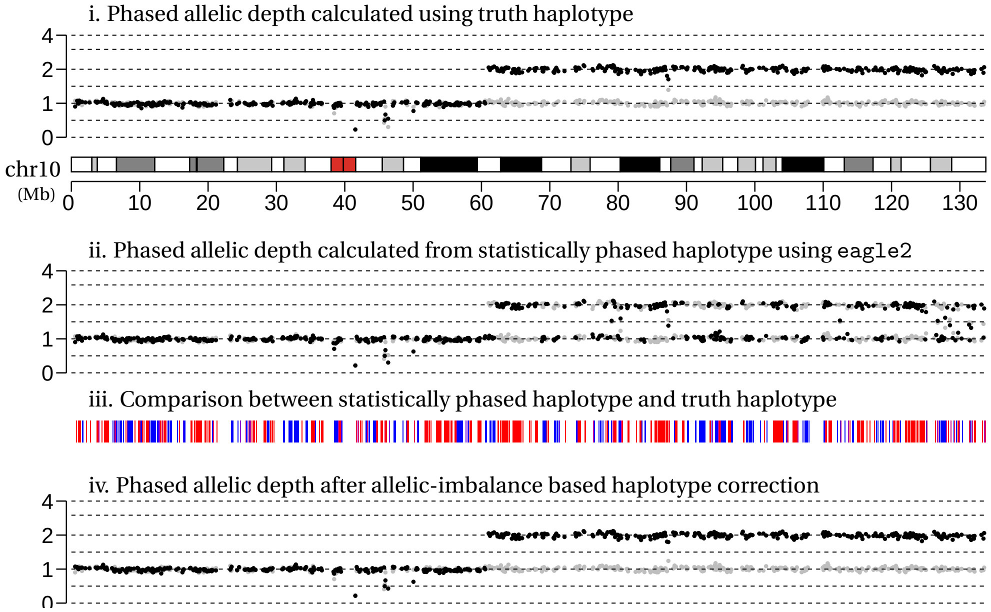


**Fig. S9.** An example of haplotype inference from a single region with contiguous allelic imbalance. Black and gray dots correspond to 100kb allelic depths calculated using the complete chromosomal haplotype phase determined from long-range sequencing (top, i), statistical haplotype (middle, ii), and allelic-imbalance corrected haplotype (bottom, iv). Note that black and gray dots are arbitrarily assigned to haplotype A and B in the truth data (i) and in the statistical phasing results (ii and iii) as they are calculated independently. When comparing the statistical haplotype or the allelic-imbalance corrected haplotype against the truth haplotype data, we use red and blue bars to denote agreement with complementary parental haplotypes: Oscillations between blue and red indicate haplotype switching; consecutive blue or red indicate consistency with a single parental haplotype. The example here showed that our algorithm correctly resolves the long-range haplotype in the region of allelic imbalance (from 61Mb to the q-terminus) by eliminating switching errors (oscillation between blue and red) in the statistical haplotype. Note that the long-range haplotype inference is restricted to the region with allelic imbalance and leaves out the region in allelic balance (p-terminus to 61Mb).

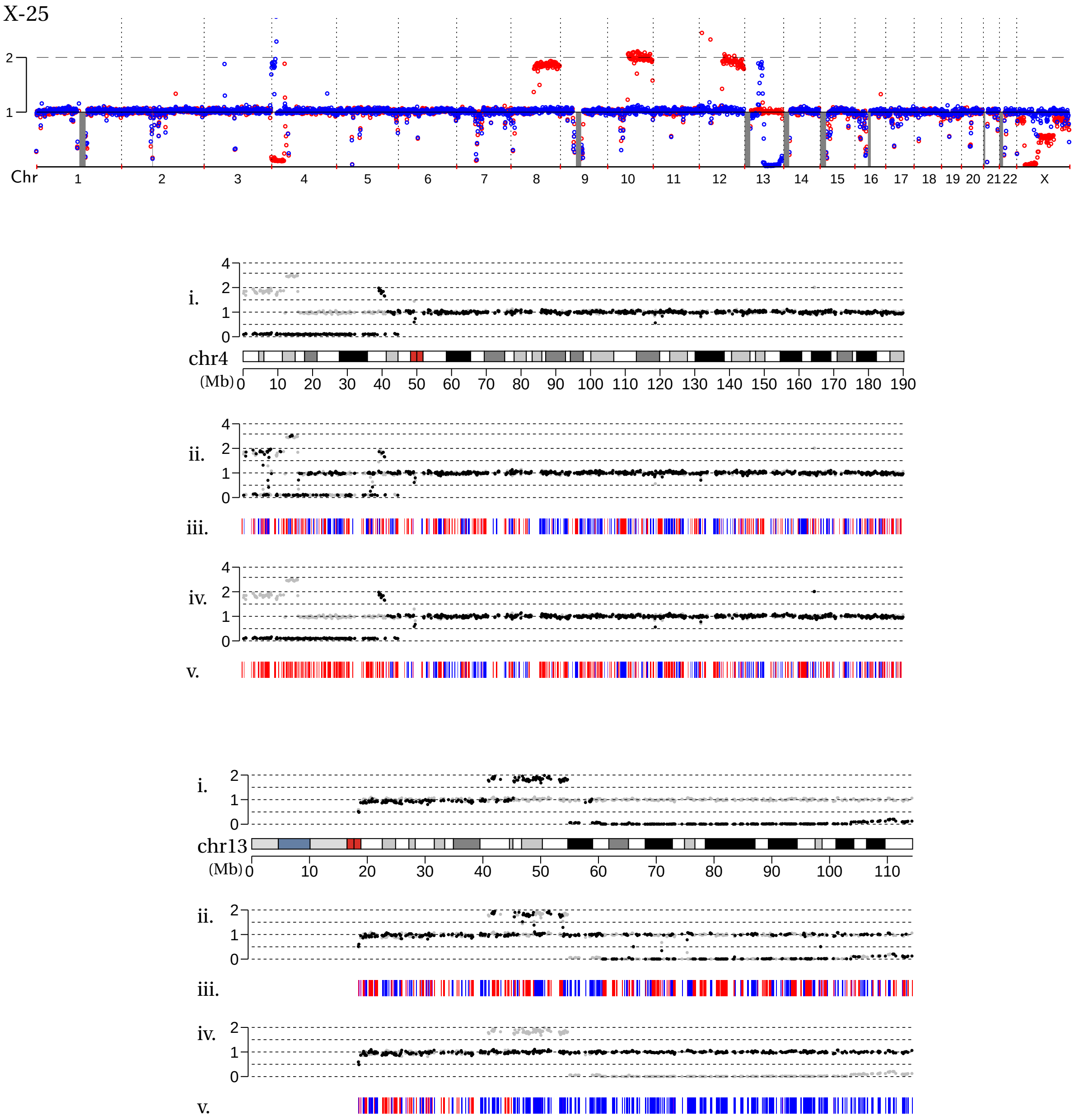


**Fig. S10.** Examples of haplotype inference based on changes in the allelic depths of both homologs (Chr.4) and multiple regions of allelic imbalance on a single homolog (Chr.13). Both examples are from the X-25 clone after telomere crisis. The top panel shows genome-wide DNA copy number of both homologs (red and blue) that are calculated using the truth haplotype. The lower panels show results of allelic-imbalance based haplotype correction. Both examples validate the correct haplotype inference in regions of allelic imbalance and further demonstrate the capability to resolve complex segmental alterations. For Chr.4p, the requirement that each allelic depth changepoint only affects one homolog (Fig. S3i) is sufficient to determine the 4p haplotype despite SCNAs affecting both homologs. For Chr.13q, the phasing of interspersed segmental gains and the terminal deletion to a single homolog reflects the assumption that segmental copy-number changes more often arise on a single homolog (Fig.S3ii, middle) and is also consistent with the outcomes of sequential breakage-fusion-bridge cycles (see Fig. S4iii and its caption); moreover, the phasing of both alterations to a single homolog is directly verified by the truth haplotype data. Note that the switching errors in the gained segment (red bars) are restricted to regions of allelic imbalance between interspersed segmental gains; such errors do not impact the phasing and interpretation of segmental copy-number alterations.

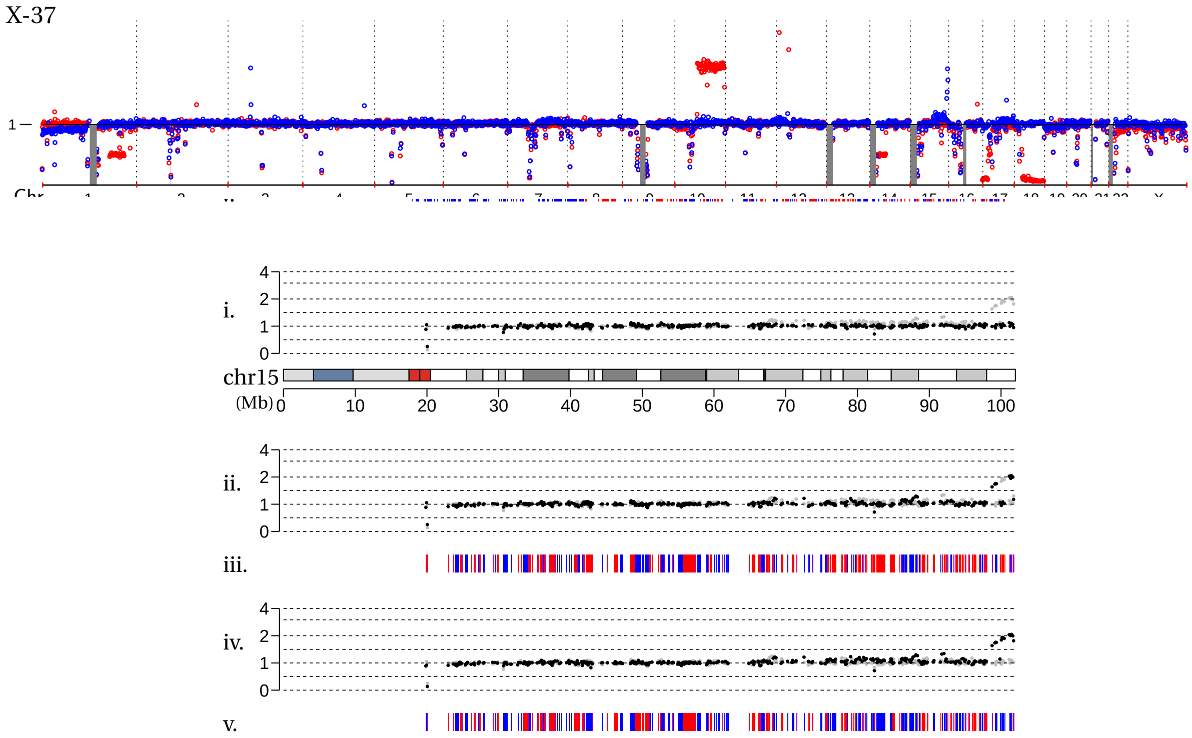


copy-number homolog) and near the 6q terminus.

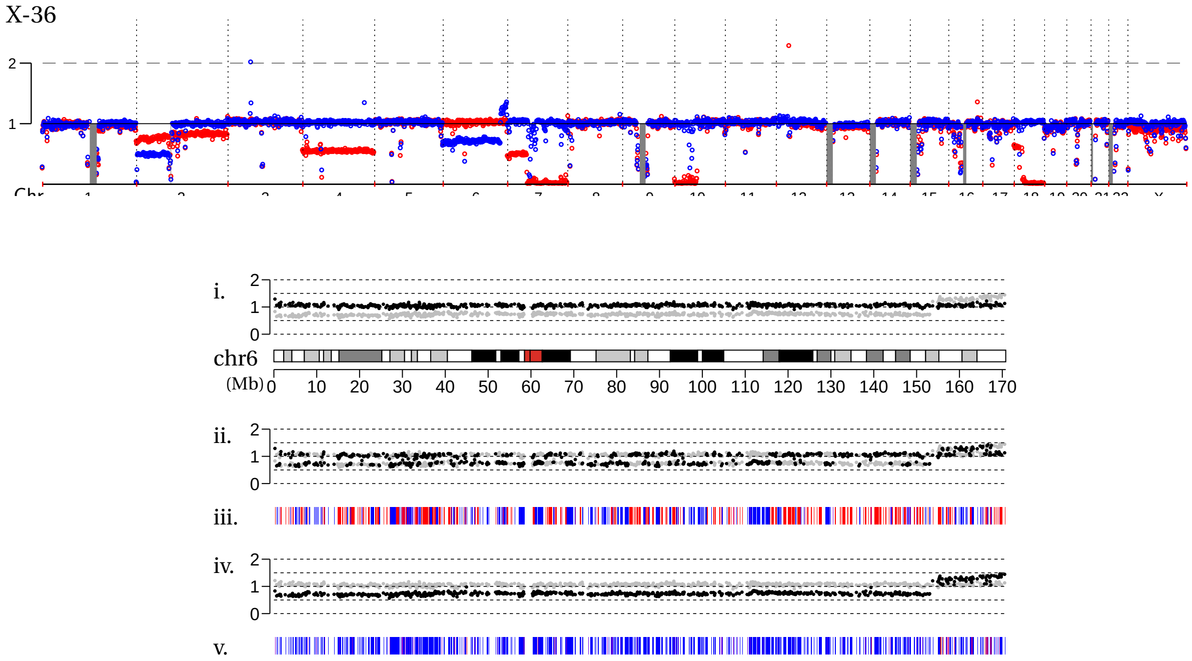


**Fig. S11.** Examples of sloping copy number variation on a single haplotype in two post-crisis RPE-1 clones X-37 (Chr.15) and X-36 (Chr.6). The plots are the same as in Fig.S10. Note the correct phasing of sloping copy number on one homolog and constant copy number on the opposite homolog. This result not only validates the correct haplotype inference based on allelic imbalance but provides further evidence that telomere crisis or breakage-fusion-bridge cycles can generate sloping copy-number variation.

**Additional data S1 (DownSamplePlots.pdf)**

Validation of allelic depth coverage calculation from low-pass whole-genome sequencing

**Additional data S2 (AllelicDepthPhasing.pdf)**

Validation of allelic-imbalanced based haplotype inference
