## Supplementary material for "Genomic signatures of past and present chromosomal instability in the evolution of Barrett’s esophagus to esophageal adenocarcinoma": Methods and Captions for Supplementary Figures and Tables

**Online Methods**

**Extended Data Figure and Table legends**

**Additional References**

**Extended Data Figure 1-10 (uploaded separately)**

**Extended Data Tables 1-8 (uploaded separately)**

**Downloadable Supplementary Data**

https://www.dropbox.com/sh/ib55n6fkacz9jds/AACHxVIHMOsFykiAte3ShkOba?dl=0

BE-EAC-CopyNumberAnalysis: Coverage data and haplotype data

BE-EAC-CopyNumberData: Haplotype-specific copy-number plots of bulk BE/EAC samples

BE-EAC-RearrangementData: Rearrangement data

BE-EAC-ShortVariants: Single-nucleotide variants and insertion/deletion alterations

Single-BE-Cell-CopyNumber: Haplotype-specific copy-number plots of single BE cells

Re-analysis.Killcoyne.NatMed.2020: Normalized sequence coverage of longitudinal BE sequencing

### METHODS

#### Sample identification, DNA extraction, and sequencing data generation

***Sample identification***

With documented informed consent and institutional IRB approval, formalin fixed paraffin embedded (FFPE) endoscopic mucosal resections or esophagectomy samples were identified in the pathology archives of the Mayo Clinic, Brigham and Women's Hospital, or the University of Pittsburgh Medical Center. Patients having received either chemo/radiotherapy or endoscopic ablation prior to resection were excluded. Hematoxylin and eosin (H&E) stained slides were reviewed by two gastrointestinal pathologists (M.S. and A.A.) to determine consensus areas of Barrett's Esophagus (BE), BE with low grade dysplasia (LGD), BE with high grade dysplasia (HGD), and esophageal adenocarcinoma (EAC). (See **Extended Data Figure 1** for examples.) In cases of uncertainty of pathological classification, the samples were reviewed by a third pathologist (R.O.). Any sample without a consensus diagnosis was excluded from further analysis.

***DNA isolation and sequencing library construction***

Ten 4μm sections from each FFPE block were cut sequentially onto PEN membrane frame slides (Life Technologies, Grand Island, NY) bracketed by standard slides for H&E staining. The frame slides were stained using Arcturus paradise plus stain (Life Technologies) following the manufacturer's recommendations. The areas of interest were microdissected using the ArcturusXT laser capture microdissection Instrument (Life Technologies). When dissecting normal tissue that was used as germline reference, we avoided epithelial tissue that may contain BE or EAC cells.

DNA was isolated using the Promega (Madison, WI) FFPE DNA isolation kit following the manufacturer's protocol with the exception that the tissue was digested with proteinase K overnight. DNA was quantified using Picogreen dsDNA Quantification Reagent. Approximately 50ng of genomic DNA was fragmented by sonication (Covaris) to 250 bp and further purified using Agentcourt AMPure XP beads. Whole-genome DNA libraries were constructed from size-selected DNA using KAPA HTP Library Preparation Kit (Roche) and subjected to low-pass whole-genome sequencing (~0.1x). Samples with sufficient library complexity (i.e., estimated total number of unique sequencing fragments ≥ 100 million) were selected for deeper whole-genome sequencing (20-30x).

***Sequencing data generation and processing***

Multiplexed whole-genome sequencing (WGS) libraries were sequenced on NovaSeq6000 or HiSeq2500 instruments (Illumina) in paired-end mode (2 x 150bp). Sequencing reads were aligned to the NCBI Human Reference Genome Build GRCh37/hg19 using `bwa` (version 0.7.7). Aligned reads were processed using the standard pipeline established by the Genomics Platform at the Broad Institute, including base-quality score recalibration, duplicate removal, and realignment near indel variants as described in the GATK best practice (https://gatk.broadinstitute.org/hc/en-us/articles/360035535912).

### Generation of single-cell sequencing data

#### *Cell sorting*

Cells were harvested by endoscopic cytology brushing from a region of high-grade dysplasia and then pelleted in a falcon conical tube (Stem Cell) after trypsin digestion and washing with Dulbecco's phosphate-buffered saline (DPBS, Gibco). Freshly prepared, cold 70% ethanol (5ml) was added drop-wise while vortexing to fix and preserve cells at -20$^{o}$C. Cells were stained by DAPI (Life Technologies) and underwent fluorescence-activated sorting (FACS) into a skirted RNase, DNase-free 96-well plate (Eppendorf) with 5μl DPBS added to each well before sorting. During sorting, the first (A1) well was left empty and the last well (H1) contained 100 sorted cells, both serving as controls for single-cell genome amplification. Each plate after sorting was immediately sealed and placed on dry ice before transferred to -80$^{o}$C storage.

#### *Library construction*

Single-cell lysis and whole-genome amplification was performed using the REPLI-g Single Cell kit (Qiagen) with the following modifications: Due to having 5μl of instead of the standard 4μl starting solution, we added 3.5μl (instead of 3μl) of Buffer D2 for cell lysis and 3.5μl (instead of 3μl) of Stop Solution to terminate cell lysis. During genome amplification, we added 7μl kit water instead of 9μl into the master mix (38μl master mix was added into each well) to match the total volume at the end of reaction. Amplified DNA was purified with ethanol and quantified by Qubit dsDNA HS Assay kit (Life Technologies). About 100ng amplified DNA was sheared to ~350 bp DNA fragments (Covaris sonication) and processed with a KAPA HTP Library Preparation Kit (KK8234, KAPA Biosystem) for multiplexed Illumina sequencing library construction.

#### *Quality assessment and sequencing of single-cell libraries*

A total of 95 whole-genome amplified DNA libraries (94 single cells and one 100-cell sample) were screened by low-pass MiSeq sequencing, from which we identified 24 cells with discernable arm-level copy-number changes and 45 cells with close to diploid coverage. The remaining samples showed poor coverage uniformity and were discarded. Aneuploid cells (24 total), diploid cells (45 total), and a 100-cell sample were pooled and sent for paired-end sequencing (50bp x 2) on the NovaSeq6000 platform (S2 kit) to yield 2.9 billion read pairs (2.15 billion aligned), reaching about 1x mean coverage per cell. The sequencing data were aligned to the GRCh38 reference by `bwa`.

### Detection and filtering of somatic and germline single-nucleotide variants

#### *Mutation detection with Mutect2*

We first performed short variant discovery in each BE/EAC sample using GATK Mutect2 (version 4.0.1.2) [**m1**] and the matching germline reference as control. To filter false variants due to recurrent alignment

errors, we used a ‘reference’ panel of variants detected in 125 blood samples:
**gs://fc-16adb3e5-7c0a-4805-aa5e-374b579d03e1/wgs_hgx19_125_cancer_blood_normal_panel.vcf**
To filter rare artifacts and germline variants that were missed in the matching germline reference, we built a germline resource consisting of >10,000 genomes from gnomAD (version 2.0.2) [**m2**]. To remove low-confidence variants, we applied the following downstream filters: 8-oxoguanine (OxoG) artifacts, FFPE artifacts and alignment artifacts due to sequence similarity between two or more regions in the genome.

Commands for filtering OxoG and FFPE artifacts:

```
gatk FilterByOrientationBias \
   -V filtered.vcf.gz \
   --artifact-modes 'G/T' \
   -P tumor_artifact.pre_adapter_detail_metrics.txt \
   -O oxog_filtered.vcf.gz
```

Commands for filtering alignment artifacts:

```
gatk FilterAlignmentArtifacts \
   -R hg19.fasta
   -V somatic.vcf.gz \
   -I somatic_bamout.bam \
   --bwa-mem-index-image hg38.index_image \
   -O filtered.vcf.gz
```

The filtered variants were annotated using Oncotator (version 1.9.9.0) [**m3**]. We genotyped mutations detected in individual samples across all samples from each patient by running Mutect2 in the `GENOTYPE_GIVEN_ALLELES` mode. We considered the mutant allele to be present in a sample if there were at least three variant-supporting reads; we then used the genotype data to calculate the pairwise similarity between samples that is shown in **Extended Data Figure 2**.

***Joint variant detection by HaplotypeCaller***

We used HaplotypeCaller (GATK HaplotypeCaller v.4.0.12.0-6 [**m5**]) to detect both germline heterozygous variants and somatic variants jointly from all samples from each patient. To filter false variants due to recurrent alignment errors, we imposed the following read filters:

`--minimum-mapping-quality 30` (excluded reads having low mapping quality)

`--read-filter MateOnSameContigOrNoMappedMateReadFilter` and

`--read-filter MateDifferentStrandReadFilter` (discordant alignment positions)

`--filter-too-short 25` (excessive clipping)

`--read-filter OverclippedReadFilter` (over soft-clipping)

`--read-filter GoodCigarReadFilter` (bad CIGAR string)

`--read-filter AmbiguousBaseReadFilter` (>5 percent of N bases in the sequence). We selected only bi-allelic variants and further removed variants in low-complexity DNA sequences [**m6**], poorly mappable regions of the genome [**m7**], or within 100 base pairs of INDEL, multinucleotide changes, or other variants (`bcftools filter --SnpGap 100:indel,mnp,other,overlap`).

To select heterozygous variants for haplotype-specific copy-number calculation, we imposed the following criteria on biallelic SNVs detected by HaplotypeCaller from all samples (both germline reference and BE/EAC) in each patient: (1) variant sites were among common polymorphisms in the 1000 Genomes Project Phase 3 reference haplotype panel (only these variants were used for statistical haplotype phasing); (2) at least one sample showed the heterozygous genotype ('0/1'); (3) at least two samples showed more than two reads of the alternate genotype; (4) at least two samples showed more than two reads of the reference genotype. We further excluded variants in autosomes that were heterozygous in >50% of samples in our cohort (8/15) based on the expectation that the frequency of heterozygotes in a population following the Hardy-Weinberg equilibrium should be <50%. All these filters served to remove homozygous variants that appeared to be heterozygous due to sequencing errors, alignment errors, or technical artifacts in FFPE libraries.

To improve the detection sensitivity of cancer gene mutations, we also ran HaplotypeCaller on the cancer gene consensus [**m4**] plus three genes (*GATA4*, *GATA6*, *VEGFA*) that are recurrently altered in esophageal cancers. This analysis revealed recurrent loss-of-function mutations in *TP53*, *CDKN2A*, *ARID1A*, *ARID1B*, and *SMARCA4* that are annotated in **Figure 2**.

**Standard copy-number analysis and estimation of sample purity and ploidy**

We performed standard somatic copy number analysis using the GATK4 Somatic CNV ModelSegments pipeline (version 4.0.1.2) [**m8**]. Briefly, read counts were collected in 5kb genomic intervals, normalized to fractional coverage, and then corrected for GC-dependent bias. Recurrent coverage bias in FFPE libraries was then normalized using the sequence coverage of germline samples in our cohort as a reference panel. The normalized total sequence coverage and allelic ratio (estimated from allelic depths at heterozygous variant sites) were used as input to ModelSegments for smoothing and segmentation with the following changes to default parameters. To filter out low quality data points, we increased 'minimum-total-allele-count' to 50 (default: 30); to avoid over-segmentation, we increased 'number-of-changepoints-penalty-factor' to 1.8 (default: 1.0). We further calculated average normalized sequence coverage in 25kb genomic intervals for haplotype-specific copy-number analysis.

We estimated the clonal fraction ('purity) and average DNA copy number ('ploidy') of aneuploid BE/EAC clones using ABSOLUTE (version 1.5) [**m9**]. Input data to ABSOLUTE include: (1) normalized read depth and allelic ratio in 5kb bins; (2) segmented copy ratio; (3) allelic frequency of somatic mutations.

We manually reviewed all candidate solutions generated by ABSOLUTE to pick the optimal solution with the fewest subclonal (non-integer) copy-number states. In selecting the most likely solution, we further took into consideration the tumor purity assessed from histopathological analysis. The purity and ploidy estimates were later validated independently by haplotype-specific sequence coverage. BE samples without large SCNAs were excluded from purity/ploidy estimates: Their phylogeny was inferred from sSNVs or small focal SCNAs (Patient1: COLME and BE; Patient 4: COLME; Patient 5:BE; Patient 7:COLME). An exception was the HGD1 sample in Patient 9. This sample contained no large segmental SCNAs but several regions of focal amplification: The amplified copy number was most likely due to tumor cells from the adjacent IMEAC2 lesion (see **Figure 2)**, which was supported by the lack of amplified DNA in HGD1 from cytogenetic analysis (**Extended Data Table 7**,Tab 3). The HGD1 sample was placed next to the IMEAC2 sample in the phylogenetic tree based on this feature.

**Haplotype-specific copy number analysis**

The idea of using haplotype information to improve the accuracy of allelic fraction calculation was previously implemented for SNP array data[1], whole-genome sequencing[2], and whole-exome sequencing[3]. Our haplotype-specific copy-number analysis workflow (**Supplementary Figure 1**) combines statistical phasing (**Supplementary Figure 2**) and allelic-depth based phasing[1] (**Supplementary Figure 3**) to extend the range of haplotype inference and SCNA phasing to entire chromosomes (or arms). The ability to identify somatic copy-number alterations on each parental chromosome (**Supplementary Figure 4** and **5**) further enables us to determine the relationship between SCNA breakpoints (**Supplementary Figure 6)** and relate copy-number evolution patterns in BE/EAC genomes to the copy-number outcomes of unstable chromosomes.

***Identification of polymorphisms in parental chromosomes***

The identification of heterozygous variant sites on parental chromosomes was described in "**Joint variant detection by HaplotypeCaller**". Because BE/EAC samples also contain DNA from normal cells, joint variant detection from both BE/EAC samples and the matching germline reference achieves better variant detection sensitivity than variant detection solely from the germline reference, especially for germline samples with low sequencing coverage (<15x in Patients 8-11). The joint detection strategy consistently reveals 1.5-1.7 million common heterozygous variants (identified in the 1000-genome project cohort) in all 15 patients and 1.1-1.3 million variants in each individual sample (**Extended Data Table 1**). The high density of heterozygous variants (~1 per 3kb) ensures the accuracy of allelic copy-number calculation.

***Statistical phasing of parental haplotypes***

The heterozygous genotypes in each patient were uploaded to the Sanger Imputation Server for statistical phasing using EAGLE2[4] (version 2.0.5) [**m10**] and reference haplotype data from the 1000-Genome Phase

3 release [**m11**]. Although EAGLE2 can directly perform statistical phasing using both heterozygous and homozygous genotypes, based on benchmarking using reference haplotype data[5], we found that the haplotype phase calculated using only heterozygous genotypes was slightly more accurate than the haplotype phase inferred from both heterozygous and homozygous genotypes. We therefore used the haplotype derived from statistical phasing applied to only high-confidence heterozygous variant sites.

***Calculation of haplotype fraction and allelic sequence coverage***

To calculate phased average allelic fractions, the allelic depths of heterozygous genotypes (**Suppl Fig. 1 ii**) were first converted to haplotype-specific (A or B) allelic depths based on the phase of genotypes (A or B) (**Suppl Fig. 1.ii**). Based on the allelic fractions $F_A(x)$ and $F_B(x)$, and the normalized sequence coverage $R(x)$ (described in "**Standard copy-number analysis and estimation of sample purity and ploidy**"), we then calculated local allelic sequence coverage as $A(x) = R(x)F_A(x)$ and $B(x) = R(x)F_B(x)$. (Note that the total sequence depth $R(x)$ is centered at 2 such that $A(x)$ and $B(x)$ center at 1 in a diploid genome.) These data are plotted in "`xxx_phased.pdf`" files in the **Downloadable Supplementary Data** and used for downstream copy-number calculation.

The combination of normalized total sequence coverage $R(x)$ and phased average allelic fractions $F_A(x)$ and $F_B(x)$ has two advantages. First, as the total sequence coverage $R(x)$ is calculated using both phased reads (i.e., those overlapping with heterozygous sites) and unphased reads, it shows less variability than the combined allelic depths at heterozygous sites. Second, the phased average of allelic fractions at multiple adjacent variant sites reduces fluctuations of allelic fractions at single variant sites (**Suppl Fig. 1**). To see how the phased average improves the accuracy of allelic fraction calculation, we consider the distribution of allelic depths from $D$ copies of genomic DNA from both homologs (i.e., mean sequencing depth = $D$) with allelic fraction $f$ and $1 - f$. The mean allelic depths are given by $Df$ and $D(1 - f)$. Assume the allelic depths follow binomial distributions, the variance of allelic depths at a single variant site is $Df(1 - f)$ and the variance of allelic fraction at a single variant site is $f(1 - f) / D$. As the fluctuations of allelic fractions at different variant sites are largely independent, the variance of the average allelic fraction over $n$ variant sites is approximately

$$f(1-f)\sum_i \frac{1}{D_i} = \frac{f(1-f)}{n}\left\langle \frac{1}{D} \right\rangle \sim \frac{1}{n}$$

where $D_i$ is the total allelic depth at site $i$. Therefore, the statistical error of the average allelic fraction over $n$ variant sites is reduced by $1/\sqrt{n}$ relative to the statistical error of allelic fractions at single variant sites. Importantly, the average allelic fractions need to be calculated based on the parental haplotype but not the major or minor allelic fractions as the mean of major or minor allelic fraction is not equal to the allelic fraction of parental chromosomes. For example, let $F(x)$ denote the observed allelic fraction of haplotype A, then the major allelic fraction is given by $\max[F(x), 1 - F(x)] = |F(x) - 1/2| + 1/2$. It is straightforward to see that its expectation $E[|F(x) - 1/2| + 1/2] > E[F(x)] = f$.

Although the phased average allelic fraction displays less variation than the allelic fraction at single variant sites, the averaging will reduce the resolution of allelic fraction changepoints and more importantly, reduce the signal of allelic imbalance when there are switching errors in the haplotype phase[5]. The rate of switching errors in statistical phasing was estimated to be 0.1-1% [4], or one switching error every 0.1-1Mb. To avoid inaccurate allelic fraction calculation due to haplotype switching, we performed phased average allelic fractions in 25kb genomic intervals with five or more variants. In our cohort, the lowest sequence depth is ~12, and each sample has at least 1 million variants with sequence coverage (~8 per 25kb). Based on these numbers, the average statistical error of phased allelic fractions in 25kb intervals is lower than

$$\sqrt{\frac{f(1-f)}{n}\left\langle\frac{1}{D}\right\rangle} \approx \sqrt{\frac{0.5^2}{5}\cdot\frac{1}{12}} = 0.06.$$

We further note that (1) the 25kb window sets the resolution of copy-number changepoints; (2) although one can theoretically detect SCNAs down to 25kb, the real sensitivity will depend on both the clonality and the size of SCNAs.

***Validation of purity/ploidy estimates using allelic sequence coverage***

Let $c_A(x)$ and $c_B(x)$ denote the (integer) DNA copy number of parental chromosomes, $\alpha$ denote the clonal fraction of tumor cells and $\tau$ denote the average ploidy of the tumor genome, we have

$$R(x) = 2 \times \underbrace{\frac{\alpha\,(c_A(x) + c_B(x)) + 2(1-\alpha)}{\tau\alpha + 2(1-\alpha)}}_{\text{local read depth ratio}},$$

$$F_A(x) = \frac{\alpha c_A(x) + (1-\alpha)}{\alpha\,(c_A(x) + c_B(x)) + 2(1-\alpha)}, \quad F_B(x) = \frac{\alpha c_B(x) + (1-\alpha)}{\alpha\,(c_A(x) + c_B(x)) + 2(1-\alpha)},$$

where the factor of 2 in $R(x)$ is introduced such that the allelic depths

$$A(x) = R(x)F_A(x) = \frac{\alpha c_A(x) + (1-\alpha)}{\alpha\cdot\tau/2 + (1-\alpha)}, \quad B(x) = R(x)F_B(x) = \frac{\alpha c_B(x) + (1-\alpha)}{\alpha\cdot\tau/2 + (1-\alpha)} \qquad (1)$$

center at 1 in a diploid genome.

As the tumor clonal fraction $\alpha$ and ploidy $\tau$ are both constant and $c_A(x)$ and $c_B(x)$ both take integer values, the allelic sequence coverage $A(x)$ and $B(x)$ should both cluster around discrete values (**Suppl Figure 4**) starting at $(1 - \alpha)/(\tau\alpha + 2(1 - \alpha))$ corresponding to allelic copy-number state of 0, with increments of $\alpha/(\tau\alpha + 2(1 - \alpha))$ reflecting a single copy gain. The clustering pattern of allelic sequence coverage can therefore be used to verify estimates of tumor clonal fraction and ploidy. We generated scattered plots of $A(x)$ and $B(x)$ of 25kb intervals for every chromosome in each sample in "`xxx_CNstates.pdf`" in the **Downloadable Supplementary Data**. We used the allelic coverage data to validate ploidy and purity estimates calculated by ABSOLUTE.

***Haplotype refinement and phasing of somatic copy-number alterations using allelic imbalance***

In regions with allelic imbalance, the high and low allelic coverage should reflect the DNA copy number of parental chromosomes[1]. The oscillation of allelic coverage in these regions (**Suppl. Figure 4**) is caused by switching between parental haplotypes from statistical phasing. To distinguish between allelic coverage oscillation due to haplotype switching and allele-specific SCNAs, we have implemented a computational workflow to correct switching errors and phase SCNAs based on the allelic coverage (**Suppl. Figure 3**). The workflow consisted of four steps.

1. We first identified regions of allelic balance (5Mb) to estimate random fluctuations of allelic fractions. Deviation from allelic balance (allelic imbalance) can be assessed either from the mean minor allelic fraction (~0.5) or from the standard deviation of phased allelic fractions. For FFPE data with significant coverage non-uniformity, the allelic fractions display significant variation even in disomic regions; we found the standard deviation of phased allelic fractions to be a better classifier of regions of allelic balance. As the primary goal of this step is to derive a 'null' distribution of allelic fractions in regions of allelic balance, we set a stringent cutoff (≤ 0.07 for most samples) that serves to exclude large regions of allelic imbalance but not necessarily include all regions of allelic balance.

2. We next identified regions of allelic imbalance based on the reference allelic fraction distribution determined from regions of allelic balance. Due to uneven sequence coverage and the modest sequencing depths (~10x) or low tumor clonal fractions (< 0.4) of some of the samples, the allelic fractions in regions of allelic balance can display large fluctuations from the mean allelic fraction of 1/2. To discern true allelic imbalance from random fluctuations, we considered allelic fractions in multiple adjacent 25kb intervals. For $m$ consecutive intervals, let $p_i$ ($1 \le i \le m$) denote the likelihood of allelic balance in each interval (two-sided $p$-value of the observed allelic fraction relative to the reference distribution), we can approximate the log-likelihood of allelic balance for the observed allelic fractions in $m$ intervals as

$$L_0 = \sum_{1 \le i \le m} \ln p_i .$$

For true allelic imbalance due to gain or loss of DNA from one parental chromosome, we expect the phased allelic fractions to be either lower or higher than 1/2, i.e., with a unidirectional change; by contrast, random fluctuations should be bidirectional. This difference can be quantified from the frequency of allelic fraction transitions from above to below 1/2 or vice versa. Let $\delta$ be the probability that the allelic fraction transitions across 1/2 between adjacent intervals. Assume no correlation between the allelic fractions in adjacent intervals, we have

$$\delta = \min\left[p(\mathrm{AF} > 0.5), p(\mathrm{AF} < 0.5)\right].$$

In regions of allelic balance, we expect $\delta = 1/2$; In regions with allelic imbalance, we expect $\delta < 1/2$ as the mean allelic fraction is either above or below 1/2. For $m$ intervals with a total of $m$ - 1 transitions, the log-likelihood of seeing $s$ transitions is given by

$$\ln\left[\frac{(m-1)!}{s!(m-1-s)!}\delta^s(1-\delta)^{m-1-s}\right]$$
$$\approx (m-1-s)\ln(1-\delta) + s\ln\delta - (m-1-s)\ln\left(1-\frac{s}{m-1}\right) - s\ln\frac{s}{m-1}.$$

We can introduce a log-likelihood function for allelic imbalance as

$$L_1 = \underbrace{(m-s)\ln(1-\hat{\delta}) + s\ln\hat{\delta}}_{\text{allelic imbalance}} - \underbrace{m\ln 2}_{\text{allelic balance}}.$$

The first term is from the log-likelihood of the observed number of transitions under the assumption of allelic imbalance ($\hat{\delta} < 1/2$) and the second (constant) term is from the log-likelihood of allelic balance ( $\delta = 1/2$). The binomial coefficients cancel out from the subtraction. Combining the deviation ($L_0$) and the correlation ($L_1$) of allelic fractions, we can define a heuristic score of allelic imbalance based on allelic fractions in $m$ consecutive intervals as

$$L = \sum_{1\le i\le m} \ln p_i - \left[(m-s)\ln(1-\hat{\delta}) + s\ln\hat{\delta}\right].$$

We chose $m$ = 10 such that there is on average less than one switching error across the adjacent intervals (250kb) from statistical phasing. We operationally set $\hat{\delta} = 0.1$. For each 25kb bin, we calculated the allelic imbalance score based on the allelic fractions in 10 adjacent bins. To determine the cutoff for calling allelic imbalance, we compared the distribution of allelic imbalance scores in regions of allelic balance against the distribution in selected regions of allelic imbalance. The cutoffs were set individually for each sample and chosen to be permissive in order to identify subclonal SCNAs in low-purity tumor samples.

3. We next performed allelic-depth phasing on all 25kb bins of allelic imbalance on each chromosome. We concatenated all 25kb bins that were determined to be in allelic imbalance and assessed potential haplotype switching between adjacent bins causing a swap of allelic depths (see "**Calculation of haplotype fraction and allelic sequence coverage**"). The general idea is illustrated in **Suppl. Figure 3** and examples are shown in **Suppl. Figure 4**. For each interval $i$, we compared the allelic depths ($A_i$, $B_i$) in this interval with the average allelic depths ($A$, $B$) of the left 40 intervals (~1Mb) with allelic imbalance and calculated

$$\underbrace{\min(|A_i - A|, |B_i - B|)}_{\text{no swap}} - \underbrace{\min(|A_i - B|, |B_i - A|)}_{\text{swap}}.$$

If the differential metric was ≤ 0.05, we kept the haplotype phase of this interval; if the differential was above 0.05, we switched the haplotype from the $i$th interval to the end of the chromosome. The same procedure was then performed from the last interval to the first by comparing allelic depths in each interval to the right 40 bins. Allelic-depth phasing was performed individually and the results are plotted in "`xxx_ADCorr.pdf`" in **Downloadable Supplementary Data**.

4. Finally, we combined haplotype corrections from individual samples from each patient to derive the consensus parental haplotype. This step both improved the accuracy of local haplotype linkage by aggregating allelic depths from all samples with allelic imbalance and generated longer contiguous haplotypes by concatenating haplotypes determined from allelic imbalance in different samples. The latter was especially powerful given the prevalence of both arm-level and segmental allelic imbalance in BE/EAC genomes. Examples of joint haplotype inference from allelic imbalance in all samples are shown in **Suppl. Figure 5**. The final jointly inferred haplotype still contained occasional long-range switching errors (usually 1-2 switching errors on 1-3 chromosomes per individual). These errors were corrected manually. Readers can refer to codes provided in the repository (see "**Code Availability**") for both the implementation of algorithms in each step of the calculation and the final manual curation. Using the jointly inferred haplotype, we re-calculated haplotype-specific DNA copy number in 25kb intervals. These results are plotted in "`xxx_final.pdf`" in **Downloadable Supplementary Data**.

We want to add a few comments about allelic-depth based phasing. First, this strategy is restricted to regions of allelic imbalance and is necessary because haplotype mixing will average out the difference between the allelic fractions of parental chromosomes, which reduces the signal of allelic imbalance. In regions of allelic balance, the allelic copy number is simply half of the total copy number and haplotype correction is unnecessary. Second, the single-sample allelic-depth phasing algorithm assumes *a priori* that multiple SCNAs on a chromosome are more often accumulated on a single parental homolog than both homologs (**Suppl. Fig. 3ii**, middle). This assumption is based on the biological insight that it is more likely to accrue multiple SCNAs on a single unstable chromosome, either from catastrophic events such as chromothripsis or through multigenerational processes such as breakage-fusion-bridge cycles, than to acquire SCNAs on both homologs through independent DNA breaks. For segmental SCNAs interspersed with regions of allelic balance (**Suppl. Fig. 4iii**), this assumption cannot be directly validated. However, we were able to validate the long-range phasing of SCNAs either by rearrangements joining distal copy-number changepoints (**Extended Data Fig. 8F**) or using chromosome-scale haplotypes derived from whole-chromosome (**Suppl. Fig. 5i**) or large overlapping regions of allelic imbalance (**Suppl. Fig. 5ii,iii**). Finally, even when both parental chromosomes acquire SCNAs (**Suppl. Fig. 4iv, 5ii,iii**), sometimes even related to each other (Patient 7 Chr.12, **Extended Data Fig. 8G**), we were still able to separately phase SCNAs on each parental chromosome based on rearrangements and allelic copy-number changes at SCNA changepoints. The solution of these more complex cases required manual review.

***Segmentation of haplotype copy number***

For each BE/EAC sample, we first calculated the integer DNA copy number $c_A(x)$ and $c_B(x)$ of parental chromosomes in 25kb intervals from the haplotype-specific sequence coverage $A(x)$ and $B(x)$, the clonal fraction $\alpha$, and the ploidy $\tau$ of the major BE/EAC clone using Eq.(1). We then segmented $c_A(x)$ and $c_B(x)$

by an iterative procedure described previously for single-cell DNA data that only have integer copy-number states [**m12**]. Briefly, in each round of iteration, the 25kb copy number was re-calculated as the local median of 9 consecutive bins (0.23 Mb) and then rounded to the nearest integer; the iteration ended when the copy number of all bins converged to integer states. We then collected all copy-number changepoints and re-calculated the segmental copy-number states. The final segmented (using 25kb bins) and 100kb-average integer DNA copy number data are plotted in "`xxx_final_100kb.pdf`" in **Downloadable Supplementary Data**, with annotations of all SCNAs. Whole-genome duplication status was assigned if there were more parental chromosomes with duplicated copy number (CN=2) than single copy (CN=1).

Because we constrained copy-number changes to integers, the minimum detectable SCNA was dependent on both the clonality and the copy-number difference across changepoints. For clonal single-copy changes (i.e., one copy loss or gain), the minimum CNA length was ~0.12 Mb (five bins); for clonal two-copy changes (rare), the minimum CNA length was ~0.08 Mb (3 bins). As this procedure converged to only integer copy-number states, it would incorrectly segment non-integer copy-number states due to incorrect purity/ploidy estimates or in regions of subclonal SCNAs. Non-integer subclonal copy-number states were easily identifiable and annotated in the final copy-number plots; subclonal SCNAs were not used for phylogenetic inference except where stated.

***SCNA Classification and evolutionary inference***

We classified SCNAs on each parental chromosome based on the number of SCNA breakpoints and copy-number states. See **Extended Data Figure 4** for the criteria and examples for each SCNA category. SCNAs affecting the same parental chromosome in different samples were manually reviewed to determine their evolution history. SCNA breakpoints in two or multiple samples that were within 0.1Mb from each other (to account for segmentation inaccuracy) and associated with the same type of copy-number change (either gain or loss) were classified as identical. SCNA breakpoints that were within 0.1Mb but associated with opposite copy-number changes (i.e., copy-number gain in one sample and loss in another sample) were classified as complementary. Individual SCNAs (including complex SCNAs with multiple breakpoints) were classified as shared between two samples if all SCNA breakpoints were identical. If only a subset of SCNA breakpoints were identical, the SCNA patterns were classified as branching (i.e., initiated by a single ancestral event but having different downstream changes). Branching evolution also included examples where distinct SCNA breakpoints on the same parental chromosome can be explained by sequential or progressive DNA alterations (**Fig. 6D and Extended Data Figure 7**).

The timing of SCNA relative to duplication events (both whole-chromosome and whole-genome) was determined as follows. SCNAs associated with copy-number changes above one were assumed to have arisen before duplication; SCNAs with single-copy changes were assumed to have arisen after duplication. For SCNAs identified in samples inferred to have undergone whole-genome duplication (WGD), their timing was further validated based on their presence or absence in related samples from the same patient without WGD acquisition. For chromosomal or arm-level SCNAs, if the final copy-number state was one, they were assumed to have been first duplicated to two copies and then undergone whole-chromosome loss; if the final copy-number state was an odd number above one (3,5,…), they were assumed to have first undergone duplication to the nearest even copy-number state and then undergone either a single-copy gain or a single-copy loss, depending on the number of ancestral WGDs.

**SCNA and SNV-based phylogenetic inference**

Phylogenetic inference was done independently from SCNAs and SNVs. For SCNA-based phylogenetic inference, we used haplotype-specific SCNA breakpoints as linage markers as the breakpoints remain unaltered by downstream whole-chromosome or whole-genome duplication events. The phylogenetic tree was constructed based on the presence or absence of SCNA breakpoints shared by two or more samples. Arm-level or whole-chromosome SCNAs were also considered where there were no shared internal SCNA breakpoints. All phylogenetic trees were manually reviewed to ensure consistency and exclude confounding factors due to (1) subclonal mixture between different samples; (2) whole-chromosome/arm-level deletion that eliminate ancestral copy-number breakpoints.

For SNV-based phylogenetic inference, we calculated genetic similarity as the percentage of shared sSNV variants between two samples normalized by the total number of sSNVs detected in each sample. The sSNV similarity was largely consistent with the SCNA-derived phylogeny with the following discrepancies: (1) HGD lesion in Patient 1; (2) the lineage of HGD2, IMEAC2, IMEAC1 in Patient 12; (3) the lineage of all samples in Patient 15. Evidence supporting the SCNA-derived phylogeny in Patient 12 and 15 was presented in **Suppl. Figure 6**; in both cases, the sSNV similarity was less accurate due to false negative sSNV detection. For the HGD lesion of Patient 1, the dominant clone in HGD was inferred to have undergone whole-genome duplication and contain a missense mutation in *TP53* that was shared with the cancer lesions. Presumably, the HGD lesion was a polyclonal mixture of cells that were similar to BE/COLME (based on the SNV burden) and cells that were similar to EAC1/IMEAC2/IMEAC3; the lesser similarity between HGD and the cancer lesions was also because the cancer lesions had both acquired more *de novo* mutations and lost ancestral mutations in their evolution from the common ancestor with HGD.

**Somatic rearrangement detection**

We performed joint somatic rearrangement detection on all samples from each patient using SvABA (version 1.1.3) [**m13**]. To improve detection sensitivity, we decreased 'mate-lookup-min' to 1 (default: 3) and 'min-overlap' to 25bp (default: 0.4 x read length). We eliminated rearrangements with one or both breakpoints overlapping with known germline variants, blacklisted regions, or regions of low sequence complexity [**m14**]. To eliminate false rearrangements due to chimeric sequences generated in FFPE library construction, we first excluded rearrangements with breakpoints within 100kb and then only included rearrangements with both breakpoints within 100kb from copy-number changepoints. The copy-number filtering strategy inevitably removed true rearrangements without apparent copy-number changes, but was necessary due to the high false positive rate (due to chimeric sequences) and low sensitivity (due to DNA degradation) of rearrangement detection in FFPE libraries. (See **Extended Data Figure 6** for snapshots of both random chimeric sequences near true rearrangements validated by copy-number breakpoints). We therefore restricted the analysis to SCNA-related rearrangements.

**Fluorescence in situ hybridization analysis**

After deparaffinization and dehydration, FFPE tissue sections were first digested in 0.1N HCl for 20-30 minutes and then washed in phosphate-buffered saline (PBS) solution for 5 minutes at room temperature. Bacterial artificial chromosomes (BAC) probes against centromeric sequences (*CEN3*, *CEN4, CEN5, CEN8, CEN10, CEN11, CEN11q, CEN12, CEN13, CEN18, CEN22, CEN17*) and amplified oncogenes (*ERBB2, MYC, EGFR, KRAS, VEGFA,* and *FGFR2*) were fluorescently labeled (Chromosomescience laboratory, Sapporo, Japan). After dehydration and drying, each FISH probe was applied to each targeted area of tissue. The slides were sealed with coverslips, denatured at 90ºC for 10 minutes, and then followed by overnight hybridization at 37°C in a wet chamber. Hybridized slides were washed in 2x saline-sodium citrate (SSC) buffer for 5 minutes and coverslips were removed gently. The slides were washed in 50% formamide/2x SSC for 20 min at 37°C, and then kept in 1x SSC for 15 min at room temperature. The slides were counterstained with 4′,6-diamidino-2-phenylindole (DAPI). The FISH images were captured with a fluorescence microscope (BZ-X710, Keyence, Japan). The number of gene probes and corresponding centromeric probes were then manually quantified.

**Single-cell sequencing analysis**

***Calculation of total DNA copy number***

We calculated total DNA copy number from the sequence coverage of each cell in four steps. (1) Read counts were calculated in 10kb intervals and centered by the genome-wide mean. (2) The average read coverage (10kb) in each sample was then normalized for recurrent coverage bias estimated using the median coverage across all samples. (3) GC-dependent coverage variation was normalized based on %GC in 100kb intervals. (4) The 10kb normalized sequence coverage was averaged over 100 bins (1Mb) to generate local sequence coverage.

When performing step (2) above, we needed to first select a region with constant DNA copy number in each cell. We either picked the largest chromosome arm with median coverage close to the genome-wide median of arm-level median coverage (absolute deviation less than 0.05) or picked the arm with the lowest standard deviation of coverage when no arm was close to the genome-wide median (e.g., highly aneuploid genomes). Normalization of recurrent coverage bias was performed on log-transformed sequence coverage. The (log-transformed) sequence coverage in the selected region of constant DNA copy number was fitted to a cubic polynomial function of the (log-transformed) median coverage across all samples. We used the cubic function to calculate recurrent bias across the genome based on the median coverage. The recurrent coverage bias was subtracted from the (log-transformed) coverage in the original sample, which was then converted to normalized coverage by exponentiation.

***Statistical phasing of parental haplotypes***

We counted reference and alternate allelic depths in each cell using the ASEReadCounter module from GATK4 (version 4.0.1.2) [**m15**] at common SNP sites identified in 1000 Genomes Project Phase 3 reference haplotype panel [**m11**] (lifted to GRCh38). We selected variants with the minor allele observed in ≥ 5 cells as heterozygous variants for which statistical phasing was performed with EAGLE2 (version 2.4.1) [**m16**] using the 1000-Genome Phase 3 reference haplotype panel [**m11**]. The 3p-terminal region (0-33.7Mb) was exceptional as most cells (including the 100-cell sample) showed loss-of-heterozygosity. To identify samples that were (partially) heterozygous, we first estimated the heterozygosity of each sample in this region using the fraction of minor allelic coverage at common variant sites with both genotypes observed in at least one sample. This led us to identify four samples (A2,A3,E1,E7) with estimated heterozygosity > 0.05. We combined these four samples with the 100-cell sample (H1) and selected common variant sites with the minor genotype seen in at least two out of five samples as heterozygous variants. The parental haplotype phase at these variant sites were then derived from the major and minor genotypes from all samples instead of statistical phasing.

ASEReadCounter command:

```
gatk ASEReadCount -R <hg38_ref.fa> \
      -I <reads.bam> -O <allelic_depths.txt> -V <Variant_VCF>
```

with the same read filters for running HaplotypeCaller in bulk samples as described before.

EAGLE2 command:

```
eagle --vcfTarget=<filtered_hets.vcf.gz> \
      --vcfRef=<1000G_hg38_genotypes.bcf> \
      --geneticMapFile=<hg38_genetic_map.txt.gz> \
      --chrom=chr? \
      --outPrefix="phased_hets.vcf.gz" \
      --numThreads=12
```

***Two-pass haplotype correction and allelic copy-number calculation***

Compared to bulk copy number data, single-cell copy-number data have more variability but also display more significant allelic-depth differences in regions of allelic imbalance due to having only integer copy-number changes. To attenuate coverage variability due to amplification, we calculated total DNA copy number in 1Mb intervals and allelic fractions in 50kb intervals. The choice of 50kb instead of 1Mb intervals for allelic fraction calculation was because switching errors in statistical phasing occurs about once per 250kb and will attenuate allelic depth difference in 1Mb intervals in regions of allelic imbalance.

We performed allelic-depth based haplotype correction in two passes. In the first pass, we aggregated allelic-depth differences in all single cells to detect recurrent allelic imbalance and correct switching errors in these regions. After the first pass, we reviewed the copy-number data and identified aneuploid samples with large segmental allelic imbalance. These samples were used for the second round of haplotype correction. The haplotype solution after the second pass was then used to calculate the final haplotype-specific DNA copy number that was plotted in **Downloadable Supplementary Data**.

***Haplotype refinement using allelic imbalance in single cell data***

Due to the variability of single-cell DNA copy number, we used a different strategy to perform haplotype correction from allelic imbalance in single-cell data. Let $c_{A,i}(x)$ and $c_{B,i}(x)$ be the local allelic copy number in sample $i$ at genomic location $x$. For any two bins ($x$ and $y$) in a single chromosome, we expect one homolog to have constant DNA copy number in most cells, *i.e.*,

$$c_{A,i}(x) = c_{A,i}(y) \text{ or } c_{B,i}(x) = c_{B,i}(y).$$

If there is haplotype switching between $x$ and $y$, we expect

$$c_{A,i}(x) = c_{B,i}(y) \text{ or } c_{B,i}(x) = c_{A,i}(y).$$

To assess the likelihood of haplotype switching between $x$ and $y$, we introduced

$$\delta(x,y) = \sum_i \mathbb{1}\left[\min\left(\left|c_{A,i}(x) - c_{A,i}(y)\right|, \left|c_{B,i}(x) - c_{B,i}(y)\right|\right)\right] > \delta,$$

and

$$\delta'(x,y) = \sum_i \mathbb{1}\left[\min\left(\left|c_{A,i}(x) - c_{B,i}(y)\right|, \left|c_{B,i}(x) - c_{A,i}(y)\right|\right)\right] > \delta.$$

$\delta(x,y)$ represents the number of samples where the local allelic copy number states in bin $x$ and bin $y$ show a significant difference (> $\delta$) based on the current haplotype phase and $\delta'(x,y)$ represents the same number with the opposite haplotype linkage between bin $x$ and in bin $y$. In regions of allelic balance, even if there are occasional allelic copy-number differences above the threshold, we expect such differences to be random and cancel out among many cells. By contrast, in regions of allelic imbalance, the gained homolog or deleted homolog will be preserved in multiple cells, producing recurrent allelic bias that can be used for haplotype correction. We therefore used

$$\epsilon(x,y) = \delta(x,y) - \delta'(x,y)$$

as a penalty function for switching errors. $\epsilon(x,y) < 0$ favors the current haplotype linkage between $x$ and $y$ as there are fewer samples showing significant changes in both allelic copy-number states (> $\delta$), and vice versa. Let $s(x) = 1$ represent the haplotype phase of 50kb bins, we iteratively flip $s(x)$, i.e., $s(x) \to -s(x)$ to find the haplotype phase $s(x)$ that minimizes

$$E = \sum_{x,y} \epsilon(x,y) s(x) s(y).$$

It is straightforward to verify that minimization of $E$ produces the correct haplotype in regions with at least one parental chromosome having constant DNA copy number. This assumption was verified as we usually observed at least one constant allelic copy-number state across each chromosome (**Extended Data Table 3**) reflecting an unaltered homolog, even when there were multiple SCNAs on the chromosome.

In the first pass of allelic-depth based phasing, we used local haplotype-specific copy number (50kb bins) in all cells with cutoff penalty $\delta = 0.6$. In the second pass, we only used aneuploid cells with cutoff penalty $\delta = 0.8$.

***Determination of chromosomal copy number***

To determine the integer copy-number state of each chromosome in a single cell genome, we first normalized the average copy number of each chromosome by the median arm-level allelic copy number of both homologs across the genome. For near diploid genomes, the median allelic copy number is 1 and all copy-number states should be integers (0,1,2, …). As the minimum non-zero copy-number state is one, the presence of half integer copy-number states indicates duplication of the remaining chromosomes, i.e., whole-genome duplication; in this scenario, we multiplied the copy number by two to account for whole-genome duplication. We considered a genome to be near tetraploid if there was at least one chromosome arm with median allelic copy number between 0.2 and 0.8 and standard deviation of allelic coverage < 0.25.

***Joint mutation detection in single-cell samples***

We performed joint somatic mutation detection in single cells using the same command line as described above for joint variant calling in bulk samples, with the only difference being the genome reference (GRCh38 instead of GRCh37). Variants were annotated using snpEff [**m17**] with the following command line argument: `-v GRCh38.86 -ud 1000 -onlyProtein -canon`

**Longitudinal BE sequencing analysis**

***Data Processing***

With approval from the International Cancer Genome Consortium, we downloaded and re-aligned sequencing data from a previous study[54] that were available from European Genome-phenome Archive (Dataset ID: EGAD00001006033) with controlled access. The cohort consisted of 773 BE/EAC (602 NDBE/IND, 109 LGD, 37 HGD, and 25 IM/EAC) samples from 88 patients. We performed normalization of 25kb read-depth coverage in all samples using the coverage in 42 diploid NDBE samples from non-progressors (one sample from each individual) as a reference panel. Ten eigensamples generated from the reference panel by singular value decomposition were used for read-depth denoising of all samples.

***Identification and classification of SCNAs***

Due to the low sequencing coverage, we could not perform haplotype-specific copy-number calculation. We manually reviewed the copy-number data and found that many samples contained a low fraction of aneuploid cells (see NDBE sample in **Figure 8C** and examples in **Extended Data Fig. 9** and **10**). We expected that such events will likely be missed by standard copy-number segmentation algorithms and therefore manually reviewed the copy-number plot of each chromosome to identify the following SCNAs: (1) arm-level or whole-chromosome gain/loss, assessed from the genome-wide copy-number plots; (2) large segmental SCNAs (> 1Mb) that are shared by more than one sample; (3) recurrent focal deletions on 3p (near 60Mb, spanning *FHIT*) and 9p (near 21Mb, spanning *CDKN2A*); (4) complex SCNAs (including duplications/amplifications); (5) sloping copy-number variation. For complex SCNAs and sloping copy-number variation, we required at least part of the chromosome and most of the genome to have constant copy number to exclude false SCNA due to sequence coverage non-uniformity. We annotated SCNAs in each sample based on the evolution pattern (**Extended Data Table 8, Tab1**) and then generated a summary of SCNAs identified in all samples from each patient (**Extended Data Table 8, Tab2**); the latter was used to generate **Extended Data Fig. 9A**.

**Method References:**


m1. https://www.biorxiv.org/content/10.1101/861054v1 (Mutect2)

m2. https://www.nature.com/articles/s41586-020-2308-7 (gnomAD)

m3. https://onlinelibrary.wiley.com/doi/abs/10.1002/humu.22771 (Oncotator)

m4. https://nature.com/articles/s41568-018-0060-1 (COSMIC)

m5. https://www.biorxiv.org/content/10.1101/201178v3 (HaplotypeCaller)

m6. https://raw.githubusercontent.com/mskcc/ngs-filters/master/data/rmsk_mod.bed

m7. https://raw.githubusercontent.com/mskcc/ngs-filters/master/data/wgEncodeDacMapabilityConsensusExcludable.bed

m8. https://portal.firecloud.org/#methods/gatk/CNV_Somatic_Pair_Workflow/5 (GATK4 CNV)

m9. https://www.nature.com/articles/nbt.2203 (ABSOLUTE)

m10. https://www.nature.com/articles/ng.3643 (Sanger Imputation)

m11. https://www.nature.com/articles/nature15393 (1000 genomes project)

m12. https://www.nature.com/articles/nature14493 (Zhang et al., Nature 2015)

m13. https://genome.cshlp.org/content/28/4/581.full.html (SvABA for SV calling)

m14. https://data.broadinstitute.org/snowman/Submission/hg19.svaba.exclude.bed

m15. https://gatk.broadinstitute.org/hc/en-us/articles/360037054312-ASEReadCounter (ASEReadCounter)

m16. https://www.nature.com/articles/ng.3679 (EAGLE2)

m17. https://www.tandfonline.com/doi/full/10.4161/fly.19695 (snpEff)

**Supplementary Figure and Table Legends**

**Extended Data Figure 1**: Example histopathological images of Barrett's esophagus without dysplasia (NDBE), low-grade dysplasia (LGD), high-grade dysplasia (HGD), intramucosal esophageal adenocarcinoma (IMEAC), and esophageal adenocarcinoma (EAC).

**Extended Data Figure 2**: Phylogenetic trees with branch labels (copy-number alterations along each branch are listed in **Extended Data Table 2**) and annotated oncogene amplifications. Also shown are the pairwise genetic similarity matrix calculated from the fraction of shared sSNVs between each pair of samples (see **Methods**). The relative spatial locations of samples are shown next to the sSNV similarity heatmap. (This information is unavailable for Patient 2 and 5). The SCNA-derived phylogenetic trees are consistent with sSNV genetic similarity with the following exceptions: HGD from Patient 1; HGD/IMEAC1/IMEAC2 from Patient 12; all samples from Patient 15. The evidential support for the phylogenetic inference in Patient 12 and 15 is presented in **Supplementary Figure 6**. The HGD sample from patient 1 is a polyclonal mixture containing subclones of IMEAC cells; the grouping of HGD with cancer lesions is based on the shared *TP53* mutation and the presence of complex SCNAs in HGD that is absent in BE or LGD.

**Extended Data Figure 3**: Landscape of somatic copy-number alterations (SCNA) in BE and EAC lesions. **A**. Total SCNA burden (number of paternal and maternal autosomes with SCNAs) in each sample broken down by SCNA type and timing relative to whole-genome duplication (WGD). Local deletion/duplication, UPD, and arm-level SCNAs are colored the same as in **Figure 4**. Complex segmental copy-number changes are further subdivided to (1) terminal, paracentric, and pericentric gain/loss (orange); (2) complex alterations (red); and (3) focal amplifications (purple). Samples are grouped based on histopathological grading (non-dysplastic BE, low-grade dysplasia, high-grade dysplasia, and carcinoma). The mutation status of *TP53* is shown below each sample: black circles for bi-allelic inactivation, open circles for no identifiable alterations or mono-allelic inactivation. **B**. Mean SCNA burden of samples in each pathological group. **C**. Mean SCNA burden in samples with intact or inactive p53. The SCNA burdens in five HGD/EAC samples (four from Patient 9, one from Patient 1) without evidence of bi-allelic *TP53* inactivation are shown separately from NDBE/LGD samples without *TP53* inactivation. These five samples are excluded in the left plot or in **Figure 4A** (middle). **D**. Mean SCNA burden in samples without whole-genome duplication (WGD) or inferred to have occurred prior to or after WGD. **E**. Allelic distribution of SCNAs on each chromosome (Chr1-22, ChrX) in each patient. Chromosomes with SCNAs are represented by colored boxes or open boxes. (*Left*) Chromosomes with multiple SCNAs (at least three SCNA breakpoints) on at least one homolog; (*Right*) chromosomes with bi-allelic SCNAs. Open boxes represent chromosomes with one or multiple segmental SCNAs. If most SCNAs arise from independent alterations, they should more frequently affect both homologs than accrue on a single homolog. The predominance of mono-allelic SCNAs over bi-allelic SCNAs indicates a concentration of breakpoints on a single homolog that is consistent with catastrophic events or successive changes on a single unstable chromosome.

**Extended Data Figure 4**: SCNA classification based on the mutational mechanisms. I. Local sequence deletion/duplication and uniparental disomy (UPD) do not generate unstable (acentric/dicentric) chromosomes. These events comprise a majority of alterations seen in cells or clones with intact p53. We suggest UPDs arise from homology-dependent invasion of a broken chromatid into the intact homolog followed by a "half crossover" with an opposite moving replication fork, instead of conservative, unidirectional break-induced replication to the chromosome end. II-IV. Copy-number alterations resulting from single-generation chromosomal instability. II. Arm-level copy-number changes are generated by chromosome mis-segregation or abnormal mitosis, including cytokinesis failure and multipolar cell division. III. Terminal, paracentric, and pericentric segmental copy-number alterations can be generated by different types of breakage-fusion-bridge cycles (**Figure 5**). IV. Dicentric chromosome breakage or DNA damage in micronuclei can lead to chromosome fragmentation and create two-state (single-chromatid fragmentation) or three-state (sister-chromatid fragmentation) oscillating copy-number patterns. V and IV. Complex gains and focal amplifications resulting from multiple generations of chromosome breakage and asymmetric distribution. V. Chromothripsis or telomere loss can initiate breakage-fusion-bridge cycles with additional chromothripsis that generate complex copy-number gains (>3 state) or focal amplifications. VI. Focally amplified sequences can be either intrachromosomal (e.g., generated by intrachromosomal BFBs) or extrachromosomal (e.g., acentric fragments from chromothripsis).

**Extended Data Figure 5:** Copy-number outcomes of tetraploidization (**A**), micronucleation (**B**), single BFB cycle of a dicentric chromosome (**C**), and multigenerational evolution (including BFB cycles and chromothripsis) of dicentric chromosomes (**D**) determined from experimental analyses of chromosomal instability. **A**. Tetraploid (4N) cells can arise from failed cytokinesis or endoreplication (DNA replication without mitosis). The 4N cell has duplicated centrosomes that can both cause more chromosome segregation errors and generate highly aneuploid genomes after multipolar cell division. Multiple steps of this evolutionary process are suppressed by p53. **B**. Chromosome fragmentation in micronuclei results in chromothripsis with oscillating DNA copy number in both daughter cells. **C**. Resolution of dicentric bridge chromosomes results in reciprocal gain and loss of a telomere-bound segment (chromatid-type fusion) or a large internal segment (chromosome-type fusion, **Figure 5B**), with occasional DNA fragmentation near the breakage site leading to local chromothripsis (**Extended Data Fig. 8A-C**). **D**. Multigenerational evolution of a broken chromosome through breakage-fusion-bridge (BFB) cycles and chromothripsis. (*Top*) Copy-number outcomes of breakage-fusion-bridge cycles without chromosome fragmentation. (*Bottom*) Copy-number outcomes of breakage-fusion-bridge cycles after ancestral chromothripsis. The unstable chromosome generated by BFB cycles can become mitotically stable after acquiring a telomere through *de novo* telomere addition, translocation to a telomeric segment, or chromothripsis of a pair of sister chromatids. The broken chromosome can also give rise to an unstable ring chromosome (after loss of both telomeres) or acentric extrachromosomal circles (after centromeric loss), both of which can generate high-level gene amplifications.

**Extended Data Figure 6**: Additional evidence for the connection between dicentric chromosome breakage and terminal DNA loss and gain. **A**. A reciprocal translocation followed by whole-chromosome gain or loss can generate single-copy terminal loss (of the filled chromatid) or terminal gain (of the open chromatid). This model will generate an equal number of terminal losses and terminal gains (including segmental retentions). **B**. The disparity between terminal losses and terminal gains (only single-copy alterations) suggests that at least a subset of these alterations did not arise from reciprocal translocations. **C**. IGV snapshots of supporting discordant reads of two translocations shown in **Figure 6B**. **D.** A rearrangement junction between the breakpoint of the 10q-terminal gain and the telomeric breakpoint of the paracentric deletion on 12q. This junction cannot reflect a simple translocation as the translocated chromosome consisting of the 12q terminal segment and the 10q segment would be acentric and therefore not stably propagated.

**Extended Data Figure 7:** Gradual (sloping) copy-number variation on Chr.4p in the progeny population of a single cell with telomere loss engineered by CRISPR/Cas9 (Primary Clone 2b in Umbreit et al. (2020)). Shown are the bulk average DNA copy number (top) and the DNA copy number of subclones generated from single cells (bottom). The copy-number plots on the left and on the right are from different subclones. In each plot, gray dots represent total sequence coverage in 100kb intervals, red and blue lines represent segmented haplotype-specific DNA copy number of the parental homolog with progressive DNA losses. Only the parental homolog with continuous copy-number variation due to sequential BFB cycles is shown. Segmental copy-number alterations are shown below the DNA copy-number plots.

**Extended Data Figure 8:** Complex SCNAs from BFB cycles and chromothripsis. **A**. A schematic model of regional chromothripsis resulting from dicentric chromosome breakage. **B**. (*Top*) Chromothripsis near terminal deletion and duplication in HGD1. (*Bottom*) The haplotype phase in this region was determined based on allelic imbalance in IMEAC. **C**. Chromothripsis in a region of paracentric gain. The paracentric gain was likely generated in a chromosome-type BFB cycle that occurred downstream of the terminal loss generated in an earlier BFB cycle. **D**. Chromothripsis on 7q with breakpoints clustered near the boundaries of interspersed deletions. The clustering pattern of breakpoints is consistent with the "tandem-short-templates" signature reported in Umbreit et al. (2020). **E**. A schematic model of arm-level chromothripsis resulting from fragmentation of a dicentric chromosome in a micronucleus. **F**. Chromothripsis of a dicentric chromosome t(18p;17q) leading to three-state copy-number oscillation in both chromosome arms. **G**. Inter-chromosomal rearrangements and chromothripsis in Patient 7 with more than one rearrangement breakpoint near Chr8:73Mb, Chr9:4Mb and 15Mb, Chr12:89Mb, 117Mb, and 127Mb. **H** and **I.** DNA Amplifications spanning *ERBB2* generated by BFB cycles. (*Left*) DNA copy number of Chr17 shows ascending copy-number gains near *ERBB2* with 17q-terminal loss, which is consistent with BFB amplifications. The presence of interspersed deletions within the region of amplification in **I** indicates chromothripsis prior to amplification. (*Right*) Fluorescence in-situ hybridization (FISH) analysis shows amplified *ERBB2* in homogeneously staining regions (HSRs) next to Chr.17 centromeres as expected for intrachromosomal BFB amplification. **J**. (*Left*) Distinct focal amplifications spanning *MYC* on 8q in EAC and IMEAC of Patient 9. The shared boundary of amplified regions in both genomes suggests an ancestral chromosome breakage event with divergent downstream alterations. FISH analysis shows amplified *MYC* in both extrachromosomal (ecDNA) and intrachromosomal DNA. (*Right*) Amplified *FGFR2* is present as both extrachromosomal and intrachromosomal DNA. The localization of oscillating DNA deletion and amplification indicates regional chromothripsis preceding the amplification. The sharp copy-number transitions between deleted and amplified regions with few intermediate states are consistent with random segregation of intact double-minute chromosomes/ecDNA circles. All copy-number plots except in **B** and **C** only show the altered homolog. See **Downloadable Supplementary Data** for the complete plots of both homologs.

**Extended Data Figure 9:** BE copy-number evolution revealed from low-pass sequencing of longitudinal BE samples from Killcoyne et al. (2020). **A.** Association between BE progression and copy-number features in longitudinal BE or EAC samples from each patient, including (1) focal deletion/duplication (bottom), (2) arm-level gain/loss (second to bottom), (3) simple and complex segmental SCNAs (middle); and (4) aneuploidy (five or more chromosomes with either arm-level or segmental SCNAs) in longitudinal BE samples from non-progressors (left) and progressors (right). For each patient, the copy-number features are assessed from all longitudinal

BE/EAC samples, the number of which is shown above the copy-number feature heatmap and separated by pathological grading. The most distinguishing features are the presence of aneuploidy ($p = 10^{-10}$), complex SCNAs/focal amplifications ($p = 10^{-8}$) and segmental SCNAs ($p = 10^{-7}$). Sporadic arm-level alterations ($p = 0.05$) or focal deletion/duplication events ($p = 0.83$) show little difference between the two groups. All $p$-values are calculated by Fisher's exact test. **B.** Genome-wide DNA copy-number profiles of selected BE samples from Patient 72 and Patient 86 showing many private SCNAs (red arrows) indicating branching copy-number evolution.

**Extended Data Figure 10:** Evidence of chromosomal instability in BE cells from longitudinal BE sequencing **A**. Representative examples of large terminal (top) and internal (bottom) SCNAs in non-dysplastic BE samples. Notably, the left two examples were taken from a non-dysplastic BE sample from a non-progressor (Patient 45). Although we cannot rule out the possibility that this patient may eventually develop advanced EAC, the absence of EAC or high-grade BE at 60 months after this NDBE sample was taken indicates chromosomal instability being present at a very early stage of BE progression. Notably, this patient does not show 17p loss that is the most recurrent alteration in progressors (see **Extended Data Fig. 9B** for two examples) and contributes to p53 loss. The lack of complex segmental gains or copy-number heterogeneity also suggests limited chromosomal instability in the ancestor cell where the observed alterations first arose. **B**. Representative examples of chromosomes with both shared (solid lines) and private SCNA breakpoints (magenta lines) in longitudinal BE samples from three progressors. Note the sloping copy-number pattern in the HGD lesion (96 months) from Patient 88 in a region on Chr11q that displays clonal (discrete) copy-number changes in a BE with indetermined histopathology (IND) at 84 months.

**Extended Data Table 1**: Sample information and metrics: clinical information (Columns A-C), mean sequencing depth (Column D), inferred clonal fraction (Column E), ploidy (Column F), and WGD status (Column G) of the major aneuploid clone, basic metrics of sequencing data (columns H-L), allelic sequence coverage (columns M-R). Samples without identifiable SCNAs (> 0.1 Mb) are assigned as diploid with purity not assigned. Column M shows the number of heterozygous sites jointly detected from all samples (including both BE/EAC and normal reference). Column N shows the number of heterozygous sites with sequence coverage in each sample. Reference and alternate genotype coverage and standard deviation are calculated using heterozygous variant sites in each individual but excluding those with ≥100 allelic depth.

**Extended Data Table 2**: List of somatic copy-number alterations (SCNA) in individual samples (Tab 1) and individual evolutionary branches (Tab 2). In Tab 1, Patient_ID (Column A) and Sample_ID

(Column B) are according to **Extended Data Table 1**; Column C: unique ID for each SCNA; Columns D-F denote the altered chromosome and its parental haplotype; Column G: SCNA classification based on **Extended Data Figure 4** (1: focal deletion/duplication; 2: uniparental disomy; 3; arm-level SCNAs; 4: terminal SCNAs; 5: paracentric SCNAs; 6: pericentric SCNAs; 7: complex deletions/losses; 8: complex duplications/gains; 9: focal amplifications.); Column H: SCNA outcome (LOH: loss-of-heterozygosity through uniparental disomy; deletion: allelic DNA copy number = 0; loss: allelic DNA copy number < basal copy-number state; gain: allelic DNA copy number > basal copy-number state; amplification: allelic DNA copy number at or above 8); Column I: maximum allelic copy-number state; Column J: additional features of complex SCNAs. In Tab 2, Branch_IDs (Column A) are according to annotations in **Extended Data Figure 3**; Column B: number of progeny clones; Column C: relative timing to p53 loss (0 for prior to p53 loss, 1 for after p53 loss); Column D: relative timing to WGD (0 for pre-WGD branches, including all branches in patients without WGD; 1 for branches with WGD acquisition; 2 for post-WGD branches); Column E: number of SCNAs assigned to each branch, with individual SCNAs (annotated by unique IDs according to Column C of Tab 1) listed in Column F. Tab 3 contains a summary of the total number of SCNAs of each class grouped by their relative timing to WGD. Tab 4 contains a summary of the number of SCNAs of each class along different evolutionary branches grouped by their relative timing to WGD.

**Extended Data Table 3**: Summary of genetic alterations in single BE cells related to Figure 3. Tab 1: *TP53* mutation status. Tab 2-7 list single cells from different clones/groups and their shared or private SCNAs. Aneuploid cells (Tab 7) are separated into two groups (near diploid and near tetraploid) with allelic copy-number states of each homolog annotated individually. Alterations shared by support more recent See **Downloadable Supplementary Data** for detailed copy-number plots of every cell.

**Extended Data Table 4**: List of focal SCNAs (<1Mb) (Tab 1) and uniparental disomy (Tab 2) alterations detected in bulk BE and EAC samples. Each row represents a SCNA that is annotated for the region of alteration (Column B), the altered haplotype according to copy-number plots in **Downloadable Supplementary Data** (Column C), copy number outcome (Column D), affected samples (Column E), phylogenetic pattern (Column F), relative timing to *TP53* inactivation (Column G), and relative timing to WGD (Column H). We further added events inferred to have occurred prior to UPDs in column I of Tab 2.

**Extended Data Table 5**: List of SCNAs consistent with the outcomes of single (Tab 1) or multigenerational (Tab 2) BFB cycles. For single-generation BFB outcomes (Tab 1), Column D shows the copy-number classification; for multigenerational outcomes (Tabs 2), Column D shows the inferred evolutionary sequence. Columns E-G are similar to **Extended Data Table 4** except for the omission of relative timing to *TP53* inactivation as most of these events were inferred to have occurred after *TP53* inactivation. Additional remarks are put in Column H.

**Extended Data Table 6**: List of SCNAs indicating branching outcomes of BFB cycles. Tab 1: Progressive DNA losses identified in different BE/EAC genomes. Tab 2: Distinct SCNAs on the same parental homolog identified in different BE/EAC genomes from the same patient. Tab 3: SCNAs corresponding to complementary DNA retention and deletion. Annotations are similar to **Extended Data Table 5**, except for the addition of the span of SCNA in Column D of Tab 2 and Tab 3.

**Extended Data Table 7:** List of chromothripsis (Tab 1) and focal amplifications (Tab 2) detected in bulk BE/EAC samples, with results of fluorescence in-situ hybridization analyses listed in Tab 3. In Tab 1, the annotations are similar to **Extended Data Table 5**, with additional annotations of the span of chromothripsis (Column G), status of telomere loss (Column H), chromothripsis subclassification (Column I), and copy-number feature (Column J). The evolutionary sequence (column K), including preceding (Column L) and downstream (Column M) alterations relative to chromothripsis, is inferred based on the copy-number features. The mechanistic inference related to bridge resolution (Column N) and micronucleation (Column O) is made based on the span and copy-number feature of chromothripsis. In Tab 2, additional annotations include potential oncogenes (Column I), EAC oncogene (Column J), and co-localization with GISTIC peak regions (Column K) of each amplified region.

**Extended Data Table 8**: Re-analysis of longitudinal BE/EAC sequencing data. Tab 1: List of ancestral (Column F), shared (Column G), and private (Column H) SCNAs identified in each sample. Risk classification from the original study is kept in Columns I-K. Tab 2: Patient-level summary of SCNAs detected in all longitudinal samples from each individual. Focal/segmental losses/gains are annotated as 1p- or 1p+, bi-allelic/overlapping deletions are annotated as 9p--, gain or loss of entire chromosome arms are annotated as +1p, +8q, etc.
